# FREDDIE: A comprehensive tool for detecting exonization of retrotransposable elements in short and long RNA sequencing data

**DOI:** 10.1101/2024.04.22.590610

**Authors:** Rafael L. V. Mercuri, Thiago L. A. Miller, Filipe F. dos Santos, Matheus F. de Lima, Aline Rangel-Pozzo, Pedro A. F. Galante

## Abstract

**Background:** Transposable elements (TEs) constitute a significant portion of mammalian genomes, accounting for about 50% of the total DNA. Intragenic TEs are of particular interest as they are co-transcribed with their host genes in pre-mRNA, potentially leading to the formation of novel chimeric transcripts and the exonization of TEs. The abundance of RNA sequencing data currently available offers a unique opportunity to explore transcriptomic variations. However, a significant limitation is the capability of existing computational tools. Here, we introduce FREDDIE, an innovative algorithm designed to detect the exonization of retrotransposable elements using RNA-seq data. FREDDIE can process short and long RNA sequencing data, assemble and quantify transcripts, evaluate coding potential, and identify protein domains in chimeric transcripts involving exonized TEs and retrocopies.

**Results:** To demonstrate the efficacy of FREDDIE, we analyzed and validated TE exonization in two human cancer cell lines, K562 and U251. We have identified 322 chimeric transcripts, of which 126 were from K562, and 196 were from U251. Among these chimeric transcripts, there were 35 that showed similar exonization patterns and host genes. These transcripts involve protein-coding genes of the host and exonization of LINE-1 (L1), Alu elements, and retrocopies of coding genes. We have selected some candidates and validated them experimentally through RT-PCR. The validation rate for these candidates was 70%, later confirmed by long-read sequencing.

Additionally, we applied FREDDIE to analyze TE exonization across 157 glioblastoma samples, identifying 1,010 chimeric transcripts. The majority of these transcripts involved the exonization of Alu elements (69.8%), followed by L1 (20.6%) and retrocopies (9.6%). Notably, we discovered a highly expressed L1 exonization within the ROS gene, resulting in a truncated open reading frame (ORF) with the deletion of two protein domains.

**Conclusions:** FREDDIE is an efficient and user-friendly tool for identifying chimeric transcripts that involve exonization of intragenic TEs. Overall, FREDDIE enables comprehensive investigations into the contributions of TEs to transcriptome evolution, variation, and disease-associated abnormalities, and it operates effectively on standard computing systems.

FREDDIE is publicly available: https://github.com/galantelab/freddie

## BACKGROUND

Transposable elements (TEs) are ubiquitously found across all animal genomes and represent a significant portion of mammalian DNA. While DNA transposons remain largely inactive in most mammals, retrotransposons—here referred to as TEs—have played a dynamic role in mammalian evolution [1]. Consequently, some species, including humans and other primates, have genomes in which more than half of the content consists of TEs [2]. Particularly notable are the LINE-1 (L1) and Alu elements in primates, which have been instrumental in driving genomic diversity and complexity across several species. These elements contribute to genetic variation through mechanisms such as insertions, deletions, and recombination events, thereby influencing gene expression and regulatory networks. However, their mobility also introduces risks, potentially leading to genomic instability and disease [3].

Numerous studies and tools have been developed to explore the activities of TEs in the contexts of evolution, health, and disease [4,5]. Yet, fixed TEs, which have become stable components of the host genome and lost their transpositional capability due to mutations, deletions, or epigenetic modifications, have received comparatively less attention. Notably, a particular class of fixed TEs, those inserted into intronic regions of protein-coding genes (intragenic TEs), has emerged as a subject of interest. Although most multi-exonic mammalian protein-coding genes contain at least one intragenic TE, these are predominantly spliced out during post-transcriptional processing of pre-mRNA [6]. Nevertheless, the exonization of TEs, particularly of Alu elements [7], has yet to be explored despite its occurrence in healthy and cancerous tissues [3,8].

The advent and evolution of RNA sequencing (RNA-seq) methodologies have revolutionized the study of transcriptomes and their variations [9]. Before the advent of RNA-seq, techniques were constrained by their reliance on pre-existing knowledge of gene sequences and variations or by their low throughput. In contrast, RNA-seq offers a high-throughput, comprehensive, and unbiased approach, enabling detailed exploration of gene expression, the identification of previously uncharacterized gene isoforms [10,11], and even the study of processed pseudogenes [12]. However, the full potential of RNA-seq is often limited by the capabilities of computational algorithms to analyze the generated data, presenting significant challenges in fully assessing and interpreting the complexities of novel genes and isoforms uncovered by this methodology.

Here, we introduce FREDDIE, an innovative algorithm to identify exonization events involving intragenic TEs within mammalian genomes. FREDDIE adeptly processes RNA sequencing data to assemble, quantify, and assess the coding potential of chimeric transcripts resulting from such molecular phenomena. The FREDDIE algorithm was applied to two human cancer cell lines, K562 and U251, and to glioblastoma multiforme (GBM) samples to demonstrate its utility. Our findings suggest that FREDDIE is a powerful tool for exploring the roles of TEs in transcriptome evolution, variability, and disease. These results underscore its potential for broader applications in genomic research.

## MATERIAL AND METHODS

### RNA sequencing data

We obtained RNA sequencing data from two human cell lines: K562, lymphoblast cells isolated from a patient with chronic myelogenous leukemia, and U251, cells derived from a malignant glioblastoma tumor. Specifically, for the K562 cell line, we utilized both short-read (76 bp) and long-read (>1000 bp) sequencing data. The short-read data were obtained from polyA+ libraries, with the sequence data available under accession numbers SRR315336 and SRR315337. The long-read sequencing data were accessed under accession numbers SRR14638109, SRR14638110, SRR10838645, and SRR10838646. In the case of the U251 cell line, we analyzed short-read sequencing data from three replicates, available under the accession number GSE141013. Additionally, we incorporated short-read sequencing data from 172 glioblastoma samples available through The Cancer Genome Atlas (TCGA), accessed via the portal at https://portal.gdc.cancer.gov/.

### Intragenic identification of TEs and retrocopies in protein-coding genes

For our analysis across multiple species, including humans, we extracted the genomic coordinates for various TEs — such as LINE-1, SINEs, SVAs, and LTRs—from the Dfam database [13]. Locations of retrocopies were sourced from RCPedia [14,15]. We employed GENCODE (version 36) [16] for comprehensive genomic annotation of all protein-coding genes in the human genome.

Custom in-house scripts were developed to intersect the genomic coordinates of TEs and retrocopies with those of known genes, specifically targeting intragenic TEs and retrocopies. This method was similarly applied to other species, including chimpanzee (panTro6), cow (bosTau9), dog (ROS_Cfam_1.0), marmoset (calJac4), mouse (mm39), opossum (monDom5), platypus (mOrnAna1.p.v1), rat (rn6), and rhesus (rheMac10). For these species, we adapted our approach to include species-specific retrocopies from RCPedia and transcriptome references from RefSeq. The datasets generated from this analysis are available for further use in our public repository, accessible at https://bioinfohsl-tools.s3.amazonaws.com/freddie/databases/

### STAR read alignment step

To utilize FREDDIE, FASTQ files must first be aligned to a reference genome. This critical step is carried out using the STAR aligner [17], which is integrated into FREDDIE’s pipeline. The alignment process accommodates various types of sequencing data, including paired-end, single-end, and long-reads. Detailed documentation on this integration is available at https://github.com/galantelab/freddie.

Following the alignment, the pipeline is meticulously configured only to select uniquely mapped reads. These reads are characterized by an alignment score of Q=255 in the STAR output, ensuring the accuracy of subsequent analyses. This selection process is fully automated within the FREDDIE pipeline, requiring no manual intervention from the user.

Reference files necessary for the STAR alignment, including genome and transcriptome annotations, can be downloaded from our dedicated repository at https://bioinfohsl-tools.s3.amazonaws.com/freddie/databases/. Alternatively, users can generate these files independently, following the guidelines provided in the STAR documentation at https://github.com/alexdobin/STAR/.

### Transcriptome assembly and selection of TEs under exonization with FREDDIE

FREDDIE integrates StringTie2 [18] for transcriptome assembly, streamlining the data processing within its pipeline. Once the assembly is complete, FREDDIE utilizes bedtools [19] to perform a detailed comparative analysis of the assembled transcripts against the genomic locations of intragenic TEs and retrocopies. This comparison allows FREDDIE to identify and select chimeric transcripts that demonstrate exonization of TEs or retrocopies intertwined with exonic sequences from their host genes. This functionality highlights the complex interactions between TEs and genomic architecture.

During this pipeline phase, FREDDIE offers users the flexibility to adjust the stringency of event identification based on their research needs. Users can choose the “reciprocal” mode, where an event is recognized only if the TE constitutes a minimum of 50% of the exon’s length and the exon comprises at least 50% of the transcript. Alternatively, the “not reciprocal” mode, which is the default setting, applies a less stringent criterion: it requires only that at least 50% of the TE overlaps with the exon.

### Classification of coding chimeras and domain analysis with FREDDIE

FREDDIE assesses chimeric transcripts for the presence of a reliable and putatively functional Open Reading Frame (ORF). This process begins with the use of RNASamba [20], a neural network-based tool designed to evaluate sequences with coding potential. FREDDIE applies RNASamba to identify chimeric transcripts that exhibit TE exonization and demonstrate a high probability (greater than 90%) of containing a real ORF.

For those chimeric transcripts deemed to be coding, FREDDIE proceeds to extract their protein sequences and employs HMMER [21]—alongside the Pfam database [22] —to conduct a thorough search for protein domains. This search aids in identifying and quantifying the protein domains present within the transcripts, including those from the host gene.

Additionally, FREDDIE’s criterion for recognizing novel transcripts involves detecting significant alterations in domain composition or instances where a domain exhibits a substantial reduction in length - specifically, reductions exceeding 50% compared to the corresponding domain in the host gene.

### Expression profile

FREDDIE also incorporates functionality for transcript expression quantification. To achieve this, FREDDIE utilizes StringTie2, which is adept at handling both short and long-read data. Notably, beyond using the standard reference transcriptome (e.g., GENCODE), FREDDIE integrates the chimeric transcripts identified during the initial assembly step. This integration enables comprehensive quantification, encompassing all chimeric transcripts generated earlier in the processing pipeline.

### Reverse transcription-PCR and gene expression analysis

To synthesize the first strand of cDNA, we used one μg of total RNA from U251 and K562 cell lines and employed the LunaScript® RT SuperMix Kit for reverse transcription. Primers were meticulously designed to target novel exon junctions, facilitating the detection of chimeric transcripts (refer to Table S1 for primer details). Following reverse transcription, PCR amplification was performed, and the resultant PCR fragments were analyzed on a 1% agarose gel, followed by sequencing using the Sanger method. For quantitative analysis, quantitative polymerase chain reaction (qPCR) was conducted using the LunaScript® Multiplex One-Step RT-PCR Kit (New England Biolabs Inc., MA, USA), strictly following the manufacturer’s instructions. The human hemoglobin subunit beta (HBB) gene was the reference for Ct normalization. Relative quantification of gene expression was calculated using the 2^(-ΔCt) method, where ΔCt represents the difference in threshold cycle values between the gene of interest and the HBB gene (ΔCt = Ct_Gene_-Ct_HBB_).

## RESULTS

### Implementation and usage of FREDDIE

FREDDIE’s execution is designed to be straightforward. Users are required only to provide their RNA-seq data in FASTQ format and follow a series of steps to generate chimeric transcripts that exhibit exonization of TEs. Figure 1 illustrates the entire workflow for operating FREDDIE.

**Figure 1.**
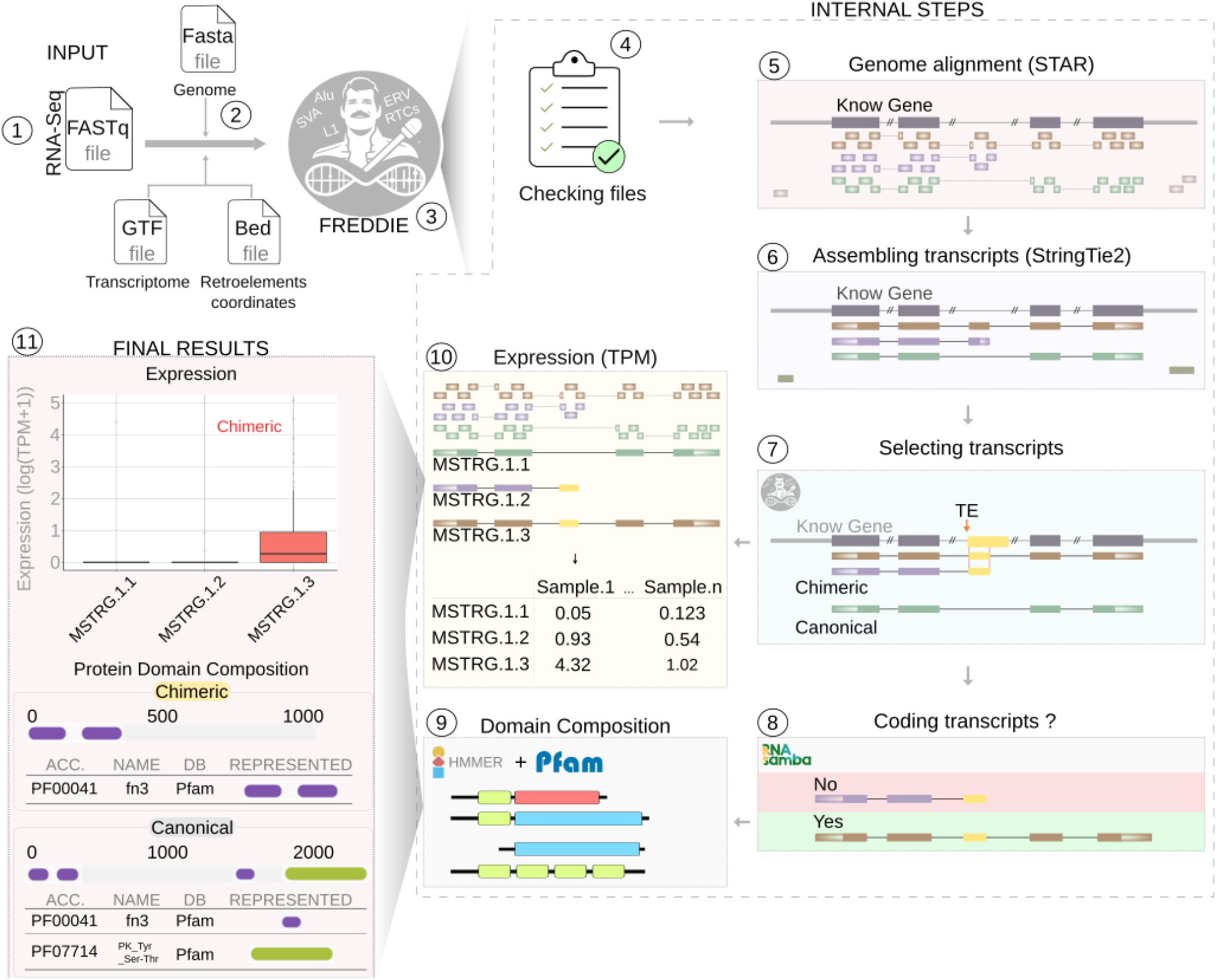
The step-by-step process of the FREDDIE pipeline, from input to final results. 1. Input: Users input RNA-seq data in FASTQ format along with the FASTA genome file, GTF file of the transcriptome, and BED file of retroelement coordinates. 2.Reference Retrieval: FREDDIE acquires the necessary reference genome, transcriptome, and retroelement coordinates. 3. Execution: The user initiates the FREDDIE pipeline. 4. File Checking: FREDDIE verifies the completeness and integrity of the input files. 5. Genome Alignment: The STAR aligner within FREDDIE aligns reads to the reference genome. 6. Transcript Assembly: StringTie2 is used to assemble transcripts based on the reference genes. 7. Transcript Selection: FREDDIE selects chimeric transcripts containing exonized TEs (termed ‘Chimeric’) and other transcripts (termed ‘Canonical’). 8. Coding Potential Analysis: RNASamba assesses the coding potential of the chimeric transcripts. 9. Domain Composition: HMMER and Pfam analyze the protein domains of the coding transcripts. 10. Expression Quantification: Expression levels, measured in TPM (Transcripts Per Million), are quantified for all transcripts using StringTie. 11. Final Results: The final output includes expression levels of chimeric transcripts and a comparison of protein domain compositions between chimeric and canonical transcripts. This comprehensive visualization captures FREDDIE’s functionality from raw data input to producing actionable biological insights.

Initially, users submit their RNA-seq data (step #1). Subsequently, the necessary reference genome, reference transcriptome, and retroelement coordinates for all species are obtained (step #2); these reference datasets are available in our repository (refer to the documentation for details). Once the setup is complete, users can execute FREDDIE (step #3) and await the generation of results (as represented in step #11).

In the interim, FREDDIE conducts several processes to achieve the final results. It begins by verifying the completeness of all input data (step #4). It then proceeds to align reads against the reference genome using the STAR aligner integrated within FREDDIE (step #5). Following alignment, FREDDIE performs transcript assembly using the reference genes via StringTie, also integrated within the tool (step #6). It then selects transcripts that contain exonization of TEs (termed ‘Chimeric Transcripts’) and those that do not (termed ‘Canonical Transcripts’) using custom scripts (step #7). The chimeric transcripts are further assessed for their coding potential using RNASamba (step #8), ensuring a reliable set of coding transcripts is identified. These coding transcripts are then evaluated for their protein domains using HMMER and the Pfam database (step #9). Simultaneously, all transcripts undergo expression quantification via StringTie (step #10). Finally, FREDDIE generates comprehensive results detailing chimeric and canonical transcripts’ expression and protein domain composition. (step #11).

### Chimeric transcripts in cell lines

Utilizing FREDDIE, we examined the exonization of TEs in K562 and U251 cell lines, detecting 322 chimeric transcripts. In K562, we identified 126 chimeric transcripts (comprising 10 retrocopies, 30 LINE1, and 86 Alu elements), while in U251, there were 196 (containing 10 retrocopies, 39 LINE1, and 147 Alu elements), detailed in Table S2. To assess the accuracy of FREDDIE’s detections, we conducted an *in silico* validation using long-read sequencing data for K562. Selection criteria for genes included having chimeric transcripts with a minimum of 10 reads covering isoforms and exact matches in exonization and exon junctions for validity. This validation confirmed approximately 70% of the identified transcripts, with 70% having all exons accurately identified (Figure S1, Table S3).

Regarding retrocopies, most chimeric transcripts (55%) featured the event in the last exon. Moreover, 70% of these transcripts were predicted as coding (with a probability of ≥90%), implying that these retrocopies could be exapted, potentially affecting the translation of the host transcript (Figure 2A). Additionally, domain alterations within host genes were maintained or deleted (in whole or partially) in about 86% of cases (Figure 2B, Table S4).

**Figure 2:**
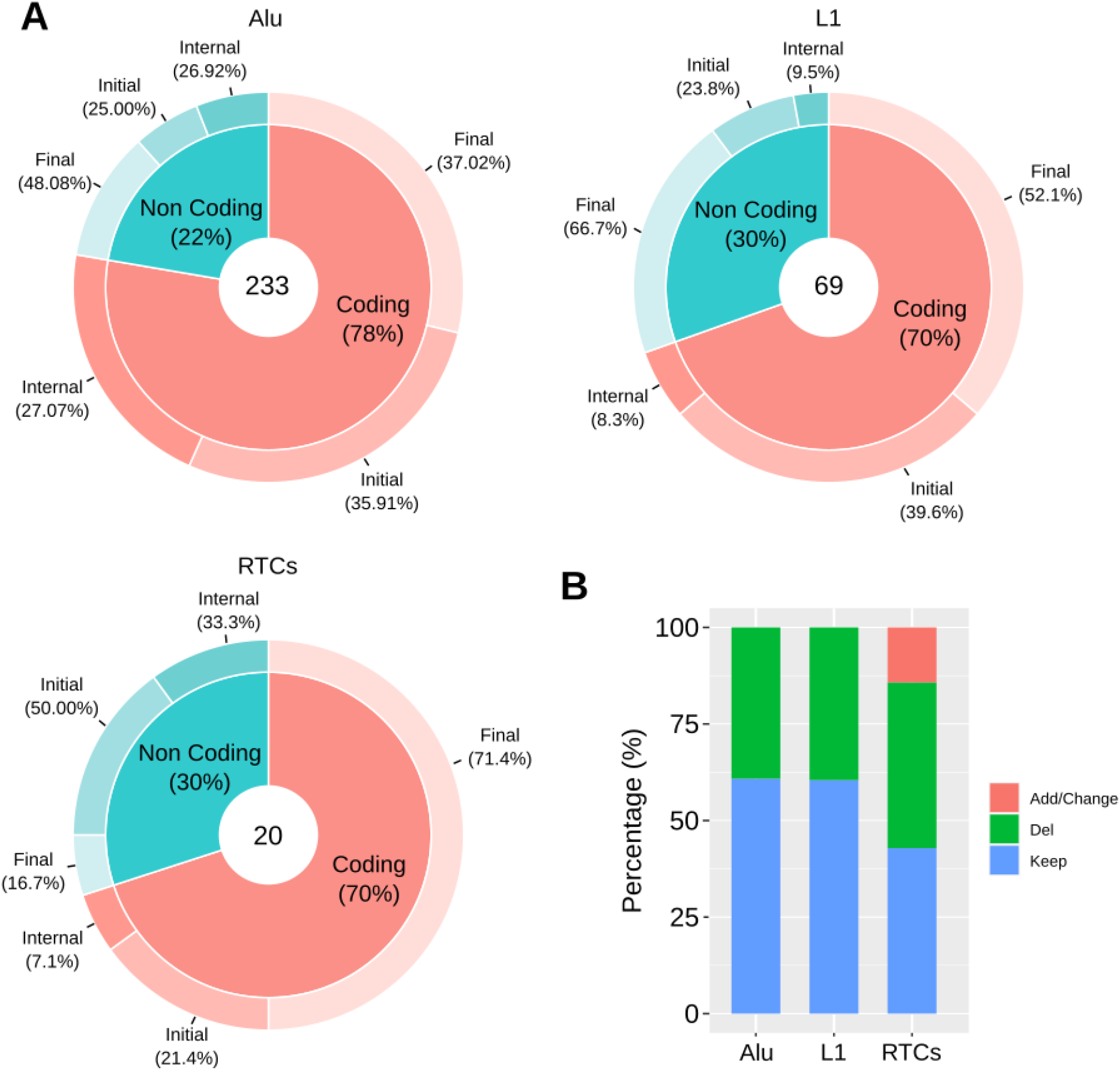
Retroelement exonization and domain alteration profiles in chimeric transcripts. A) Coding potential and exonization location analysis. Donut charts represent exonization distribution in Alu, L1, and retrocopies within chimeric transcripts from cell lines K562 and U251. The charts are divided into ‘Coding’ and ‘Non-Coding’ segments, with the percentage of each category indicated. The location of the exonization events—’Initial’, ‘Internal’, and ‘Final’—is annotated within each segment, providing insight into the positional preference of exonization events within transcripts. B) Domain alteration analysis. A stacked barplot illustrates the frequency of domain alterations in chimeric transcripts involving Alu, L1, and retrocopy elements. Each bar represents 100% of the transcripts for the element type, segmented by the type of alteration: domain addition or change (green), domain deletion (red), and complete domain maintenance (blue). The plot offers a comparative overview of exonization’s structural impact on protein domains across different retroelement types.

Distinct patterns were observed for LINE1 and Alu element chimeric transcripts. Specifically, 56.5% of LINE1 exonization events were located in the last exon, a trend resembling that of retrocopies. Alu element exonizations, however, predominantly occurred in the last exon (39.48%) but also significantly in the first exon (33.47%), with a notable majority on the opposite strand (74.35% in first exon transcripts), supporting existing literature on such orientation-specific prevalence [7,23].

Similar percentages of coding RNAs were noted for both LINE1 (69.57%) and Alu (77.68%). The impact of exonization on domain alteration is less detrimental for LINE1 and Alu, as 60.41% and 60.77%, respectively, maintained complete domain integrity (Figure 2, Table S3).

### Expression analysis of chimeric transcripts

We quantified the expression of chimeric transcripts utilizing the StringTie expression module from FREDDIE, which proved effective in delineating the transcriptional profiles of our identified transcripts. The expression levels of chimeric transcripts were clear, often representing more than 50% of the corresponding host gene expression levels in the short-read RNA-seq data from the K562 cell line, Figure 2.This relative expression indicates that the chimeric transcripts are not merely transcriptional noise but may play a substantial role in the cellular transcriptome. Figure 3 and Table S5 provide a comprehensive visual and tabular summary of these expression analyses.

**Figure 3:**
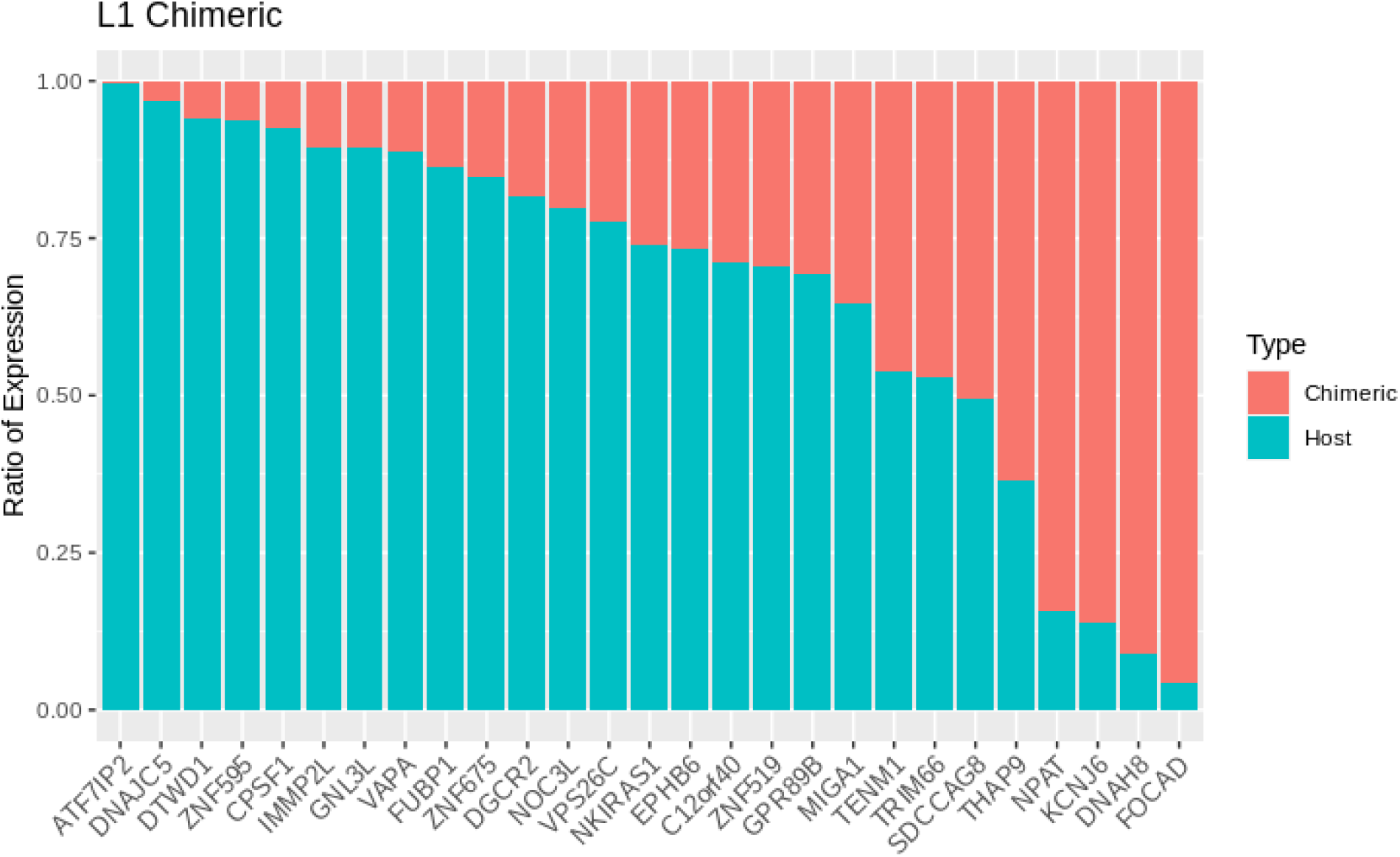
Comparative expression analysis of L1 chimeric transcripts. The bar graph displays the expression ratio between chimeric (turquoise) and host (salmon) transcripts for LINE-1 elements across various genes identified in the K562 cell line. The vertical axis represents the expression ratio, and the horizontal axis lists the genes with detected chimeric transcripts. This result shows chimeric transcript expression’s prevalence and potential significance relative to their host genes.

### RT-PCR and long-read validation of novel chimeric transcripts

To substantiate the computational predictions made by FREDDIE, we implemented a dual validation strategy using long-read data and RT-PCR validations. Initially, we utilized reverse transcription-PCR (RT-PCR), followed by Sanger sequencing and quantitative PCR (qPCR), designing primers that target novel exon junctions. This approach enabled the experimental confirmation of newly identified chimeric transcripts. We selected two representative transcripts, EFHC2-RRM2 and LRRFIP2-UBE2F, based on their characterization and expression profiles from our dataset (Table S2). The experimental procedures confirmed the co-optation of the retrocopy by the host gene, as visualized in Figures 4A and 4B. The quantification of expression differences between the transcripts utilized average Ct values. In U251 cells, the transcripts referred to as Can 2 and U1A exhibited delta Ct fold differences of 15.63 and 7.05, respectively.

**Figure 4.**
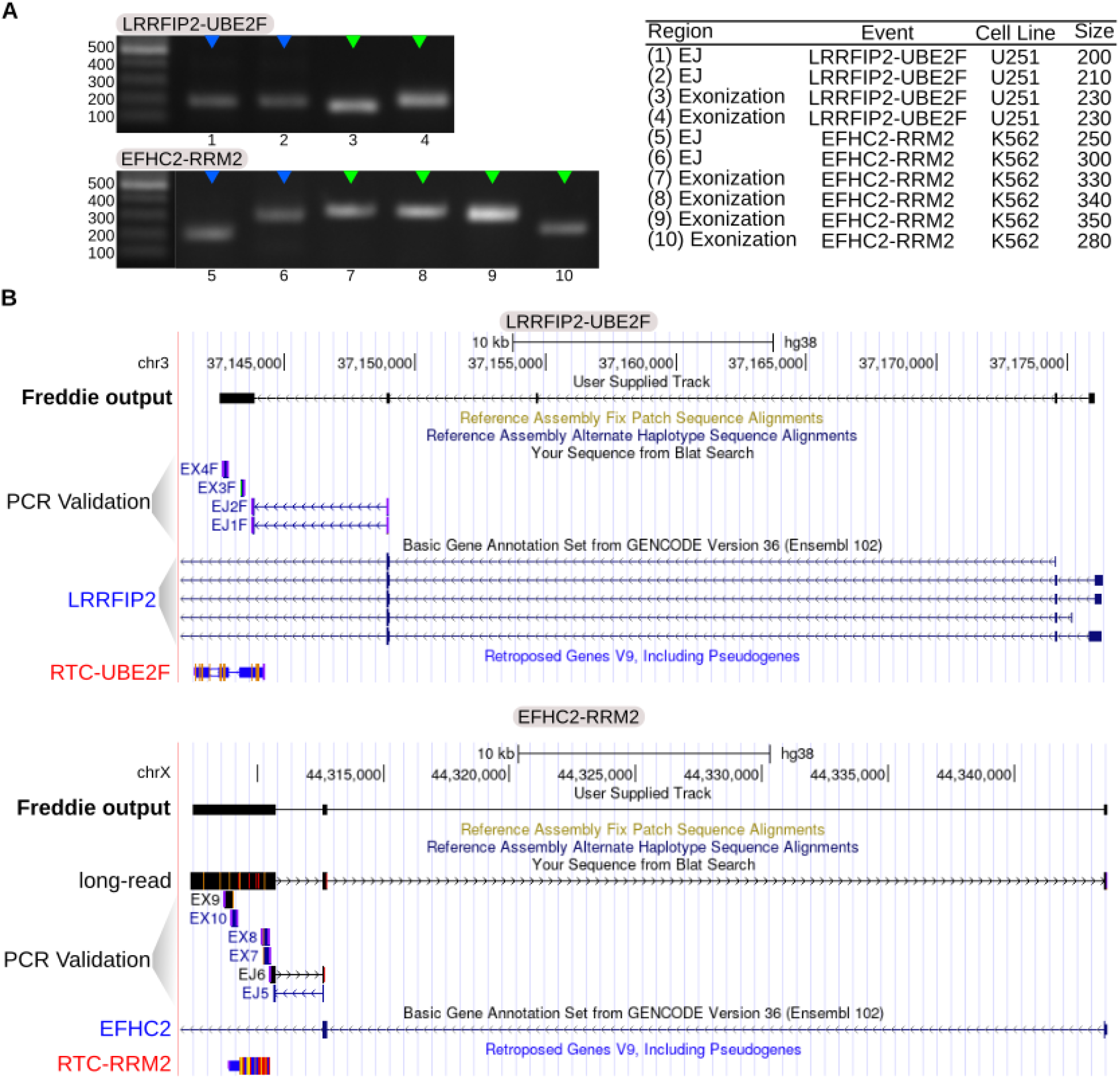
PCR validations of chimeric transcripts between retrocopies and their host genes. A) RT-PCR results for selected chimeric transcripts from the K562 and U251 cell lines. The gel images show the amplified fragments, with blue and green arrows indicating the specific bands corresponding to retrocopies and host gene exon junctions. The adjacent table enumerates the bands, detailing their regions, events, cell lines, and size in base pairs. B) Genomic alignment visualization of the sequences obtained from RT-PCR against the human reference genome. These alignments demonstrate the genomic integration points of chimeric transcripts for LRRFIP2, EFHC2 (host genes) and UBE2F, RRM2 (representing retrocopies), respectively. Annotations include exon-intron structures, with exonic regions of the chimeric transcripts highlighted to correlate with the RT-PCR findings. The tracks display both the FREDDIE output and PCR validation alignments to provide a comprehensive view of the transcript structure in the genomic context.

Conversely, in the K562 cell line, transcript P1 did not demonstrate a significant difference, with a delta Ct fold change of only 1.27. Moreover, we also searched for reads from a long-read dataset of the K562 cell line, seeking overlaps with at least one exon junction identified in the chimeric transcripts assembled by our pipeline, Figure 4B. Altogether, Figure 4 comprehensively demonstrates the experimental validation of chimeric transcripts identified by FREDDIE. It clearly illustrates the precise exonization events and offers solid empirical evidence to support the reliability of the computational predictions made by the algorithm.

### Case Study in GBM Samples: L1 and HERVL Exonization in ROS1

In an analysis of 157 TCGA GBM samples, we identified 1,010 chimeric transcripts linked to fixed TE events (refer to Table S6). Alu elements were notably involved in chimeric transcripts at the first exon (316 out of 706 cases) and were frequently oriented opposite to the host gene (547 out of 706). In contrast, LINE1 elements and retrocopies typically featured in the last exon (96 out of 209 for LINE1 and 55 out of 97 for retrocopies) and aligned with the orientation of the host gene.

We turned our attention to the tumor suppressor gene ROS1, as classified by COSMIC v94, to examine the effects of exonization events. Notably, two variants of chimeric transcripts were observed: ROS1 with L1 integration (MSTRG.35072.8) and ROS1 with concurrent L1 and HERVL integration (MSTRG.35072.7), as displayed in Figure 5A. Both variants were subjected to an exhaustive analysis using the FREDDIE pipeline. The analysis revealed that the exonization events led to premature truncation of transcription, resulting in a coding sequence devoid of two domains typically present in the canonical ROS1 gene (Figure 5B).

**Figure 5:**
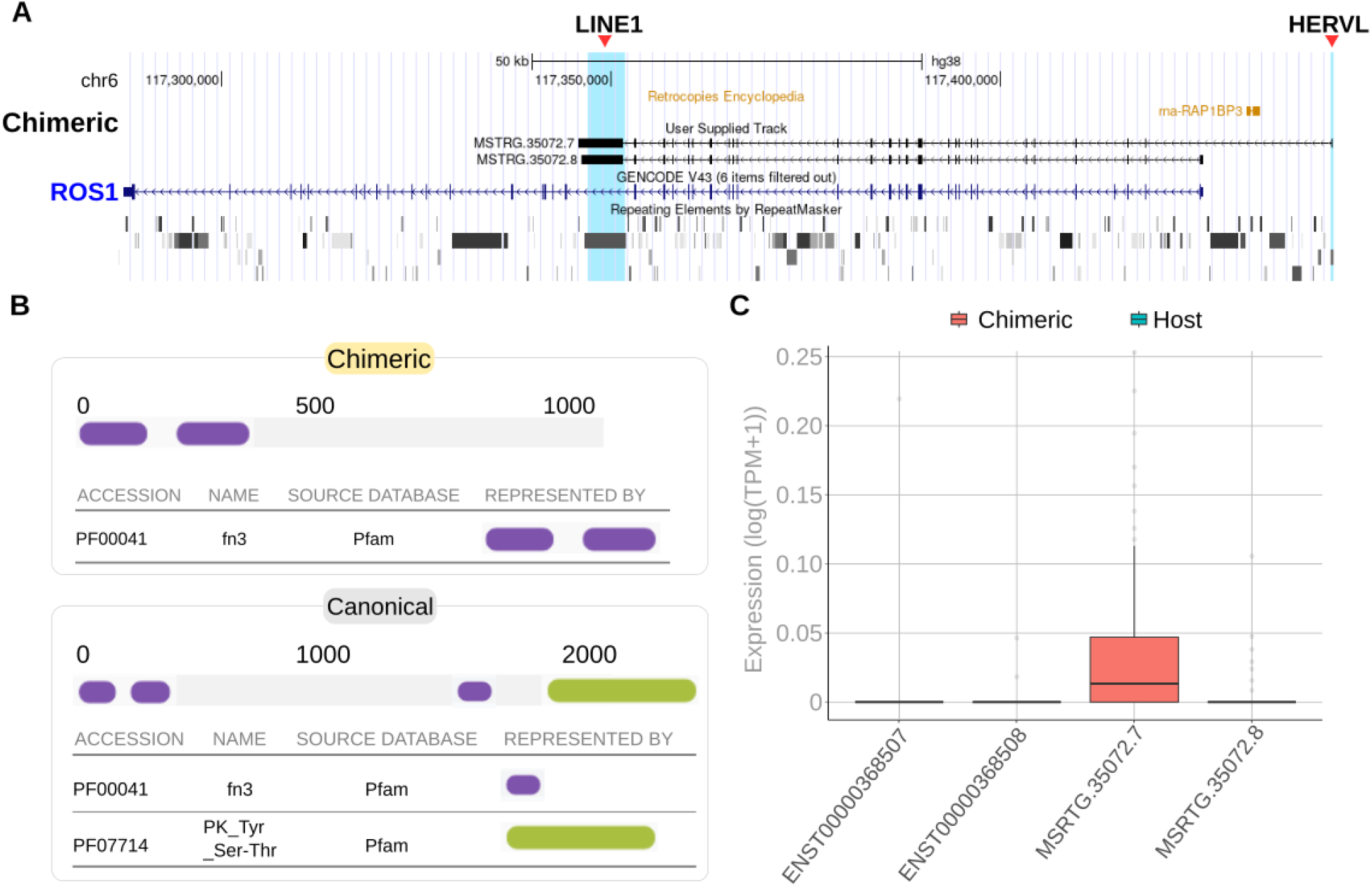
Analysis of L1 and HERVL exonization in ROS1 and resultant chimeric transcripts. A) Genomic context of L1 and HERVL exonization in ROS1. MSTRG.35072.7 and MSTRG.35072.8 are the chimeric transcripts identified. The red arrow indicates the L1 or HERVL integration. B) Protein domain profiles from chimeric and canonical transcripts. Comparative protein domain analyses depict the differences between the chimeric transcript (bearing the L1 exonization) and the canonical ROS1 transcript. The domain compositions, sourced from Pfam, are illustrated, and missing domains in the chimeric transcript are highlighted. C) Expression levels of chimeric versus host transcripts. The box plot shows the expression levels, measured in transcripts per million (TPM), of the chimeric transcripts (red) and those of the host ROS1 gene.

Expression analysis indicated that one chimeric transcript (MSTRG.35072.7) was particularly abundant, with its expression statistically surpassing that of the host gene’s other isoforms, suggesting significant transcriptional impact (Figure 5C). This transcript (MSTRG.35072.7) also highlighted the role of HERVL in initiating transcription, reflecting the intricate dynamics of TEs in gene expression regulation.

Overall, these findings elucidate the consequences of TE and retrocopy exonization on the architecture and expression profiles of protein-coding genes, with potential repercussions for the genes’ functionality and protein synthesis.

## DISCUSSION

Reverse-transcribed elements like TEs and retrocopies are predominantly located in the intronic regions of protein-coding genes [1,12]. Identifying chimeric transcripts between these elements and their host genes is crucial for understanding health and disease contexts [24]. To facilitate and improve this, we introduced FREDDIE, a novel algorithm designed to detect the exonization of retrotransposable elements and retrocopies using short or long RNA-seq data. FREDDIE can process RNA-seq data, assemble and quantify transcripts, evaluate coding potential, and identify protein domains in chimeric transcripts with exonized TEs. FREDDIE is publicly available (https://github.com/galantelab/freddie), straightforward to use, well-documented, and comes with pre-processed data for multiple species, including humans and mice.

Previous studies have explored retroelements as sources of new chimeric transcripts in various contexts [23,25–27]. In terms of algorithms and pipelines, most strategies target specific retroelements (e.g., LINE1, Alu, and retrocopy) without exploring other characteristics like coding properties, domain changes, or expression levels [28–30]. FREDDIE, however, provides a comprehensive solution, allowing the detection of any fixed event and exploration of diverse genomic conditions as the user desires.

Our results demonstrated FREDDIE’s capability to unveil a considerable number of chimeric transcripts involving TEs and retrocopies. The identification of 1,010 chimeric transcripts in GBM indicates the significant prevalence and diversity of exonization phenomena within the human genome, particularly in cancer. Additionally, FREDDIE’s results shed light on critical features of TEs, such as orientation and positional preferences, suggesting a complex regulatory mechanism affecting transcript diversity. These findings contribute to our understanding of how TEs and retrocopies can alter transcriptomic landscapes in cancer genomes, reinforcing earlier studies linking TE integration with gene regulation, genomic stability, and evolutionary processes.

The case study focusing on ROS1 exemplifies the potential functional impact of these exonization events. The detection of chimeric transcripts with truncated domains in a known tumor suppressor gene emphasizes the possible pathogenic implications of TEs in cancer (GBM).

The presence of chimeric transcripts with high levels of expression, exceeding those of normal host gene versions, can have a significant impact on gene function and phenotype. This is particularly important in the case of GBM, where disruptions in gene expression play a key role in the development of tumors [11,31–33]. Additionally, the co-occurrence of HERVL indicates that various TEs may interact with each other, potentially leading to synergistic or competing effects on gene expression.

Despite its utility, FREDDIE has certain limitations, notably its reliance on pre-processed data and dependence on other algorithms. To address these issues, we’ve pre-processed data for various species and incorporated necessary tools (e.g., STAR, StringTie, RNASamba) directly within FREDDIE, minimizing user effort for installation and version compatibility. Future updates will expand the list of supported species to accommodate user needs.

Our results have far-reaching implications for understanding the importance of TEs and retrocopies exonization and its role in disease [34,35]. Enriching chimeric transcripts at specific loci and their variable expression profiles suggests that TEs could be more involved in shaping the transcriptome landscape than previously thought. FREDDIE may open new avenues for future research to explore TE-driven chimerism’s functional consequences.

In conclusion, FREDDIE is a complete, user-friendly, and robust pipeline for detecting exonization events in chimeric transcripts involving TEs and retrocopies. Given the widespread presence of these elements within mammalian genes, FREDDIE can potentially drive discoveries in basic and translational research, with applications ranging from fundamental research to disease contexts, particularly in cancer studies.

