## Supplementary material for "FREDDIE: A comprehensive tool for detecting exonization of retrotransposable elements in short and long RNA sequencing data"

FREDDIE: a tool to identify exonization of retrotransposable elements using RNA-seq data.

### Further information about FREDDIE

In general, FREDDIE is a straightforward pipeline designed to identify chimeric transcripts of retroelements fixed in genomes based on RNA sequencing data. The program was written in Python3 (version 3.6), R (version 4.2.1) and Bash. FREDDIE receives as input either single-end, paired-end or long-reads FASTQ files and performs a set of data analyses, from data processing and alignment (both short and also long-reads) until the discovery of possible functional roles of newly found chimeric transcripts. In the end, two files, besides several figures, are generated: a TSV and a GTF file, which contains an assembly transcriptome, in addition to a handful of other information regarding each one of the transcripts. The source code is distributed under the GNU General Public License.

#### How does FREDDIE work?

FREDDIE has seven subcommands: *star*, *string*, *chimeric*, *coding*, *pfam*, *expression* and *results*.

The input for the "star" subcommand is a list of FASTQ files, which in turn are aligned to a reference genome (hg38) with the STAR aligner ("longSTAR" for long-reads) using default parameters, except for the following: --outSAMstrandField "intronMotif" and --outSAMattributes "NH HI AS nM MD XS". The output of this step is a sorted BAM file filtered by unique alignments (-q 255).

In the second step, the "string" subcommand, FREDDIE performs an assembly guided by StrigTie2 using the BAM file previously generated in the "star" step. This step has as output one GFT (merge.gtf) file containing the transcriptome of all files uploaded by the user.

Sequentially, in the third step, "chimeric", FREDDIE selects all the new transcripts assembled which have an event chosen by the user. As output in this step is a GTF and a fasta file with all chimeric events identified.

The fasta file generated in the previous step is use in the #4 step (“coding”). The pipeline will use RNASamba tool to classify if the chimeric transcript is or not a coding and predict which is the most probable open reading frame (ORF).

Then, the “pfam” step performs a comparison between the coding chimeric transcript identified and the host gene domains. This step uses HMMER to determine the domains.

The “expression” step uses the merge gtf generated in the “string” step to calculate the expression by samples and create a TPM matrix. And to collapse all this information obtained by the pipeline the user should use the “results” step.

#### Installing FREDDIE

The source code for FREDDIE can be obtained in our github page using this command line:

```
$ git clone https://github.com/galantelab/freddie.git
```

FREDDIE is distributed under the GNU General Public License.

Inside FREDDIE’s directory, build a docker image:

```
$ cd freddie
$ docker build -f Dockerfile -t freddie .
```

Set an alias as “freddie”:

```
$ alias freddie='docker run --rm -u $(id -u):$(id -g) -w $(pwd) -v <reference-files-path>:/home/ref/ -v <input-path>:/home/input/ -v <output-path>:/home/output/ -v <output-tostring-path>:/home/output_str freddie'
```

### Using FREDDIE

#### General Syntax

FREDDIE has seven subcommands: “star”, “string”, “chimeric”, “coding”, “pfam”, “expression” and “results”.

To verify the installation process and execute an initial example, users can invoke it in the following manner:

```
freddie 1.0.0

Usage: freddie [-hV]
Usage: freddie <command> [options]

Options:
  -h, --help          Print this help
  -V, --version       Print current version

Commands:
  star                Align RNA-seq data against the genome using
                      STAR (DOI: 10.1093/bioinformatics/bts635)
  string              Assemble sequenced reads (compatible with
                      both short and long reads) using StringTie2
                      (DOI: 10.1186/s13059-019-1910-1)
  chimeric            Identify potential chimeric transcripts
  coding              Compute the coding potential of (chimeric)
                      transcripts using RNASamba
                      (DOI: https://doi.org/10.1093/nargab/lqz024)
  pfam                Search for protein domains using HMMer
                      (DOI: 10.1093/nar/gkr367) and Pfam protein
                      families and domains
                      (https://doi.org/10.1093/nar/gkaa913)
  expression          Estimate transcript expression using
                      StringTie2 (DOI: 10.1186/s13059-019-1910-1)
  results             Compile the final results of chimeric
                      transcripts incorporating inputs from
                      previous steps
```

#### FREDDIE: star

The first step in the FREDDIE's pipeline is the "star". The inputs to this command are FASTQ files and a STAR index (pre-made available at: www.bioinfo.mochsl.org.br/freddiesdata/STAR_index/). The sorted and filtered BAM aligned file resulting from this command will become the input to the next step. This command supports all types of RNA-Seq data (paired-end, single-end and long-reads), either compressed (.gz) or not. This below command shows all possible options for this step:

```
$ freddie star
```

**Arguments:**
One or more sequencing file(s) in FASTQ format.

Mandatory Options:

```
-o, --output-dir
```

> Output directory. Creates the directory if it does not exist

```
-i, --index-dir
```

> STAR index directory

```
-f, --file
```

> File containing a newline separated list of sequencing files in FASTQ format. This option is not mandatory if one or more FASTQ files are passed as argument

Options:

```
-h, --help
```

> Print this help

```
-t, --threads
```

> Number of threads [default: 8]

```
-S, --short-reads
```

> Set the sequencing to short reads [default]

```
-L, --long-reads
```

> Set the sequencing to long reads

```
-s, --single-end
```

> For short reads '-S', set the type of sequencing to single-end

```
-p, --paired-end
```

> For short reads '-S', set the type of sequencing to paired-end. In this case, the FASTQ files will be processed, being considered forward (R1) and reverse complement (R2) according to the order in which they are passed [default]

For example, if the user has 2 FASTQ files (paired-end):

```
$ cat file.tsv
sample1_R1.fastq.gz        sample1_R2.fastq.gz
sample2_R1.fastq.gz        sample2_R2.fastq.gz

$ freddie star -o TEST -i STAR_INDEX/ -f file.tsv
```

Or 2 independent FASTQ files (single-end):

```
$ cat file.tsv
sample1.fastq.gz
sample2.fastq.gz

$ freddie star -o TEST -i STAR_INDEX/ -f file.tsv -s
```

Or 2 FASTQ files from a long-read sequencing strategy:

```
$ cat file.tsv
sample1.fastq.gz
sample2.fastq.gz

$ freddie star -o TEST -i STAR_INDEX/ -f file.tsv -L
```

#### FREDDIE: string

The next step in the pipeline is “string”. This command performs a transcriptome assembly with the BAMs generated in the previous step (or custom BAMs provided by the user). The output of this analysis is a GTF file representing the transcriptome from all samples.

This command shows all options for this step:

```
$ freddie string
```

**Arguments:**
One or more alignment file(s) in BAM format.

Mandatory Options:

```
-o, --output-dir
```

> Output directory. Creates the directory if it does not exist

```
-a, --annotation
```

> Gene annotation of the reference genome transcriptome in GTF format

```
-f, --file
```

> File containing a newline separated list of alignment files in BAM format. This option is not mandatory if one or more BAM files are passed as argument or if the 'star' command has been executed

Options:

```
-h, --help
```

> Print this help

```
-t, --threads
```

> Number of threads [default: 8]

```
-S, --short-reads
```

> Set the sequencing to short reads [default]

```
-L, --long-reads
```

> `Set the sequencing to long reads`

Here, the user can execute this command without a BAM file or a file with BAM paths:

```
$ freddie string -o TEST -a annotation.gtf
```

Or using a BAM file:

```
$ freddie string -o TEST -a annotation.gtf sample_sorted.bam
```

Or using a file with BAM paths:

```
$ cat files.tsv
bams/sample1_sorted.bam
bams/sample2_sorted.bam

$ freddie string -o TEST -a annotation.gtf -f files.txt
```

#### FREDDIE: chimeric

In the “chimeric” step, the pipeline identifies novel transcripts based on the GTF file generated from the “string” command. Here, FREDDIE uses a list of events provided by the user to find transcripts with overlap between exons and the given events. Again, a GTF file and also a FASTA file with all transcripts found are the outputs provided.

This command shows all options for this step:

```
$ freddie chimeric
```

Mandatory Options:

```
-o, --output-dir
```

> `Output directory. Creates the directory if it does not exist`

```
-a, --annotation
```

> `Gene annotation of the reference genome in GTF format`

```
-g, --genome
```

```
FASTA file of the reference genome,which is the same file used
for reads alignment using STAR
```

```
-e, --stringtie-out
```

```
StringTie2 output events file in BED4
```

Options:

```
-h, --help
```

```
Print this help
```

```
-T, --tmp-dir
```

```
Uses directory for temporaries [default: /tmp]
```

```
-r, --reciprocal
```

```
Criteria for identifying chimeric events is 50% overlap of the
event with the exon and 50% overlap of the exon with the event
```

```
-R, --irreciprocal
```

```
Criteria for identifying chimeric events is 50% overlap of the
event with the exon [default]
```

Here, the user can execute this command with a "-R" default:

```
$ head -n5 event.bed
chrN  1000  1500  X
chrN  2000  2500  Y
chrN  3000  3500  X
chrN  4000  4500  Z
chrN  5000  5500  X

$ freddie chimeric -o TEST -g annotation.gtf -G reference.fa -i
events.bed
```

However, we strongly recommend the "reciprocal" option filter for small events (< 250bp) to guarantee less false positives:

```
$ head -n5 event.bed
chrN  1000  1250  X
chrN  2000  2250  Y
chrN  3000  3250  X
chrN  4000  4250  Z
chrN  5000  5250  X

$ freddie chimeric -o TEST -g annotation.gtf -G reference.fa -i events.bed -r
```

#### FREDDIE: coding

The "coding" command classifies the novel transcripts identified in the "chimeric" step as coding or non coding. Here, FREDDIE uses a model trained by RNASamba (available at: www.bioinfo.mochsl.org.br/freddiesdata/RNASamba_model.hdf5) to calculate the probability of a transcript being coding. In the end, a FASTA file with the protein sequences of all transcripts considered coding by our criteria is created.

This command shows all options for this step:

```
$ freddie coding
```

Mandatory Options:

```
-o, --output-dir
```

> Output directory. Creates the directory if it does not exist

```
-m, --protein-model
```

> File with the model of RNASamba

```
-d, --protein-db
```

> File with the protein sequences

Options:

```
-h, --help
```

> Print this help

```
-P, --probability
```

> Set the cutoff for selecting transcripts considered to be protein-coding, based on the probability provided by RNASamba [default: 0.9]

Here, the user can execute this command:

```
$ freddie coding -o TEST -m RNASamba_model.hd5 -d protein.fa
```

If the user needs (or wants) to reduce the coding filter:

```
$ freddie coding -o TEST -m RNASamba_model.hd5 -d protein.fa -P 0.7
```

Or to increase it:

```
$ freddie coding -o TEST -m RNASamba_model.hd5 -d protein.fa -P 0.99
```

#### FREDDIE: pfam

The “pfam” step searches for protein domains in the novel transcripts that passed the user’s predefined coding probability and subsequently compares them with the host’s protein domains. In order to identify the protein domains, we used HMMER trained with the PFAM database (available at: www.bioinfo.mochsl.org.br/freddiesdata/Pfam-A.hmm). The output of this command is a TSV file comparing the protein domains of the novel transcripts identified with those of the host genes.

This command shows all options for this step:

```
$ freddie pfam
```

Mandatory Options:

```
-o, --output-dir
```

> Output directory. Creates the directory if it does not exist

```
-M, --pfam-model
```

> A database of protein domain families to be used as an index for HMmer tool

Options:

```
-h, --help
```

> Print this help

```
-T, --temp-dir
```

> Uses directory for temporaries [default: /tmp]

```
-t, --threads
```

> Number of threads [default: 4]

```
-E, --e-value
```

> In the HMMER per-target output, reports target sequences with an e-value lesser than NUM [default: 1e-6]

Here, the user can execute this command:

```
$ freddie pfam -o TEST -M Pfam-A.hmm
```

#### FREDDIE: expression

The "expression" command quantifies all the transcriptomes assembled by the StringTie2 "expression" function. The expression results, in TPM (transcript per million) per transcript per sample, are made available as a TSV file.

This command shows all options for this step:

```
$ freddie expression
```

**Arguments:**
One or more alignment file(s) in BAM format.

Mandatory Options:

```
-o, --output-dir
```

> `Output directory. Creates the directory if it does not exist`

```
-f, --file
```

> `File containing a newline separated list of alignment files in BAM format. This option is not mandatory if one or more BAM files are passed as argument or if the 'star' command has been executed`

Options:

```
-h, --help
```

> `Print this help`

```
-T, --temp-dir
```

> `Uses directory for temporaries [default: /tmp]`

```
-t, --threads
```

> `Number of threads [default: 4]`

```
-S, --short-reads
```

> `Set the sequencing to short reads [default]`

```
-L, --long-reads
```

> `Set the sequencing to long reads`

Here, the user can execute this command:

```
$ freddie expression -o TEST
```

#### FREDDIE: results

Finally, the "results" command compiles all relevant information from the previous steps. In addition, if the novel transcripts contribute to the expression of their host genes, this step further generates boxplots to show the relative contribution of such expression patterns.

This command shows all options for this step:

```
$ freddie results
```

Mandatory Options:

```
-o, --output-dir
```

> Output directory. Creates the directory if it does not exist

Options:

```
-h, --help
```

> Print this help

```
-T, --temp-dir
```

> Uses directory for temporaries [default: /tmp]

Here, the user can execute this command:

```
$ freddie results -o TEST
```

### A Practical Workflow

In order to execute FREDDIE, we selected RNA-seq paired-end data of 2 samples related to the cell line K562 from the ENCODE Project.

First, you should download the data:

```
$ mkdir fastq
$ cd fastq/
## ENCLB063ZZZ R1
$ wget \
https://www.encodeproject.org/files/ENCFF001RWF/@@download/ENCFF001RW
F.fastq.gz
$ mv ENCFF001RWF.fastq.gz ENCLB063ZZZ_R1.fastq.gz
## ENCLB063ZZZ R2
$ wget \
```

```
https://www.encodeproject.org/files/ENCFF001RWC/@@download/ENCFF001RW
C.fastq.gz
$ mv ENCFF001RWC.fastq.gz ENCLB063ZZZ_R2.fastq.gz
## ENCLB059ZZZ R1
$ wget \
https://www.encodeproject.org/files/ENCFF001RDE/@@download/ENCFF001RD
E.fastq.gz
$ mv ENCFF001RDE.fastq.gz ENCLB059ZZZ_R1.fastq.gz
## ENCLB059ZZZ R2
$ wget \
https://www.encodeproject.org/files/ENCFF001RCW/@@download/ENCFF001RC
W.fastq.gz
$ mv ENCFF001RCW.fastq.gz ENCLB059ZZZ_R2.fastq.gz
$ cd ..
```

And download the databases of FREDDIE:

```
$ mkdir db
$ cd db/
## STAR Index (Based on Human hg38 - Gencode v36)
$ wget \
https://bioinfohsl-tools.s3.amazonaws.com/freddie/databases/star_inde
x.tar.gz
$ tar -xvf star_index.tar.gz
## Gencode v36 as the human annotation
$ wget \
https://bioinfohsl-tools.s3.amazonaws.com/freddie/databases/gencode.v
36.annotation.gtf
## hg38 as the human reference genome
$ wget \
https://bioinfohsl-tools.s3.amazonaws.com/freddie/databases/hg38.fa
## Aminoacid sequence
$ wget \
https://bioinfohsl-tools.s3.amazonaws.com/freddie/databases/hg38.pep.
fa
## RNASamba model
$ wget \
https://bioinfohsl-tools.s3.amazonaws.com/freddie/databases/human38_m
odel.hdf5
```

```
## HMMer model
$ wget \
https://bioinfohsl-tools.s3.amazonaws.com/freddie/databases/Pfam-A.hm
m
$ wget \
https://bioinfohsl-tools.s3.amazonaws.com/freddie/databases/Pfam-A.hm
m.h3f
$ wget \
https://bioinfohsl-tools.s3.amazonaws.com/freddie/databases/Pfam-A.hm
m.h3i
$ wget \
https://bioinfohsl-tools.s3.amazonaws.com/freddie/databases/Pfam-A.hm
m.h3m
$ wget \
https://bioinfohsl-tools.s3.amazonaws.com/freddie/databases/Pfam-A.hm
m.h3p
## Retrocopies events
wget
https://bioinfohsl-tools.s3.amazonaws.com/rcpedia/downloads/beds/RCP_
9606.bed
cut -f 1,2,3,5 RCP_9606.bed > RCP_9606.bed4
$ cd ..
```

Then, prepare a file with the FASTQ PATHs:

```
$ ls fastq/*fastq.gz | awk '{print "/home/freddie/"$1}' > files.txt
```

After that, install and build a docker image:

```
$ git clone https://github.com/galantelab/freddie.git
$ cd freddie
$ sudo docker build -f Dockerfile -t freddie .
$ cd ..
```

Finally, you will be able to execute FREDDIE as follows:

- “star” step:

```
# Input file: files.txt
```

```
# Output file: K562/star/*sorted.bam
# Average time: 127m13.567s

$ time docker run --rm -u $(id -u):$(id -g) \
        -w $(pwd) \
        -v $PWD:/home/freddie \
        freddie star -o /home/freddie/K562 \
                -i /home/freddie/db/star_index \
                -f /home/freddie/files.txt
```

- “string” step:

```
# Input file: K562/star/*sorted.bam
# Output file: K562/string/merge.gtf
# Average time: 15m44.366s

$ time docker run --rm -u $(id -u):$(id -g) \
        -w $(pwd) \
        -v $PWD:/home/freddie \
        freddie string -o /home/freddie/K562 \
                -a /home/freddie/db/gencode.v36.annotation.gtf
```

- “chimeric” step:

```
# Input file: K562/string/merge.gtf
# Output file: K562/chimeric/chimeric.fasta
# Average time: 2m28.100s

$ time docker run --rm -u $(id -u):$(id -g) \
        -w $(pwd) \
        -v $PWD:/home/freddie \
        freddie chimeric -o /home/freddie/K562 \
                -a /home/freddie/db/gencode.v36.annotation.gtf \
                -g /home/freddie/db/hg38.fa \
                -e /home/freddie/db/RCP_9606.bed4
```

- “coding” step:

```
# Input file: K562/chimeric/chimeric.fasta
# Output file1: K562/coding/novel_proteins.fa
# Output file2: K562/coding/ann_proteins.fa
# Average time: 0m12.679s

$ time docker run --rm -u $(id -u):$(id -g) \
        -w $(pwd) \
        -v $PWD:/home/freddie \
        freddie coding -o /home/freddie/K562 \
                -m /home/freddie/db/human38_model.hdf5 \
                -d /home/freddie/db/hg38.pep.fa
```

- “pfam” step:

```
# Input file1: K562/coding/novel_proteins.fa
# Input file2: K562/coding/ann_proteins.fa
# Output file: K562/pfam/info_dom.tsv
# Average time: 1m52.118s

$ time docker run --rm -u $(id -u):$(id -g) \
        -w $(pwd) \
        -v $PWD:/home/freddie \
        freddie pfam -o /home/freddie/K562 \
                -M /home/freddie/db/Pfam-A.hmm
```

- “expression” step:

```
# Input file1: K562/star/*sorted.bam
# Input file2: K562/string/merge.gtf
# Output file: K562/expression/expression.tsv
# Average time: 7m33.230s

$ time docker run --rm -u $(id -u):$(id -g) \
        -w $(pwd) \
        -v $PWD:/home/freddie \
        freddie expression -o /home/freddie/K562
```

- “results” step:

```
# Output file: K562/results/results.tsv
# Average time: 1m42.659s

$ time docker run --rm -u $(id -u):$(id -g) \
        -w $(pwd) \
        -v $PWD:/home/freddie \
        freddie results -o /home/freddie/K562
```

All information related to the chimeric transcripts identified by FREDDIE are available in the final output named “results.tsv”. If you wish to inspect these transcripts in a Genome Browser such as at UCSC (https://genome.ucsc.edu/cgi-bin/hgGateway) you can easily upload the “K562/chimeric/chimeric.gtf” file, also provided by the FREDDIE’s pipeline, to the “custom tracks”.

### Creating index and databases

All indexes and databases from human (hg38), chimpanzee (panTro6), cow (bosTau9), dog (ROS_Cfam_1.0), marmoset (calJac4), mouse (mm39), opossum (monDom5), platypus (mOrnAna1.p.v1), rat (rn6), and rhesus (rheMac10) are available at https://bioinfohsl-tools.s3.amazonaws.com/freddie/databases/ . Despite that, users are welcome to easily create their own indexes and other databases to be used with FREDDIE.

Please note that to execute the commands below it is necessary to previously install the appropriate corresponding programs.

#### STAR Index

To create a STAR index, use the following command:

```
$ STAR --runThreadN 4 --runMode genomeGenerate --genomeDir .
--genomeFastaFiles reference.fa --sjdbGTFfile gencode.gtf > log
```

The user needs:

- A Reference Genome (reference.fa in the command);
- A GTF file (gencode.gtf in the command; We recommend the RefSeq GTFs for any organism).

#### RNASamba Model

To create a training model for RNASamba, use the command below:

```
$ rnasamba train -v 2 model.hdf5 coding_transcripts.fa lncRNA_transcripts.fa
```

The user needs:

- Output name (model.hdf5)
- Coding transcripts FASTA file (coding_transcripts.fa in the command);
- Non-coding transcripts FASTA file (lncRNA_transcripts.fa in the command).

PS: Be careful when running this command. It generally consumes all CPUs available!

#### Protein Annotation

#### HMMER model

To create a hmm model for HMMER, users can run the command:

```
$ hmmbuild model.hmm align.sto
```

The user needs:

- Output name (model.hmm)
- Stockholm align (align.sto)
