## Supplementary Figures and Tables for "FREDDIE: A comprehensive tool for detecting exonization of retrotransposable elements in short and long RNA sequencing data"

### Retrocopy

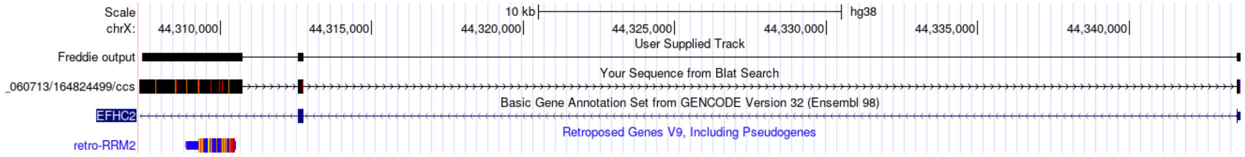

### *Alu*

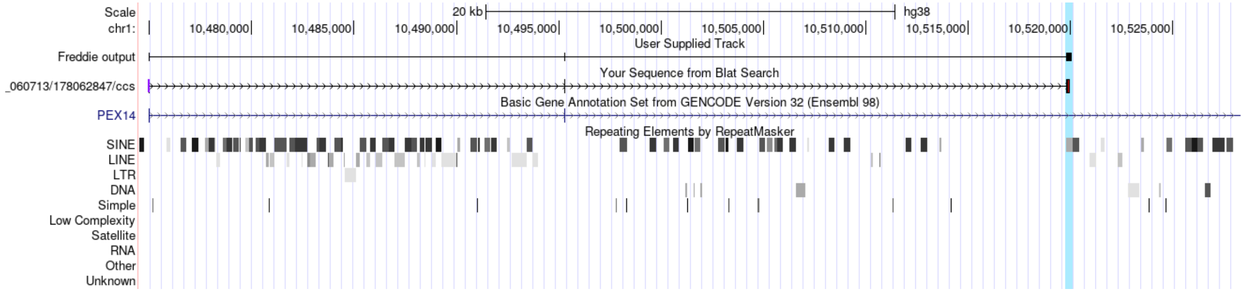

### LINE1

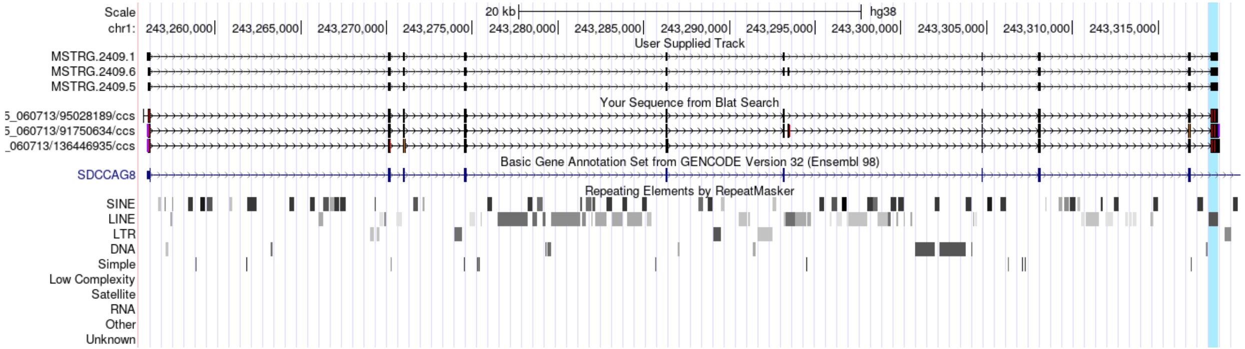

**Figure S1. Examples of chimeric validation by long-reads for each type of TEs and retrocopy selected.** A Genome browser view of the genomic region of the chimeric transcripts. Where in the first track is related to freddie output (Named as “Freddie Output” in retrocopy and Alu examples and by MSTRG prefix in LINE1 example). In the second track is the long-read alignment. The third track is the GENCODEv32 annotation of the host gene. And the last one is the RepeatMasker track showing the TEs and in the retrocopy example is the Retroposed Gene V9.

**Table S1. Primers to reverse transcription-PCR**

| Primer | Sequence |
|---|---|
| HBB (F) | CACCTTTGCCACACTGAGTGAG |
| HBB (R) | CCACTTTCTGATAGGCAGCCTG |
| K562_1_F | AAGGAGATGGAAGTGATGTACC |
| K562_1_R | GACGGAGAGCATGGTGGTAG |
| K562_2_F | GGCCCCTCACTCTCCTTTGC |
| K562_2_R | TCCGCCTTCTCATACATCTGC |

| | |
|---|---|
| K562_3_F | CGCTGCTGCACATTACGGA |
| K562_3_R | CCATGGCAATTTGGAAGCCA |
| K562_4_F | TTGGTGGAGCGATTTAGCCA |
| K562_4_R | TTGGAAAACGTGAGGCCAGG |
| K562_5_F | AAGAAACGAGGACCGATGCC |
| K562_5_R | ACACCACTCCCATCCTCTGA |
| K562_6_F | CTAGCCACACCATGAATTGTCC |
| K562_6_R | GCATCTTTCAACTTTCAACTCCCT |
| K562_7_F | CATTTAACCAGCACAGGGAGT |
| K562_7_R | AATGGAACGGCTTCTCACACA |
| K562_8_F | TGTGTGAGAAGCCGTTCCATTT |
| K562_8_R | CTCACACAACGTACTTACACACA |
| K562_9_F | GGTGGTTCCTACAAGTTGTTGTT |
| K562_9_R | TGTCTCAGCTTTCTTCTCCCA |
| K562_10_F | TGGGAGAAGAAAGCTGAGACA |
| K562_10_R | ACAGGACAATGTTCAAATAGCGA |
| K562_11_F | CGCTATTTGAACATTGTCCTGTTCT |
| K562_11_R | AGCTACTGGAAAAACTGGCAACA |
| K562_12_F | ACGTTTTCAACTTGTTGCCAGT |
| K562_12_R | TGAGCAAGGCATTTTGGTGG |
| K562_13_F | TTGGGAGAAGAAAGCTGAGACA |
| K562_13_R | CTTGAATTGTGTCATCTTCAGGGT |
| U251_1_F | GGATGGAAAGCCACCCGTAA |
| U251_1_R | GTGAATGGACTTCCCGGTCT |
| U251_2_F | TGCTAACATGCCTTGGCTCA |
| U251_2_R | CAGCTGTTTCCATTGGTGGG |
| U251_3_F | TGGACACGTAAATTCGGGCA |
| U251_3_R | GGTGGCTTTCCATCCCTTGA |
| U251_4_F | CCCACAATGGGTTTGCCAC |
| U251_4_R | TGCCCGAATTTACGTGTCCA |
| U251_5_F | AAGGGAAAGGGGTGGCTCAT |
| U251_5_R | TGTTCTGGTGGCAAACCCAT |
| U251_6_F | GCCCTTCTGGCTTACACACT |
| U251_6_R | TCCCAAAATAAGCCCCTTCTGA |
| U251_1_F | AACACAAGATGGGGACTCCTG |

| | |
|---|---|
| U251_1_R | GGCATTTCACTTTGGAAGGCA |
| U251_2_F | GCTCGAAACTGGACTTCCTGA |
| U251_2_R | CCATGGGCCTGCAATCTTCT |
| U251_3_F | GGCCCATGGACTGTGTGTTAC |
| U251_3_R | TCCATCTACAACGGGCAGTC |
| U251_4_F | TCTTCATGACTGCCCGTTGT |
| U251_4_R | CATGAGCATCACTTCGGGAGA |
| U251_5_F | AAATTGAGGCTGGAGCTTTCT |
| U251_5_R | ACATGTGGACCTGAGAGCAA |
| U251_6_F | CGTACTGTTTAGGCTGTTGGTG |
| U251_6_R | CGTTCCAGTTCTCTCATGCGT |

**Table S2. Chimeric transcripts identified by FREDDIE.**

| Transcript_id | Cell Line | Genomic Position | Retroelement | Gene_Host |
|---|---|---|---|---|
| MSTRG.476.7 | K562 | chr1:27322281-27336595 | ACTG1P20 | ENSG00000186501.14 |
| MSTRG.212.3 | K562 | chr1:10474950-10520094 | Alu | ENSG00000142655.13 |
| MSTRG.1502.6 | U251 | chr1:109656122-109666420 | Alu | ENSG00000168765.17 |
| MSTRG.1502.11 | U251 | chr1:109656390-109666420 | Alu | ENSG00000168765.17 |
| MSTRG.1502.14 | U251 | chr1:109656721-109666420 | Alu | ENSG00000168765.17 |
| MSTRG.1543.22 | U251 | chr1:112575004-112619139 | Alu | ENSG00000007341.19 |
| MSTRG.36.13 | U251 | chr1:1255264-1262169 | Alu | ENSG00000160087.20 |
| MSTRG.1676.11 | U251 | chr1:144421494-144461674 | Alu | ENSG00000266338.6 |
| MSTRG.1549.2 | K562 | chr1:149054033-149102884 | Alu | ENSG00000269713.7 |
| MSTRG.1794.5 | U251 | chr1:149054033-149103169 | Alu | ENSG00000269713.7 |
| MSTRG.1794.3 | U251 | chr1:149054033-149103685 | Alu | ENSG00000269713.7 |
| MSTRG.1794.10 | U251 | chr1:149054047-149103685 | Alu | ENSG00000269713.7 |
| MSTRG.1794.12 | U251 | chr1:149054062-149103685 | Alu | ENSG00000269713.7 |
| MSTRG.1794.13 | U251 | chr1:149054112-149103685 | Alu | ENSG00000269713.7 |
| MSTRG.1794.17 | U251 | chr1:149059035-149103169 | Alu | ENSG00000269713.7 |
| MSTRG.1794.18 | U251 | chr1:149059763-149103169 | Alu | ENSG00000269713.7 |
| MSTRG.1794.20 | U251 | chr1:149091429-149103169 | Alu | ENSG00000269713.7 |
| MSTRG.1794.21 | U251 | chr1:149093034-149103685 | Alu | ENSG00000269713.7 |
| MSTRG.1832.2 | U251 | chr1:150620611-150629612 | Alu | ENSG00000143420.18 |
| MSTRG.1866.6 | U251 | chr1:151284855-151292176 | Alu | ENSG00000143373.18 |

| | | | | |
|---|---|---|---|---|
| MSTRG.1695.10 | K562 | chr1:155199346-155202458 | Alu | ENSG00000169231.13 |
| MSTRG.87.1 | K562 | chr1:1784988-1880272 | Alu | ENSG00000078369.18 |
| MSTRG.87.9 | K562 | chr1:1785286-1880272 | Alu | ENSG00000078369.18 |
| MSTRG.356.5 | K562 | chr1:20742679-20747101 | Alu | ENSG00000127483.18 |
| MSTRG.2275.12 | K562 | chr1:224228973-224330189 | Alu | ENSG00000143748.18 |
| MSTRG.2710.3 | U251 | chr1:225486849-225653142 | Alu | ENSG00000154380.17 |
| MSTRG.2710.8 | U251 | chr1:225496885-225653142 | Alu | ENSG00000154380.17 |
| MSTRG.2710.15 | U251 | chr1:225500802-225590396 | Alu | ENSG00000154380.17 |
| MSTRG.499.6 | U251 | chr1:26256719-26279236 | Alu | ENSG00000130695.15 |
| MSTRG.652.14 | U251 | chr1:32073959-32094971 | Alu | ENSG00000121775.18 |
| MSTRG.589.17 | K562 | chr1:32820399-32858879 | Alu | ENSG00000116497.18 |
| MSTRG.1059.2 | K562 | chr1:75202134-75660240 | Alu | ENSG00000137968.16 |
| MSTRG.3907.14 | U251 | chr10:101791676-101843920 | Alu | ENSG00000198408.14 |
| MSTRG.4159.15 | U251 | chr10:132348746-132365854 | Alu | ENSG00000148814.18 |
| MSTRG.2744.7 | K562 | chr10:42589119-42638154 | Alu | ENSG00000196693.15 |
| MSTRG.2484.11 | K562 | chr10:4826730-4848278 | Alu | ENSG00000165568.18 |
| MSTRG.2862.12 | K562 | chr10:50082430-50133673 | Alu | ENSG00000099290.17 |
| MSTRG.2915.16 | K562 | chr10:68489023-68527523 | Alu | ENSG00000122912.15 |
| MSTRG.2930.3 | K562 | chr10:68636472-68694487 | Alu | ENSG00000138336.9 |
| MSTRG.3003.9 | K562 | chr10:73442962-73496024 | Alu | ENSG00000107758.15 |
| MSTRG.3087.3 | K562 | chr10:87095196-87191468 | Alu | ENSG00000122376.11 |
| MSTRG.3697.4 | U251 | chr10:87095203-87104836 | Alu | ENSG00000122376.11 |
| MSTRG.3161.1 | K562 | chr10:94321955-94362968 | Alu | ENSG00000173145.11 |
| MSTRG.3186.8 | K562 | chr10:96130509-96164028 | Alu | ENSG00000177853.14 |
| MSTRG.3771.5 | K562 | chr11:17354862-17383674 | Alu | ENSG00000188211.9 |
| MSTRG.3771.6 | K562 | chr11:17355455-17380648 | Alu | ENSG00000188211.9 |
| MSTRG.4788.17 | U251 | chr11:62669214-62671774 | Alu | ENSG00000162194.12 |
| MSTRG.4791.3 | U251 | chr11:62728069-62739947 | Alu | ENSG00000162222.14 |
| MSTRG.4115.10 | K562 | chr11:64226300-64228266 | Alu | ENSG00000149761.9 |
| MSTRG.3520.6 | K562 | chr11:651498-694961 | Alu | ENSG00000177030.16 |
| MSTRG.4433.11 | K562 | chr11:77924073-77994671 | Alu | ENSG00000149262.17 |
| MSTRG.4399.16 | U251 | chr11:9956568-10300495 | Alu | ENSG00000133812.15 |
| MSTRG.5833.2 | K562 | chr12:101745508-101830959 | Alu | ENSG00000111670.16 |
| MSTRG.7267.13 | U251 | chr12:122936254-122966509 | Alu | ENSG00000150967.18 |
| MSTRG.7267.14 | U251 | chr12:122936421-122951005 | Alu | ENSG00000150967.18 |

| | | | | |
|---|---|---|---|---|
| MSTRG.7267.15 | U251 | chr12:122936523-122953084 | Alu | ENSG00000150967.18 |
| MSTRG.7265.6 | U251 | chr12:123258097-123271856 | Alu | ENSG00000111328.7 |
| MSTRG.6135.1 | K562 | chr12:123601537-123620943 | Alu | ENSG00000111364.16 |
| MSTRG.6210.7 | K562 | chr12:131723258-131739342 | Alu | ENSG00000061936.9 |
| MSTRG.5152.14 | K562 | chr12:28225285-28581511 | Alu | ENSG00000123106.10 |
| MSTRG.5758.21 | U251 | chr12:2890905-2940529 | Alu | ENSG00000078246.17 |
| MSTRG.5758.23 | U251 | chr12:2904394-2941142 | Alu | ENSG00000078246.17 |
| MSTRG.6324.2 | U251 | chr12:49988798-50025450 | Alu | ENSG00000161800.13 |
| MSTRG.6417.5 | U251 | chr12:53381011-53416446 | Alu | ENSG00000185591.10 |
| MSTRG.6491.1 | U251 | chr12:55901413-55927267 | Alu | ENSG00000170473.17 |
| MSTRG.6638.10 | U251 | chr12:63590865-63668534 | Alu | ENSG00000177990.12 |
| MSTRG.5782.11 | U251 | chr12:6459893-6462243 | Alu | ENSG00000139192.12 |
| MSTRG.8021.15 | U251 | chr13:114282561-114305817 | Alu | ENSG00000169062.15 |
| MSTRG.8021.16 | U251 | chr13:114284088-114305817 | Alu | ENSG00000169062.15 |
| MSTRG.7461.15 | U251 | chr13:24280032-24307074 | Alu | ENSG00000182957.16 |
| MSTRG.6374.6 | K562 | chr13:40943440-41061440 | Alu | ENSG00000120690.16 |
| MSTRG.9047.8 | U251 | chr14:102322394-102342702 | Alu | ENSG00000022976.15 |
| MSTRG.9125.39 | U251 | chr14:103629361-103694559 | Alu | ENSG00000126214.21 |
| MSTRG.8137.12 | U251 | chr14:23306906-23317235 | Alu | ENSG00000129473.9 |
| MSTRG.8383.8 | U251 | chr14:35090757-35122517 | Alu | ENSG00000092020.10 |
| MSTRG.8631.2 | U251 | chr14:69379925-69397940 | Alu | ENSG00000100632.11 |
| MSTRG.7250.9 | K562 | chr14:70352972-70417090 | Alu | ENSG00000258644.6 |
| MSTRG.8737.5 | U251 | chr14:73851864-73862918 | Alu | ENSG00000140043.11 |
| MSTRG.8768.17 | U251 | chr14:75053054-75064033 | Alu | ENSG00000119640.9 |
| MSTRG.8797.38 | U251 | chr14:75882719-75955249 | Alu | ENSG00000119685.20 |
| MSTRG.9452.7 | U251 | chr15:42575714-42615358 | Alu | ENSG00000159433.12 |
| MSTRG.9771.23 | U251 | chr15:55454864-55498584 | Alu | ENSG00000256061.7 |
| MSTRG.9771.24 | U251 | chr15:55454896-55498584 | Alu | ENSG00000256061.7 |
| MSTRG.9754.1 | U251 | chr15:59119734-59372871 | Alu | ENSG00000157483.9 |
| MSTRG.9871.2 | U251 | chr15:66386837-66485558 | Alu | ENSG00000169032.9 |
| MSTRG.9949.11 | U251 | chr15:71889472-72117883 | Alu | ENSG00000066933.16 |
| MSTRG.8354.4 | K562 | chr15:77108164-77388615 | Alu | ENSG00000173517.10 |
| MSTRG.8354.3 | K562 | chr15:77108164-77420210 | Alu | ENSG00000173517.10 |
| MSTRG.10101.8 | U251 | chr15:78787136-78798113 | Alu | ENSG00000136378.15 |
| MSTRG.10101.9 | U251 | chr15:78787380-78798113 | Alu | ENSG00000136378.15 |

| | | | | |
|---|---|---|---|---|
| MSTRG.8701.4 | K562 | chr16:1833973-1872178 | Alu | ENSG00000162039.15 |
| MSTRG.8701.6 | K562 | chr16:1833983-1872178 | Alu | ENSG00000162039.15 |
| MSTRG.10864.10 | U251 | chr16:22421709-22435812 | Alu | ENSG00000140743.8 |
| MSTRG.9190.6 | K562 | chr16:29906963-29926236 | Alu | ENSG00000174943.11 |
| MSTRG.9357.14 | K562 | chr16:47593327-47701523 | Alu | ENSG00000102893.16 |
| MSTRG.11217.22 | U251 | chr16:53239735-53329422 | Alu | ENSG00000177200.17 |
| MSTRG.10431.29 | U251 | chr16:664789-667833 | Alu | ENSG00000161996.19 |
| MSTRG.11458.7 | U251 | chr16:74408634-74414076 | Alu | ENSG00000140839.11 |
| MSTRG.11774.1 | U251 | chr16:90004673-90020157 | Alu | ENSG00000003249.13 |
| MSTRG.11786.5 | U251 | chr17:1464212-1487814 | Alu | ENSG00000197879.17 |
| MSTRG.12361.3 | U251 | chr17:30832356-30856640 | Alu | ENSG00000176208.9 |
| MSTRG.10552.11 | K562 | chr17:36500895-36535251 | Alu | ENSG00000278259.4 |
| MSTRG.12556.9 | U251 | chr17:38737587-38748195 | Alu | ENSG00000277258.5 |
| MSTRG.12572.5 | U251 | chr17:38776119-38799905 | Alu | ENSG00000276293.4 |
| MSTRG.13134.6 | U251 | chr17:64507011-64518040 | Alu | ENSG00000258890.7 |
| MSTRG.13243.21 | U251 | chr17:67898212-67984584 | Alu | ENSG00000171634.18 |
| MSTRG.13349.3 | U251 | chr17:76679569-76692208 | Alu | ENSG00000182534.13 |
| MSTRG.13485.27 | U251 | chr17:82256044-82273731 | Alu | ENSG00000141551.14 |
| MSTRG.11595.12 | K562 | chr18:23529596-23531211 | Alu | ENSG00000141452.10 |
| MSTRG.11729.5 | K562 | chr18:50257583-50281574 | Alu | ENSG00000141644.17 |
| MSTRG.14564.3 | U251 | chr19:14566087-14568276 | Alu | ENSG00000099795.7 |
| MSTRG.14626.7 | U251 | chr19:17267437-17278053 | Alu | ENSG00000105393.16 |
| MSTRG.12390.6 | K562 | chr19:17331522-17334855 | Alu | ENSG00000074855.11 |
| MSTRG.14157.1 | U251 | chr19:1985263-1989337 | Alu | ENSG00000133243.10 |
| MSTRG.15048.3 | U251 | chr19:40320712-40347439 | Alu | ENSG00000160392.13 |
| MSTRG.12806.1 | K562 | chr19:43606822-43608939 | Alu | ENSG00000131116.12 |
| MSTRG.13176.13 | K562 | chr19:55158663-55166890 | Alu | ENSG00000167646.13 |
| MSTRG.14306.11 | U251 | chr19:6584103-6590475 | Alu | ENSG00000125726.11 |
| MSTRG.11865.8 | K562 | chr19:959130-975939 | Alu | ENSG00000116017.11 |
| MSTRG.12139.1 | K562 | chr19:9641664-9658668 | Alu | ENSG00000171466.10 |
| MSTRG.16753.3 | U251 | chr2:132654219-132671579 | Alu | ENSG00000150551.11 |
| MSTRG.14434.9 | K562 | chr2:172558454-172629793 | Alu | ENSG00000152256.13 |
| MSTRG.17458.15 | U251 | chr2:218659719-218667681 | Alu | ENSG00000074582.14 |
| MSTRG.14863.10 | K562 | chr2:233168174-233178262 | Alu | ENSG00000168918.14 |
| MSTRG.13439.1 | K562 | chr2:26242781-26290465 | Alu | ENSG00000138029.14 |

| | | | | |
|---|---|---|---|---|
| MSTRG.16276.4 | U251 | chr2:73729485-73737400 | Alu | ENSG00000144034.15 |
| MSTRG.16276.5 | U251 | chr2:73729557-73737301 | Alu | ENSG00000144034.15 |
| MSTRG.14018.12 | K562 | chr2:96875882-96909141 | Alu | ENSG00000168754.15 |
| MSTRG.17893.14 | U251 | chr20:17941647-17957591 | Alu | ENSG00000089006.16 |
| MSTRG.17991.5 | U251 | chr20:33556618-33646159 | Alu | ENSG00000078699.21 |
| MSTRG.18014.22 | U251 | chr20:34516367-34540958 | Alu | ENSG00000125971.16 |
| MSTRG.18040.10 | U251 | chr20:35268691-35278131 | Alu | ENSG00000126005.17 |
| MSTRG.15629.8 | K562 | chr21:17596982-17606084 | Alu | ENSG00000154640.14 |
| MSTRG.15776.25 | K562 | chr21:37154508-37187341 | Alu | ENSG00000182670.13 |
| MSTRG.15762.3 | K562 | chr21:38121473-38172513 | Alu | ENSG00000157551.19 |
| MSTRG.18771.3 | U251 | chr21:46188446-46197852 | Alu | ENSG00000160285.15 |
| MSTRG.18820.7 | U251 | chr22:18079038-18101611 | Alu | ENSG00000215193.12 |
| MSTRG.19162.7 | U251 | chr22:37946163-37953601 | Alu | ENSG00000128346.11 |
| MSTRG.19266.9 | U251 | chr22:41866414-41907307 | Alu | ENSG00000198911.12 |
| MSTRG.19370.3 | U251 | chr22:46296870-46334255 | Alu | ENSG00000075218.19 |
| MSTRG.19370.6 | U251 | chr22:46297055-46334255 | Alu | ENSG00000075218.19 |
| MSTRG.19377.4 | U251 | chr22:49903961-49917839 | Alu | ENSG00000182858.14 |
| MSTRG.20380.5 | U251 | chr3:119468975-119492001 | Alu | ENSG00000163389.12 |
| MSTRG.17459.2 | K562 | chr3:138652699-138776907 | Alu | ENSG00000051382.8 |
| MSTRG.20599.3 | U251 | chr3:138654018-138834938 | Alu | ENSG00000051382.8 |
| MSTRG.21051.19 | U251 | chr3:187028978-187078553 | Alu | ENSG00000073849.15 |
| MSTRG.21108.1 | U251 | chr3:194585205-194628773 | Alu | ENSG00000145014.17 |
| MSTRG.17833.3 | K562 | chr3:195274745-195442307 | Alu | ENSG00000114331.15 |
| MSTRG.16676.1 | K562 | chr3:33388336-33416834 | Alu | ENSG00000153560.12 |
| MSTRG.16798.8 | K562 | chr3:44840705-44874067 | Alu | ENSG00000163808.17 |
| MSTRG.21839.14 | U251 | chr4:105402019-105474067 | Alu | ENSG00000138777.20 |
| MSTRG.21836.6 | U251 | chr4:107670683-107720234 | Alu | ENSG00000138801.9 |
| MSTRG.21965.1 | U251 | chr4:118837257-119061247 | Alu | ENSG00000172403.11 |
| MSTRG.18615.8 | K562 | chr4:127972865-128010808 | Alu | ENSG00000164074.15 |
| MSTRG.21999.2 | U251 | chr4:133149292-133189057 | Alu | ENSG00000138650.9 |
| MSTRG.18804.4 | K562 | chr4:173232452-173323967 | Alu | ENSG00000109586.12 |
| MSTRG.22315.16 | U251 | chr4:182306266-182803024 | Alu | ENSG00000218336.9 |
| MSTRG.21186.16 | U251 | chr4:456051-474167 | Alu | ENSG00000182903.16 |
| MSTRG.18285.5 | K562 | chr4:70902378-70988350 | Alu | ENSG00000173542.8 |
| MSTRG.19570.7 | K562 | chr5:116003702-116027619 | Alu | ENSG00000172901.20 |

| | | | | |
|---|---|---|---|---|
| MSTRG.23307.4 | U251 | chr5:134661119-134727909 | Alu | ENSG00000113615.13 |
| MSTRG.23371.18 | U251 | chr5:138814269-138935174 | Alu | ENSG00000044115.21 |
| MSTRG.19829.9 | K562 | chr5:141969105-142011162 | Alu | ENSG00000013561.18 |
| MSTRG.23599.3 | U251 | chr5:154819750-154831601 | Alu | ENSG00000170271.11 |
| MSTRG.19030.1 | K562 | chr5:34897057-34915626 | Alu | ENSG00000113456.19 |
| MSTRG.19030.2 | K562 | chr5:34897068-34915626 | Alu | ENSG00000113456.19 |
| MSTRG.19030.3 | K562 | chr5:34897068-34915626 | Alu | ENSG00000113456.19 |
| MSTRG.22542.1 | U251 | chr5:34897073-34915626 | Alu | ENSG00000113456.19 |
| MSTRG.22542.2 | U251 | chr5:34897172-34915626 | Alu | ENSG00000113456.19 |
| MSTRG.19090.4 | K562 | chr5:43289395-43313512 | Alu | ENSG00000112972.15 |
| MSTRG.19103.5 | K562 | chr5:43491717-43515148 | Alu | ENSG00000172244.9 |
| MSTRG.19339.13 | K562 | chr5:79614121-79623489 | Alu | ENSG00000164329.13 |
| MSTRG.23078.2 | U251 | chr5:96768097-96784798 | Alu | ENSG00000164307.13 |
| MSTRG.24834.1 | U251 | chr6:105096250-105137157 | Alu | ENSG00000112276.14 |
| MSTRG.21087.3 | K562 | chr6:117689642-117710813 | Alu | ENSG00000153989.8 |
| MSTRG.21102.12 | K562 | chr6:118662112-118710075 | Alu | ENSG00000111860.14 |
| MSTRG.20236.1 | K562 | chr6:17614793-17688618 | Alu | ENSG00000124789.11 |
| MSTRG.20114.1 | K562 | chr6:2885720-2903309 | Alu | ENSG00000170542.6 |
| MSTRG.23892.23 | U251 | chr6:3004897-3028869 | Alu | ENSG00000124588.20 |
| MSTRG.24239.6 | U251 | chr6:31160338-31164331 | Alu | ENSG00000137310.12 |
| MSTRG.24344.10 | U251 | chr6:33261060-33271904 | Alu | ENSG00000223501.9 |
| MSTRG.24375.3 | U251 | chr6:34587299-34688934 | Alu | ENSG00000196821.10 |
| MSTRG.24414.5 | U251 | chr6:35805839-35838145 | Alu | ENSG00000197753.11 |
| MSTRG.24428.9 | U251 | chr6:36898111-36928964 | Alu | ENSG00000198663.16 |
| MSTRG.24481.1 | U251 | chr6:43005173-43013983 | Alu | ENSG00000124733.5 |
| MSTRG.20720.12 | K562 | chr6:43557837-43561008 | Alu | ENSG00000124571.18 |
| MSTRG.20754.7 | K562 | chr6:52281323-52284881 | Alu | ENSG00000112118.20 |
| MSTRG.24826.7 | U251 | chr6:99521534-99535489 | Alu | ENSG00000279170.2 |
| MSTRG.24826.10 | U251 | chr6:99525965-99535489 | Alu | ENSG00000279170.2 |
| MSTRG.22135.10 | K562 | chr7:100853604-100867010 | Alu | ENSG00000146828.18 |
| MSTRG.26392.7 | U251 | chr7:103297443-103306739 | Alu | ENSG00000105819.14 |
| MSTRG.25570.5 | U251 | chr7:10973861-11143488 | Alu | ENSG00000106443.16 |
| MSTRG.25570.6 | U251 | chr7:10973868-11143488 | Alu | ENSG00000106443.16 |
| MSTRG.22431.1 | K562 | chr7:138460217-138589996 | Alu | ENSG00000122779.18 |
| MSTRG.22431.3 | K562 | chr7:138460232-138589996 | Alu | ENSG00000122779.18 |

| | | | | |
|---|---|---|---|---|
| MSTRG.26686.5 | U251 | chr7:138495631-138589996 | Alu | ENSG00000122779.18 |
| MSTRG.26686.6 | U251 | chr7:138495631-138589996 | Alu | ENSG00000122779.18 |
| MSTRG.25477.2 | U251 | chr7:2251770-2311807 | Alu | ENSG00000106266.11 |
| MSTRG.25742.18 | U251 | chr7:32017776-32071205 | Alu | ENSG00000154678.18 |
| MSTRG.25941.5 | U251 | chr7:56010108-56051604 | Alu | ENSG00000146733.14 |
| MSTRG.25941.6 | U251 | chr7:56010114-56051604 | Alu | ENSG00000146733.14 |
| MSTRG.25941.12 | U251 | chr7:56010226-56042349 | Alu | ENSG00000146733.14 |
| MSTRG.25941.14 | U251 | chr7:56010360-56051604 | Alu | ENSG00000146733.14 |
| MSTRG.21826.2 | K562 | chr7:56010484-56051604 | Alu | ENSG00000146733.14 |
| MSTRG.25941.28 | U251 | chr7:56042350-56051604 | Alu | ENSG00000146733.14 |
| MSTRG.21931.22 | K562 | chr7:74752897-74760692 | Alu | ENSG00000263001.6 |
| MSTRG.22056.1 | K562 | chr7:96120220-96309684 | Alu | ENSG00000004864.13 |
| MSTRG.23306.1 | K562 | chr8:101686358-101988056 | Alu | ENSG00000104490.18 |
| MSTRG.27880.5 | U251 | chr8:130052107-130289093 | Alu | ENSG00000153317.15 |
| MSTRG.27880.11 | U251 | chr8:130052111-130289093 | Alu | ENSG00000153317.15 |
| MSTRG.27042.6 | U251 | chr8:22367363-22422762 | Alu | ENSG00000104635.14 |
| MSTRG.22876.1 | K562 | chr8:30578182-30653983 | Alu | ENSG00000197265.9 |
| MSTRG.27247.8 | U251 | chr8:41529261-41531297 | Alu | ENSG00000147536.12 |
| MSTRG.27319.10 | U251 | chr8:53928547-53959303 | Alu | ENSG00000147509.14 |
| MSTRG.27319.11 | U251 | chr8:53928566-53959303 | Alu | ENSG00000147509.14 |
| MSTRG.27411.11 | U251 | chr8:70208001-70245375 | Alu | ENSG00000140396.13 |
| MSTRG.24212.1 | K562 | chr9:112683926-112718149 | Alu | ENSG00000148153.14 |
| MSTRG.29005.16 | U251 | chr9:125052209-125143526 | Alu | ENSG00000173611.17 |
| MSTRG.24317.15 | K562 | chr9:125052214-125143506 | Alu | ENSG00000173611.17 |
| MSTRG.29204.17 | U251 | chr9:136405373-136410614 | Alu | ENSG00000165689.17 |
| MSTRG.23818.3 | K562 | chr9:35782054-35809732 | Alu | ENSG00000159899.14 |
| MSTRG.23846.8 | K562 | chr9:36572882-36677683 | Alu | ENSG00000165304.8 |
| MSTRG.28557.1 | U251 | chr9:74946583-74953412 | Alu | ENSG00000135045.7 |
| MSTRG.24016.3 | K562 | chr9:88388444-88478694 | Alu | ENSG00000106723.17 |
| MSTRG.28720.25 | U251 | chr9:95003917-95087218 | Alu | ENSG00000148120.16 |
| MSTRG.28720.29 | U251 | chr9:95004236-95087218 | Alu | ENSG00000148120.16 |
| MSTRG.24131.5 | K562 | chr9:97691929-97697357 | Alu | ENSG00000136936.10 |
| MSTRG.29349.15 | U251 | chrX:10464216-10620754 | Alu | ENSG00000101871.14 |
| MSTRG.30089.33 | U251 | chrX:111690667-111764602 | Alu | ENSG00000101901.12 |
| MSTRG.30361.1 | U251 | chrX:154401236-154411361 | Alu | ENSG00000013563.14 |

| | | | | |
|---|---|---|---|---|
| MSTRG.30361.13 | U251 | chrX:154403268-154411361 | Alu | ENSG00000013563.14 |
| MSTRG.24962.3 | K562 | chrX:78131806-78139679 | Alu | ENSG00000187325.5 |
| MSTRG.29888.4 | U251 | chrX:78131993-78139667 | Alu | ENSG00000187325.5 |
| MSTRG.30431.3 | U251 | chrY:7303981-7332211 | Alu | ENSG00000099725.14 |
| MSTRG.19194.9 | K562 | chr5:62332565-62348167 | CRTC2P1 | ENSG00000068796.16 |
| MSTRG.17512.6 | K562 | chr3:142449238-142553398 | EIF2AK1P1 | ENSG00000175054.16 |
| MSTRG.1140.1 | K562 | chr1:88659495-88836255 | ELOCP19 | ENSG00000065243.20 |
| MSTRG.4394.16 | U251 | chr11:9956568-10300495 | GLULP8 | ENSG00000133812.15 |
| MSTRG.8232.5 | U251 | chr14:30893896-30964509 | HIGD1AP17 | ENSG00000196792.12 |
| MSTRG.23395.7 | K562 | chr8:123072723-123123682 | HMGB1P19 | ENSG00000156787.17 |
| MSTRG.8543.8 | U251 | chr14:58427709-58522247 | HNRNPCP1 | ENSG00000100578.17 |
| MSTRG.8543.23 | U251 | chr14:58459864-58522247 | HNRNPCP1 | ENSG00000100578.17 |
| MSTRG.21813.6 | U251 | chr4:102631534-102730171 | KRT8P46 | ENSG00000109323.11 |
| MSTRG.1676.8 | U251 | chr1:144421487-144442127 | L1 | ENSG00000266338.6 |
| MSTRG.1478.4 | K562 | chr1:147928436-147980088 | L1 | ENSG00000188092.15 |
| MSTRG.415.18 | U251 | chr1:20753862-20787323 | L1 | ENSG00000127483.18 |
| MSTRG.2893.1 | U251 | chr1:243255927-243318486 | L1 | ENSG00000054282.16 |
| MSTRG.2409.1 | K562 | chr1:243256034-243318523 | L1 | ENSG00000054282.16 |
| MSTRG.2893.4 | U251 | chr1:243256041-243318486 | L1 | ENSG00000054282.16 |
| MSTRG.2409.5 | K562 | chr1:243256053-243318523 | L1 | ENSG00000054282.16 |
| MSTRG.2409.6 | K562 | chr1:243256053-243318523 | L1 | ENSG00000054282.16 |
| MSTRG.1084.9 | K562 | chr1:77779853-77789133 | L1 | ENSG00000180488.15 |
| MSTRG.1094.14 | K562 | chr1:77955255-77979110 | L1 | ENSG00000162613.16 |
| MSTRG.1251.14 | U251 | chr1:78656909-78664078 | L1 | ENSG00000137965.11 |
| MSTRG.1251.15 | U251 | chr1:78657226-78664078 | L1 | ENSG00000137965.11 |
| MSTRG.221.10 | U251 | chr1:9848277-9894431 | L1 | ENSG00000178585.15 |
| MSTRG.3378.7 | U251 | chr10:45975219-46003876 | L1 | ENSG00000265354.4 |
| MSTRG.3161.2 | K562 | chr10:94332763-94345247 | L1 | ENSG00000173145.11 |
| MSTRG.4602.1 | K562 | chr11:108154744-108205417 | L1 | ENSG00000149308.17 |
| MSTRG.4614.12 | U251 | chr11:43680738-43816910 | L1 | ENSG00000149084.13 |
| MSTRG.4669.1 | U251 | chr11:47214465-47223136 | L1 | ENSG00000134574.11 |
| MSTRG.3683.5 | K562 | chr11:8612123-8664255 | L1 | ENSG00000166436.16 |
| MSTRG.5222.3 | K562 | chr12:39626184-39693491 | L1 | ENSG00000180116.15 |
| MSTRG.6659.2 | U251 | chr12:66302579-66343455 | L1 | ENSG00000127311.9 |
| MSTRG.7960.14 | K562 | chr15:49622292-49637228 | L1 | ENSG00000104047.15 |

| | | | | |
|---|---|---|---|---|
| MSTRG.9887.5 | U251 | chr15:66708308-66782849 | L1 | ENSG00000137834.15 |
| MSTRG.8889.20 | K562 | chr16:10454764-10483638 | L1 | ENSG00000166669.13 |
| MSTRG.11245.2 | U251 | chr16:58249936-58290676 | L1 | ENSG00000103021.9 |
| MSTRG.11684.6 | U251 | chr16:87830025-87866330 | L1 | ENSG00000103257.9 |
| MSTRG.12033.1 | U251 | chr17:11953889-11996151 | L1 | ENSG00000154957.14 |
| MSTRG.12068.1 | U251 | chr17:15699577-15709568 | L1 | ENSG00000187607.16 |
| MSTRG.12112.10 | U251 | chr17:15976569-15991441 | L1 | ENSG00000214941.8 |
| MSTRG.12115.10 | U251 | chr17:16031404-16198021 | L1 | ENSG00000141027.21 |
| MSTRG.12115.11 | U251 | chr17:16031404-16198021 | L1 | ENSG00000141027.21 |
| MSTRG.11560.12 | K562 | chr18:14119916-14132490 | L1 | ENSG00000175322.11 |
| MSTRG.11534.1 | K562 | chr18:9914002-9960120 | L1 | ENSG00000101558.13 |
| MSTRG.11534.8 | K562 | chr18:9914658-9960120 | L1 | ENSG00000101558.13 |
| MSTRG.14555.5 | U251 | chr19:14408850-14415459 | L1 | ENSG00000123136.15 |
| MSTRG.14784.7 | U251 | chr19:21359014-21377034 | L1 | ENSG00000172687.13 |
| MSTRG.12521.9 | K562 | chr19:23658965-23687220 | L1 | ENSG00000197372.10 |
| MSTRG.15592.7 | U251 | chr19:58567100-58572775 | L1 | ENSG00000099326.9 |
| MSTRG.14329.6 | U251 | chr19:7982650-8005659 | L1 | ENSG00000066044.15 |
| MSTRG.16688.11 | U251 | chr2:120268855-120294711 | L1 | ENSG00000144118.14 |
| MSTRG.17445.11 | U251 | chr2:218525701-218553724 | L1 | ENSG00000135913.11 |
| MSTRG.17632.11 | U251 | chr2:238177519-238203708 | L1 | ENSG00000132323.9 |
| MSTRG.17716.6 | U251 | chr2:241735319-241759955 | L1 | ENSG00000180902.18 |
| MSTRG.17946.18 | U251 | chr20:25211997-25228172 | L1 | ENSG00000197586.13 |
| MSTRG.15508.18 | K562 | chr20:63926086-63936031 | L1 | ENSG00000101152.11 |
| MSTRG.15736.1 | K562 | chr21:37221767-37267943 | L1 | ENSG00000157538.14 |
| MSTRG.15760.6 | K562 | chr21:37960247-38121360 | L1 | ENSG00000157542.11 |
| MSTRG.18831.3 | U251 | chr22:19036223-19052578 | L1 | ENSG00000070413.20 |
| MSTRG.15934.8 | K562 | chr22:19039183-19122454 | L1 | ENSG00000070413.20 |
| MSTRG.18994.1 | U251 | chr22:24555958-24577104 | L1 | ENSG00000286070.1 |
| MSTRG.16613.10 | K562 | chr3:23898251-23911453 | L1 | ENSG00000197885.10 |
| MSTRG.17908.2 | K562 | chr4:52799-70174 | L1 | ENSG00000272602.6 |
| MSTRG.18371.6 | K562 | chr4:82903064-82920499 | L1 | ENSG00000168152.13 |
| MSTRG.22914.7 | U251 | chr5:80128163-80205455 | L1 | ENSG00000164300.17 |
| MSTRG.24335.8 | U251 | chr6:33405842-33410153 | L1 | ENSG00000237649.8 |
| MSTRG.20640.5 | K562 | chr6:38722932-38887011 | L1 | ENSG00000124721.18 |
| MSTRG.24547.1 | U251 | chr6:49463356-49477570 | L1 | ENSG00000031691.7 |

| | | | | |
|---|---|---|---|---|
| MSTRG.24547.3 | U251 | chr6:49463357-49477570 | L1 | ENSG00000031691.7 |
| MSTRG.24547.5 | U251 | chr6:49463382-49477570 | L1 | ENSG00000031691.7 |
| MSTRG.22262.12 | K562 | chr7:111533468-111562517 | L1 | ENSG00000184903.9 |
| MSTRG.26698.14 | U251 | chr7:139780454-139796788 | L1 | ENSG00000059377.17 |
| MSTRG.22503.2 | K562 | chr7:142855072-142871094 | L1 | ENSG00000106123.12 |
| MSTRG.26005.16 | U251 | chr7:66682239-66716342 | L1 | ENSG00000154710.17 |
| MSTRG.26032.10 | U251 | chr7:71704934-71713601 | L1 | ENSG00000185274.12 |
| MSTRG.23559.19 | K562 | chr8:144405128-144409421 | L1 | ENSG00000071894.17 |
| MSTRG.27101.9 | U251 | chr8:25459789-25511370 | L1 | ENSG00000184661.14 |
| MSTRG.23716.2 | K562 | chr9:20683978-20747310 | L1 | ENSG00000188352.12 |
| MSTRG.25186.3 | K562 | chrX:124375912-124484655 | L1 | ENSG00000009694.13 |
| MSTRG.24868.1 | K562 | chrX:54530181-54570766 | L1 | ENSG00000130119.16 |
| MSTRG.17969.23 | U251 | chr20:31547446-31572048 | MCTS2P | ENSG00000101294.17 |
| MSTRG.5216.1 | K562 | chr12:32725220-32755897 | NAP1L1P4 | ENSG00000139131.13 |
| MSTRG.5216.2 | K562 | chr12:32725220-32755897 | NAP1L1P4 | ENSG00000139131.13 |
| MSTRG.25232.10 | U251 | chr6:150721046-150830028 | PDCL3P5 | ENSG00000120278.16 |
| MSTRG.25232.11 | U251 | chr6:150721065-150830028 | PDCL3P5 | ENSG00000120278.16 |
| MSTRG.24208.9 | K562 | chr9:112080074-112174724 | RPL29P49 | ENSG00000106868.16 |
| MSTRG.22343.4 | U251 | chr4:186105635-186158843 | RPSAP70 | ENSG00000109794.13 |
| MSTRG.24753.3 | K562 | chrX:44307435-44343672 | RRM2P3 | ENSG00000183690.13 |
| MSTRG.19672.22 | U251 | chr3:37142514-37176059 | UBE2FP1 | ENSG00000093167.18 |
| MSTRG.8404.10 | K562 | chr15:79898840-79918528 | ZNF768P1 | ENSG00000180953.11 |

**Table S3. Chimeric transcripts validation by long-reads**

| Transcript_id | Genomic Position | Retroelement | Gene Coverage | Validated | Is correct ? |
|---|---|---|---|---|---|
| MSTRG.1695.10 | chr1:155199346-155202458 | Alu | <10 reads | N | N |
| MSTRG.4433.11 | chr11:77924073-77994671 | Alu | <10 reads | N | N |
| MSTRG.5833.2 | chr12:101745508-101830959 | Alu | <10 reads | N | N |
| MSTRG.16798.8 | chr3:44840705-44874067 | Alu | <10 reads | N | N |
| MSTRG.19090.4 | chr5:43289395-43313512 | Alu | <10 reads | N | N |
| MSTRG.21102.12 | chr6:118662112-118710075 | Alu | <10 reads | N | N |
| MSTRG.20720.12 | chr6:43557837-43561008 | Alu | <10 reads | N | N |
| MSTRG.17512.6 | chr3:142449238-142553398 | EIF2AK1P1 | <10 reads | N | N |

| | | | | | |
|---|---|---|---|---|---|
| MSTRG.3683.5 | chr11:8612123-8664255 | L1 | <10 reads | N | N |
| MSTRG.8889.20 | chr16:10454764-10483638 | L1 | <10 reads | N | N |
| MSTRG.20640.5 | chr6:38722932-38887011 | L1 | <10 reads | N | N |
| MSTRG.25186.3 | chrX:124375912-124484655 | L1 | <10 reads | N | N |
| MSTRG.14863.10 | chr2:233168174-233178262 | Alu | <10 reads | Y | Y |
| MSTRG.15760.6 | chr21:37960247-38121360 | L1 | <10 reads | Y | N |
| MSTRG.24753.3 | chrX:44307435-44343672 | RRM2P3 | <10 reads | Y | Y |
| MSTRG.356.5 | chr1:20742679-20747101 | Alu | >10 reads | N | N |
| MSTRG.1059.2 | chr1:75202134-75660240 | Alu | >10 reads | N | N |
| MSTRG.2862.12 | chr10:50082430-50133673 | Alu | >10 reads | N | N |
| MSTRG.2930.3 | chr10:68636472-68694487 | Alu | >10 reads | N | N |
| MSTRG.3003.9 | chr10:73442962-73496024 | Alu | >10 reads | N | N |
| MSTRG.3771.6 | chr11:17355455-17380648 | Alu | >10 reads | N | N |
| MSTRG.4115.10 | chr11:64226300-64228266 | Alu | >10 reads | N | N |
| MSTRG.5152.14 | chr12:28225285-28581511 | Alu | >10 reads | N | N |
| MSTRG.9190.6 | chr16:29906963-29926236 | Alu | >10 reads | N | N |
| MSTRG.12806.1 | chr19:43606822-43608939 | Alu | >10 reads | N | N |
| MSTRG.12139.1 | chr19:9641664-9658668 | Alu | >10 reads | N | N |
| MSTRG.14434.9 | chr2:172558454-172629793 | Alu | >10 reads | N | N |
| MSTRG.14018.12 | chr2:96875882-96909141 | Alu | >10 reads | N | N |
| MSTRG.15776.25 | chr21:37154508-37187341 | Alu | >10 reads | N | N |
| MSTRG.17459.2 | chr3:138652699-138776907 | Alu | >10 reads | N | N |
| MSTRG.18615.8 | chr4:127972865-128010808 | Alu | >10 reads | N | N |
| MSTRG.18285.5 | chr4:70902378-70988350 | Alu | >10 reads | N | N |
| MSTRG.20236.1 | chr6:17614793-17688618 | Alu | >10 reads | N | N |
| MSTRG.20114.1 | chr6:2885720-2903309 | Alu | >10 reads | N | N |
| MSTRG.22135.10 | chr7:100853604-100867010 | Alu | >10 reads | N | N |
| MSTRG.22431.1 | chr7:138460217-138589996 | Alu | >10 reads | N | N |
| MSTRG.22431.3 | chr7:138460232-138589996 | Alu | >10 reads | N | N |
| MSTRG.21931.22 | chr7:74752897-74760692 | Alu | >10 reads | N | N |
| MSTRG.24131.5 | chr9:97691929-97697357 | Alu | >10 reads | N | N |
| MSTRG.24962.3 | chrX:78131806-78139679 | Alu | >10 reads | N | N |
| MSTRG.19194.9 | chr5:62332565-62348167 | CRTC2P1 | >10 reads | N | N |
| MSTRG.1140.1 | chr1:88659495-88836255 | ELOCP19 | >10 reads | N | N |
| MSTRG.23395.7 | chr8:123072723-123123682 | HMGB1P19 | >10 reads | N | N |

| | | | | | |
|---|---|---|---|---|---|
| MSTRG.1094.14 | chr1:77955255-77979110 | L1 | >10 reads | N | N |
| MSTRG.3161.2 | chr10:94332763-94345247 | L1 | >10 reads | N | N |
| MSTRG.4602.1 | chr11:108154744-108205417 | L1 | >10 reads | N | N |
| MSTRG.17908.2 | chr4:52799-70174 | L1 | >10 reads | N | N |
| MSTRG.18371.6 | chr4:82903064-82920499 | L1 | >10 reads | N | N |
| MSTRG.23559.19 | chr8:144405128-144409421 | L1 | >10 reads | N | N |
| MSTRG.8404.10 | chr15:79898840-79918528 | ZNF768P1 | >10 reads | N | N |
| MSTRG.476.7 | chr1:27322281-27336595 | ACTG1P20 | >10 reads | Y | N |
| MSTRG.212.3 | chr1:10474950-10520094 | Alu | >10 reads | Y | Y |
| MSTRG.1549.2 | chr1:149054033-149102884 | Alu | >10 reads | Y | Y |
| MSTRG.87.1 | chr1:1784988-1880272 | Alu | >10 reads | Y | N |
| MSTRG.87.9 | chr1:1785286-1880272 | Alu | >10 reads | Y | N |
| MSTRG.2275.12 | chr1:224228973-224330189 | Alu | >10 reads | Y | N |
| MSTRG.589.17 | chr1:32820399-32858879 | Alu | >10 reads | Y | N |
| MSTRG.2744.7 | chr10:42589119-42638154 | Alu | >10 reads | Y | Y |
| MSTRG.2484.11 | chr10:4826730-4848278 | Alu | >10 reads | Y | Y |
| MSTRG.2915.16 | chr10:68489023-68527523 | Alu | >10 reads | Y | Y |
| MSTRG.3087.3 | chr10:87095196-87191468 | Alu | >10 reads | Y | Y |
| MSTRG.3161.1 | chr10:94321955-94362968 | Alu | >10 reads | Y | Y |
| MSTRG.3186.8 | chr10:96130509-96164028 | Alu | >10 reads | Y | Y |
| MSTRG.3771.5 | chr11:17354862-17383674 | Alu | >10 reads | Y | N |
| MSTRG.3520.6 | chr11:651498-694961 | Alu | >10 reads | Y | N |
| MSTRG.6135.1 | chr12:123601537-123620943 | Alu | >10 reads | Y | Y |
| MSTRG.6210.7 | chr12:131723258-131739342 | Alu | >10 reads | Y | N |
| MSTRG.6374.6 | chr13:40943440-41061440 | Alu | >10 reads | Y | Y |
| MSTRG.7250.9 | chr14:70352972-70417090 | Alu | >10 reads | Y | N |
| MSTRG.8354.4 | chr15:77108164-77388615 | Alu | >10 reads | Y | Y |
| MSTRG.8354.3 | chr15:77108164-77420210 | Alu | >10 reads | Y | Y |
| MSTRG.8701.4 | chr16:1833973-1872178 | Alu | >10 reads | Y | Y |
| MSTRG.8701.6 | chr16:1833983-1872178 | Alu | >10 reads | Y | Y |
| MSTRG.9357.14 | chr16:47593327-47701523 | Alu | >10 reads | Y | N |
| MSTRG.10552.11 | chr17:36500895-36535251 | Alu | >10 reads | Y | Y |
| MSTRG.11595.12 | chr18:23529596-23531211 | Alu | >10 reads | Y | N |
| MSTRG.11729.5 | chr18:50257583-50281574 | Alu | >10 reads | Y | N |
| MSTRG.12390.6 | chr19:17331522-17334855 | Alu | >10 reads | Y | Y |

| | | | | | |
|---|---|---|---|---|---|
| MSTRG.13176.13 | chr19:55158663-55166890 | Alu | >10 reads | Y | N |
| MSTRG.11865.8 | chr19:959130-975939 | Alu | >10 reads | Y | Y |
| MSTRG.13439.1 | chr2:26242781-26290465 | Alu | >10 reads | Y | Y |
| MSTRG.15629.8 | chr21:17596982-17606084 | Alu | >10 reads | Y | N |
| MSTRG.15762.3 | chr21:38121473-38172513 | Alu | >10 reads | Y | Y |
| MSTRG.17833.3 | chr3:195274745-195442307 | Alu | >10 reads | Y | Y |
| MSTRG.16676.1 | chr3:33388336-33416834 | Alu | >10 reads | Y | N |
| MSTRG.18804.4 | chr4:173232452-173323967 | Alu | >10 reads | Y | Y |
| MSTRG.19570.7 | chr5:116003702-116027619 | Alu | >10 reads | Y | Y |
| MSTRG.19829.9 | chr5:141969105-142011162 | Alu | >10 reads | Y | Y |
| MSTRG.19030.1 | chr5:34897057-34915626 | Alu | >10 reads | Y | Y |
| MSTRG.19030.2 | chr5:34897068-34915626 | Alu | >10 reads | Y | Y |
| MSTRG.19030.3 | chr5:34897068-34915626 | Alu | >10 reads | Y | Y |
| MSTRG.19103.5 | chr5:43491717-43515148 | Alu | >10 reads | Y | Y |
| MSTRG.19339.13 | chr5:79614121-79623489 | Alu | >10 reads | Y | Y |
| MSTRG.21087.3 | chr6:117689642-117710813 | Alu | >10 reads | Y | Y |
| MSTRG.20754.7 | chr6:52281323-52284881 | Alu | >10 reads | Y | Y |
| MSTRG.21826.2 | chr7:56010484-56051604 | Alu | >10 reads | Y | Y |
| MSTRG.22056.1 | chr7:96120220-96309684 | Alu | >10 reads | Y | N |
| MSTRG.23306.1 | chr8:101686358-101988056 | Alu | >10 reads | Y | Y |
| MSTRG.22876.1 | chr8:30578182-30653983 | Alu | >10 reads | Y | Y |
| MSTRG.24212.1 | chr9:112683926-112718149 | Alu | >10 reads | Y | Y |
| MSTRG.24317.15 | chr9:125052214-125143506 | Alu | >10 reads | Y | Y |
| MSTRG.23818.3 | chr9:35782054-35809732 | Alu | >10 reads | Y | Y |
| MSTRG.23846.8 | chr9:36572882-36677683 | Alu | >10 reads | Y | N |
| MSTRG.24016.3 | chr9:88388444-88478694 | Alu | >10 reads | Y | N |
| MSTRG.1478.4 | chr1:147928436-147980088 | L1 | >10 reads | Y | Y |
| MSTRG.2409.1 | chr1:243256034-243318523 | L1 | >10 reads | Y | Y |
| MSTRG.2409.5 | chr1:243256053-243318523 | L1 | >10 reads | Y | Y |
| MSTRG.2409.6 | chr1:243256053-243318523 | L1 | >10 reads | Y | Y |
| MSTRG.1084.9 | chr1:77779853-77789133 | L1 | >10 reads | Y | N |
| MSTRG.5222.3 | chr12:39626184-39693491 | L1 | >10 reads | Y | Y |
| MSTRG.7960.14 | chr15:49622292-49637228 | L1 | >10 reads | Y | Y |
| MSTRG.11560.12 | chr18:14119916-14132490 | L1 | >10 reads | Y | N |
| MSTRG.11534.1 | chr18:9914002-9960120 | L1 | >10 reads | Y | Y |

| | | | | | |
|---|---|---|---|---|---|
| MSTRG.11534.8 | chr18:9914658-9960120 | L1 | >10 reads | Y | Y |
| MSTRG.12521.9 | chr19:23658965-23687220 | L1 | >10 reads | Y | N |
| MSTRG.15508.18 | chr20:63926086-63936031 | L1 | >10 reads | Y | Y |
| MSTRG.15736.1 | chr21:37221767-37267943 | L1 | >10 reads | Y | N |
| MSTRG.15934.8 | chr22:19039183-19122454 | L1 | >10 reads | Y | Y |
| MSTRG.16613.10 | chr3:23898251-23911453 | L1 | >10 reads | Y | N |
| MSTRG.22262.12 | chr7:111533468-111562517 | L1 | >10 reads | Y | Y |
| MSTRG.22503.2 | chr7:142855072-142871094 | L1 | >10 reads | Y | Y |
| MSTRG.23716.2 | chr9:20683978-20747310 | L1 | >10 reads | Y | Y |
| MSTRG.24868.1 | chrX:54530181-54570766 | L1 | >10 reads | Y | N |
| MSTRG.5216.1 | chr12:32725220-32755897 | NAP1L1P4 | >10 reads | Y | N |
| MSTRG.5216.2 | chr12:32725220-32755897 | NAP1L1P4 | >10 reads | Y | N |
| MSTRG.24208.9 | chr9:112080074-112174724 | RPL29P49 | >10 reads | Y | N |

**Table S4. Protein analisys (coding or not) of the chimeric transcripts**

| Transcript_id | Cell Line | Strand in relation to the host | Retroelement | Event Position | coding (?) | Alter Domain |
|---|---|---|---|---|---|---|
| MSTRG.11595.12 | K562 | Opposite strand | Alu | Final | coding | Mantain |
| MSTRG.11729.5 | K562 | Opposite strand | Alu | Final | coding | Mantain |
| MSTRG.12390.6 | K562 | Same strand | Alu | Final | coding | Delete |
| MSTRG.14863.10 | K562 | Same strand | Alu | Final | noncoding | - |
| MSTRG.1695.10 | K562 | Opposite strand | Alu | Final | noncoding | - |
| MSTRG.19030.1 | K562 | Opposite strand | Alu | Final | coding | Mantain |
| MSTRG.19030.2 | K562 | Opposite strand | Alu | Final | coding | Mantain |
| MSTRG.19030.3 | K562 | Opposite strand | Alu | Final | coding | Mantain |
| MSTRG.19103.5 | K562 | Opposite strand | Alu | Final | coding | Delete |
| MSTRG.19829.9 | K562 | Opposite strand | Alu | Final | coding | Mantain |

| | | | | | | |
|---|---|---|---|---|---|---|
| MSTRG.20720.12 | K562 | Opposite strand | Alu | Final | noncoding | - |
| MSTRG.20754.7 | K562 | Opposite strand | Alu | Final | coding | Delete |
| MSTRG.21102.12 | K562 | Opposite strand | Alu | Final | noncoding | - |
| MSTRG.212.3 | K562 | Opposite strand | Alu | Final | noncoding | - |
| MSTRG.2275.12 | K562 | Opposite strand | Alu | Final | coding | Delete |
| MSTRG.24131.5 | K562 | Opposite strand | Alu | Final | coding | Delete |
| MSTRG.24317.15 | K562 | Opposite strand | Alu | Final | coding | Delete |
| MSTRG.24962.3 | K562 | Same strand | Alu | Final | coding | Mantain |
| MSTRG.2915.16 | K562 | Opposite strand | Alu | Final | coding | Mantain |
| MSTRG.3161.1 | K562 | Opposite strand | Alu | Final | coding | Mantain |
| MSTRG.3520.6 | K562 | Opposite strand | Alu | Final | coding | Delete |
| MSTRG.4115.10 | K562 | Opposite strand | Alu | Final | coding | Mantain |
| MSTRG.4433.11 | K562 | Opposite strand | Alu | Final | coding | Mantain |
| MSTRG.6210.7 | K562 | Opposite strand | Alu | Final | coding | Delete |
| MSTRG.7250.9 | K562 | Opposite strand | Alu | Final | noncoding | - |
| MSTRG.17512.6 | K562 | Same strand | EIF2AK1P1 | Final | coding | Delete |
| MSTRG.23395.7 | K562 | Same strand | HMGB1P19 | Final | coding | Delete |
| MSTRG.1084.9 | K562 | Opposite strand | L1 | Final | noncoding | - |
| MSTRG.1094.14 | K562 | Opposite strand | L1 | Final | coding | Delete |
| MSTRG.11560.12 | K562 | Opposite strand | L1 | Final | noncoding | - |
| MSTRG.12521.9 | K562 | Opposite strand | L1 | Final | noncoding | - |

| | | | | | | |
|---|---|---|---|---|---|---|
| MSTRG.1478.4 | K562 | Opposite strand | L1 | Final | noncoding | - |
| MSTRG.15736.1 | K562 | Same strand | L1 | Final | coding | Mantain |
| MSTRG.15760.6 | K562 | Opposite strand | L1 | Final | noncoding | - |
| MSTRG.16613.10 | K562 | Opposite strand | L1 | Final | coding | Delete |
| MSTRG.17908.2 | K562 | Opposite strand | L1 | Final | noncoding | - |
| MSTRG.22262.12 | K562 | Opposite strand | L1 | Final | noncoding | - |
| MSTRG.23559.19 | K562 | Opposite strand | L1 | Final | coding | Delete |
| MSTRG.23716.2 | K562 | Opposite strand | L1 | Final | coding | Delete |
| MSTRG.2409.1 | K562 | Same strand | L1 | Final | coding | Mantain |
| MSTRG.2409.5 | K562 | Same strand | L1 | Final | coding | Mantain |
| MSTRG.2409.6 | K562 | Same strand | L1 | Final | coding | Mantain |
| MSTRG.24868.1 | K562 | Opposite strand | L1 | Final | coding | Mantain |
| MSTRG.5222.3 | K562 | Opposite strand | L1 | Final | coding | Mantain |
| MSTRG.7960.14 | K562 | Opposite strand | L1 | Final | coding | Mantain |
| MSTRG.5216.1 | K562 | Same strand | NAP1L1P4 | Final | coding | Mantain |
| MSTRG.5216.2 | K562 | Same strand | NAP1L1P4 | Final | coding | Mantain |
| MSTRG.24753.3 | K562 | Same strand | RRM2P3 | Final | coding | Change |
| MSTRG.11681.3 | U251 | Opposite strand | Alu | Final | coding | Delete |
| MSTRG.12399.5 | U251 | Opposite strand | Alu | Final | coding | Mantain |
| MSTRG.1430.6 | U251 | Opposite strand | Alu | Final | coding | Mantain |
| MSTRG.1462.18 | U251 | Opposite strand | Alu | Final | coding | Mantain |
| MSTRG.16479.6 | U251 | Opposite strand | Alu | Final | coding | Mantain |
| MSTRG.18253.2 | U251 | Same strand | Alu | Final | coding | Mantain |

| | | | | | | |
|---|---|---|---|---|---|---|
| MSTRG.19236.5 | U251 | Same strand | Alu | Final | coding | Mantain |
| MSTRG.19893.1 | U251 | Opposite strand | Alu | Final | coding | Mantain |
| MSTRG.19984.15 | U251 | Same strand | Alu | Final | noncoding | - |
| MSTRG.20588.5 | U251 | Same strand | Alu | Final | coding | Delete |
| MSTRG.20746.1 | U251 | Opposite strand | Alu | Final | coding | Mantain |
| MSTRG.21785.2 | U251 | Same strand | Alu | Final | coding | Delete |
| MSTRG.22982.9 | U251 | Opposite strand | Alu | Final | coding | Delete |
| MSTRG.23451.8 | U251 | Opposite strand | Alu | Final | noncoding | - |
| MSTRG.24152.3 | U251 | Opposite strand | Alu | Final | coding | Mantain |
| MSTRG.25864.9 | U251 | Opposite strand | Alu | Final | coding | Delete |
| MSTRG.28153.4 | U251 | Same strand | Alu | Final | coding | Mantain |
| MSTRG.5968.2 | U251 | Same strand | Alu | Final | coding | Mantain |
| MSTRG.6232.7 | U251 | Opposite strand | Alu | Final | coding | Mantain |
| MSTRG.6863.13 | U251 | Opposite strand | Alu | Final | coding | Delete |
| MSTRG.6870.5 | U251 | Opposite strand | Alu | Final | noncoding | - |
| MSTRG.7682.8 | U251 | Opposite strand | Alu | Final | coding | Mantain |
| MSTRG.9313.2 | U251 | Opposite strand | Alu | Final | coding | Mantain |
| MSTRG.11388.1 | U251 | Opposite strand | L1 | Final | coding | Delete |
| MSTRG.2082.1 | U251 | Opposite strand | L1 | Final | coding | Mantain |
| MSTRG.21619.5 | U251 | Same strand | L1 | Final | coding | Delete |
| MSTRG.23216.1 | U251 | Same strand | L1 | Final | noncoding | - |
| MSTRG.23216.3 | U251 | Same strand | L1 | Final | noncoding | - |

| | | | | | | |
|---|---|---|---|---|---|---|
| MSTRG.24518.14 | U251 | Opposite strand | L1 | Final | noncoding | - |
| MSTRG.25190.12 | U251 | Same strand | L1 | Final | noncoding | - |
| MSTRG.2705.4 | U251 | Same strand | L1 | Final | coding | Mantain |
| MSTRG.4394.4 | U251 | Opposite strand | L1 | Final | noncoding | - |
| MSTRG.6294.2 | U251 | Opposite strand | L1 | Final | coding | Delete |
| MSTRG.6866.7 | U251 | Opposite strand | L1 | Final | coding | Delete |
| MSTRG.23868.9 | U251 | Same strand | PDCL3P5 | Final | coding | Mantain |
| MSTRG.1059.2 | K562 | Opposite strand | Alu | Initial | coding | Delete |
| MSTRG.11865.8 | K562 | Opposite strand | Alu | Initial | coding | Mantain |
| MSTRG.12139.1 | K562 | Same strand | Alu | Initial | coding | Delete |
| MSTRG.12806.1 | K562 | Opposite strand | Alu | Initial | noncoding | - |
| MSTRG.13439.1 | K562 | Opposite strand | Alu | Initial | coding | Mantain |
| MSTRG.14018.12 | K562 | Opposite strand | Alu | Initial | coding | Delete |
| MSTRG.15762.3 | K562 | Opposite strand | Alu | Initial | noncoding | - |
| MSTRG.15776.25 | K562 | Opposite strand | Alu | Initial | coding | Delete |
| MSTRG.16798.8 | K562 | Opposite strand | Alu | Initial | coding | Delete |
| MSTRG.17459.2 | K562 | Opposite strand | Alu | Initial | coding | Mantain |
| MSTRG.18615.8 | K562 | Opposite strand | Alu | Initial | noncoding | - |
| MSTRG.18804.4 | K562 | Opposite strand | Alu | Initial | coding | Mantain |
| MSTRG.19339.13 | K562 | Opposite strand | Alu | Initial | coding | Delete |
| MSTRG.19570.7 | K562 | Opposite strand | Alu | Initial | noncoding | - |
| MSTRG.21087.3 | K562 | Opposite strand | Alu | Initial | coding | Delete |

| | | | | | | |
|---|---|---|---|---|---|---|
| MSTRG.21931.22 | K562 | Same strand | Alu | Initial | noncoding | - |
| MSTRG.22135.10 | K562 | Opposite strand | Alu | Initial | coding | Mantain |
| MSTRG.22876.1 | K562 | Opposite strand | Alu | Initial | coding | Mantain |
| MSTRG.23306.1 | K562 | Opposite strand | Alu | Initial | coding | Mantain |
| MSTRG.23818.3 | K562 | Opposite strand | Alu | Initial | coding | Delete |
| MSTRG.2484.11 | K562 | Same strand | Alu | Initial | coding | Mantain |
| MSTRG.2744.7 | K562 | Same strand | Alu | Initial | coding | Delete |
| MSTRG.2862.12 | K562 | Opposite strand | Alu | Initial | coding | Mantain |
| MSTRG.2930.3 | K562 | Same strand | Alu | Initial | coding | Delete |
| MSTRG.356.5 | K562 | Opposite strand | Alu | Initial | noncoding | - |
| MSTRG.5152.14 | K562 | Opposite strand | Alu | Initial | coding | Mantain |
| MSTRG.87.1 | K562 | Opposite strand | Alu | Initial | coding | Mantain |
| MSTRG.87.9 | K562 | Opposite strand | Alu | Initial | coding | Mantain |
| MSTRG.9357.14 | K562 | Same strand | Alu | Initial | coding | Mantain |
| MSTRG.19194.9 | K562 | Same strand | CRTC2P1 | Initial | noncoding | - |
| MSTRG.15508.18 | K562 | Opposite strand | L1 | Initial | coding | Mantain |
| MSTRG.18371.6 | K562 | Opposite strand | L1 | Initial | coding | Delete |
| MSTRG.25186.3 | K562 | Same strand | L1 | Initial | coding | Delete |
| MSTRG.3161.2 | K562 | Opposite strand | L1 | Initial | coding | Delete |
| MSTRG.3683.5 | K562 | Opposite strand | L1 | Initial | coding | Mantain |
| MSTRG.4602.1 | K562 | Opposite strand | L1 | Initial | coding | Mantain |
| MSTRG.8889.20 | K562 | Opposite strand | L1 | Initial | coding | Mantain |
| MSTRG.24208.9 | K562 | Same strand | RPL29P49 | Initial | coding | Delete |

| | | | | | | |
|---|---|---|---|---|---|---|
| MSTRG.8404.10 | K562 | Same strand | ZNF768P1 | Initial | noncoding | - |
| MSTRG.10527.19 | U251 | Same strand | Alu | Initial | coding | Delete |
| MSTRG.12578.3 | U251 | Opposite strand | Alu | Initial | noncoding | - |
| MSTRG.14185.2 | U251 | Opposite strand | Alu | Initial | coding | Mantain |
| MSTRG.17715.2 | U251 | Opposite strand | Alu | Initial | noncoding | - |
| MSTRG.18176.7 | U251 | Same strand | Alu | Initial | coding | Mantain |
| MSTRG.19830.19 | U251 | Same strand | Alu | Initial | coding | Mantain |
| MSTRG.22884.5 | U251 | Same strand | Alu | Initial | noncoding | - |
| MSTRG.23002.3 | U251 | Opposite strand | Alu | Initial | coding | Delete |
| MSTRG.24493.6 | U251 | Opposite strand | Alu | Initial | coding | Mantain |
| MSTRG.25756.10 | U251 | Same strand | Alu | Initial | coding | Mantain |
| MSTRG.26294.6 | U251 | Opposite strand | Alu | Initial | coding | Delete |
| MSTRG.26371.5 | U251 | Same strand | Alu | Initial | coding | Mantain |
| MSTRG.26902.1 | U251 | Opposite strand | Alu | Initial | coding | Mantain |
| MSTRG.312.2 | U251 | Opposite strand | Alu | Initial | coding | Mantain |
| MSTRG.4116.13 | U251 | Opposite strand | Alu | Initial | coding | Delete |
| MSTRG.44.10 | U251 | Same strand | Alu | Initial | coding | Delete |
| MSTRG.478.6 | U251 | Opposite strand | Alu | Initial | coding | Mantain |
| MSTRG.5420.20 | U251 | Opposite strand | Alu | Initial | coding | Mantain |
| MSTRG.596.9 | U251 | Opposite strand | Alu | Initial | coding | Mantain |
| MSTRG.6073.5 | U251 | Opposite strand | Alu | Initial | coding | Mantain |
| MSTRG.8511.7 | U251 | Opposite strand | Alu | Initial | coding | Mantain |

| | | | | | | |
|---|---|---|---|---|---|---|
| MSTRG.11327.1 | U251 | Opposite strand | L1 | Initial | coding | Mantain |
| MSTRG.13746.5 | U251 | Same strand | L1 | Initial | coding | Delete |
| MSTRG.15763.11 | U251 | Opposite strand | L1 | Initial | coding | Mantain |
| MSTRG.1594.6 | U251 | Same strand | L1 | Initial | coding | Mantain |
| MSTRG.16922.16 | U251 | Opposite strand | L1 | Initial | coding | Mantain |
| MSTRG.17762.1 | U251 | Opposite strand | L1 | Initial | coding | Delete |
| MSTRG.18280.3 | U251 | Same strand | L1 | Initial | coding | Delete |
| MSTRG.24993.2 | U251 | Opposite strand | L1 | Initial | coding | Mantain |
| MSTRG.3126.4 | U251 | Same strand | L1 | Initial | coding | Delete |
| MSTRG.6866.6 | U251 | Opposite strand | L1 | Initial | coding | Delete |
| MSTRG.9308.4 | U251 | Same strand | L1 | Initial | noncoding | - |
| MSTRG.476.7 | K562 | Same strand | ACTG1P20 | Internal | noncoding | - |
| MSTRG.10552.11 | K562 | Opposite strand | Alu | Internal | coding | Mantain |
| MSTRG.13176.13 | K562 | Same strand | Alu | Internal | noncoding | - |
| MSTRG.14434.9 | K562 | Opposite strand | Alu | Internal | noncoding | - |
| MSTRG.1549.2 | K562 | Opposite strand | Alu | Internal | coding | Mantain |
| MSTRG.15629.8 | K562 | Opposite strand | Alu | Internal | noncoding | - |
| MSTRG.16676.1 | K562 | Opposite strand | Alu | Internal | coding | Delete |
| MSTRG.17833.3 | K562 | Opposite strand | Alu | Internal | coding | Delete |
| MSTRG.18285.5 | K562 | Opposite strand | Alu | Internal | coding | Mantain |
| MSTRG.19090.4 | K562 | Opposite strand | Alu | Internal | coding | Delete |
| MSTRG.20114.1 | K562 | Opposite strand | Alu | Internal | coding | Mantain |

| | | | | | | |
|---|---|---|---|---|---|---|
| MSTRG.20236.1 | K562 | Opposite strand | Alu | Internal | coding | Mantain |
| MSTRG.21826.2 | K562 | Opposite strand | Alu | Internal | coding | Mantain |
| MSTRG.22056.1 | K562 | Opposite strand | Alu | Internal | coding | Delete |
| MSTRG.22431.1 | K562 | Opposite strand | Alu | Internal | coding | Mantain |
| MSTRG.22431.3 | K562 | Opposite strand | Alu | Internal | coding | Mantain |
| MSTRG.23846.8 | K562 | Opposite strand | Alu | Internal | coding | Mantain |
| MSTRG.24016.3 | K562 | Opposite strand | Alu | Internal | coding | Mantain |
| MSTRG.24212.1 | K562 | Opposite strand | Alu | Internal | noncoding | - |
| MSTRG.3003.9 | K562 | Opposite strand | Alu | Internal | coding | Mantain |
| MSTRG.3087.3 | K562 | Opposite strand | Alu | Internal | coding | Mantain |
| MSTRG.3186.8 | K562 | Opposite strand | Alu | Internal | coding | Delete |
| MSTRG.3771.5 | K562 | Opposite strand | Alu | Internal | coding | Mantain |
| MSTRG.3771.6 | K562 | Opposite strand | Alu | Internal | noncoding | - |
| MSTRG.5833.2 | K562 | Opposite strand | Alu | Internal | coding | Delete |
| MSTRG.589.17 | K562 | Same strand | Alu | Internal | coding | Mantain |
| MSTRG.6135.1 | K562 | Opposite strand | Alu | Internal | coding | Mantain |
| MSTRG.6374.6 | K562 | Opposite strand | Alu | Internal | coding | Delete |
| MSTRG.8354.3 | K562 | Opposite strand | Alu | Internal | coding | Mantain |
| MSTRG.8354.4 | K562 | Opposite strand | Alu | Internal | coding | Mantain |
| MSTRG.8701.4 | K562 | Same strand | Alu | Internal | coding | Mantain |
| MSTRG.8701.6 | K562 | Same strand | Alu | Internal | coding | Delete |
| MSTRG.9190.6 | K562 | Opposite strand | Alu | Internal | coding | Mantain |
| MSTRG.1140.1 | K562 | Same strand | ELOCP19 | Internal | coding | Delete |
| MSTRG.11534.1 | K562 | Same strand | L1 | Internal | coding | Delete |
| MSTRG.11534.8 | K562 | Same strand | L1 | Internal | coding | Delete |
| MSTRG.15934.8 | K562 | Opposite strand | L1 | Internal | coding | Mantain |

| | | | | | | |
|---|---|---|---|---|---|---|
| MSTRG.20640.5 | K562 | Opposite strand | L1 | Internal | noncoding | - |
| MSTRG.22503.2 | K562 | Opposite strand | L1 | Internal | coding | Mantain |
| MSTRG.1594.8 | U251 | Opposite strand | Alu | Internal | coding | Mantain |
| MSTRG.1675.11 | U251 | Opposite strand | Alu | Internal | coding | Mantain |
| MSTRG.1675.12 | U251 | Opposite strand | Alu | Internal | coding | Mantain |
| MSTRG.1675.14 | U251 | Opposite strand | Alu | Internal | noncoding | - |
| MSTRG.1675.7 | U251 | Opposite strand | Alu | Internal | coding | Mantain |
| MSTRG.17010.17 | U251 | Same strand | Alu | Internal | noncoding | - |
| MSTRG.21991.3 | U251 | Opposite strand | Alu | Internal | coding | Delete |
| MSTRG.23453.2 | U251 | Opposite strand | Alu | Internal | coding | Delete |
| MSTRG.2541.4 | U251 | Opposite strand | Alu | Internal | coding | Mantain |
| MSTRG.5420.17 | U251 | Opposite strand | Alu | Internal | coding | Delete |
| MSTRG.6133.1 | U251 | Opposite strand | Alu | Internal | noncoding | - |
| MSTRG.8274.5 | U251 | Same strand | Alu | Internal | noncoding | - |
| MSTRG.8296.35 | U251 | Opposite strand | Alu | Internal | noncoding | - |
| MSTRG.9377.10 | U251 | Opposite strand | Alu | Internal | coding | Delete |
| MSTRG.22985.7 | U251 | Opposite strand | L1 | Internal | coding | Mantain |
| MSTRG.22985.8 | U251 | Opposite strand | L1 | Internal | noncoding | - |

**Table S5. Expression in TPM of all LINE1 chimeric transcripts in K562 and U251**

| K562 | | |
|---|---|---|
| transcript_id | K562_short_1 | K562_short_2 |
| MSTRG.10552.11 | 2.085931 | 2.923798 |

| | | |
|---|---|---|
| MSTRG.1059.2 | 7.469551 | 3.09976 |
| MSTRG.1084.9 | 2.407651 | 1.665078 |
| MSTRG.1094.14 | 9.863122 | 13.671737 |
| MSTRG.1140.1 | 3.305025 | 3.455124 |
| MSTRG.11534.1 | 0.01805 | 22.293026 |
| MSTRG.11534.8 | 6.876545 | 0.532243 |
| MSTRG.11560.12 | 3.681681 | 3.401977 |
| MSTRG.11595.12 | 1.335276 | 0.539804 |
| MSTRG.11729.5 | 0.213747 | 0.996922 |
| MSTRG.11865.8 | 1.700158 | 3.97897 |
| MSTRG.12139.1 | 7.420537 | 6.866727 |
| MSTRG.12390.6 | 0.695789 | 0 |
| MSTRG.12521.9 | 2.757259 | 2.924256 |
| MSTRG.12806.1 | 2.010518 | 2.095571 |
| MSTRG.13176.13 | 7.943563 | 4.28587 |
| MSTRG.13439.1 | 1.946087 | 1.52611 |
| MSTRG.14018.12 | 0.48812 | 9.157183 |
| MSTRG.14434.9 | 2.761517 | 3.64908 |
| MSTRG.1478.4 | 1.859089 | 2.918202 |
| MSTRG.14863.10 | 4.065928 | 0.432234 |
| MSTRG.1549.2 | 3.207266 | 3.882085 |
| MSTRG.15508.18 | 0 | 0.980339 |
| MSTRG.15629.8 | 4.023325 | 3.543316 |
| MSTRG.15736.1 | 16.306488 | 2.294404 |
| MSTRG.15760.6 | 3.667135 | 5.096694 |
| MSTRG.15762.3 | 7.376226 | 7.180696 |
| MSTRG.15776.25 | 1.64135 | 3.318321 |
| MSTRG.15934.8 | 4.231621 | 8.207541 |
| MSTRG.16613.10 | 3.404503 | 2.407475 |
| MSTRG.16676.1 | 2.535638 | 1.909387 |
| MSTRG.16798.8 | 2.506922 | 2.530775 |
| MSTRG.1695.10 | 2.78107 | 2.704511 |
| MSTRG.17459.2 | 10.16993 | 0.089357 |
| MSTRG.17512.6 | 2.158638 | 0 |
| MSTRG.17833.3 | 2.279561 | 0.140878 |

| | | |
|---|---|---|
| MSTRG.17908.2 | 14.292798 | 1.835099 |
| MSTRG.18285.5 | 6.293825 | 0.133109 |
| MSTRG.18371.6 | 4.961533 | 3.137846 |
| MSTRG.18615.8 | 2.621167 | 2.504836 |
| MSTRG.18804.4 | 6.617114 | 0.072841 |
| MSTRG.19030.1 | 3.439645 | 3.050813 |
| MSTRG.19030.2 | 2.094336 | 2.804544 |
| MSTRG.19030.3 | 2.889627 | 2.828544 |
| MSTRG.19090.4 | 46.142281 | 63.09119 |
| MSTRG.19103.5 | 2.426028 | 0.227454 |
| MSTRG.19194.9 | 1.722361 | 1.820515 |
| MSTRG.19339.13 | 8.165753 | 9.864581 |
| MSTRG.19570.7 | 3.391373 | 0 |
| MSTRG.19829.9 | 0.115187 | 25.108889 |
| MSTRG.20114.1 | 51.82103 | 14.186197 |
| MSTRG.20236.1 | 105.39959 | 90.807434 |
| MSTRG.20640.5 | 5.605869 | 3.753301 |
| MSTRG.20720.12 | 1.286245 | 1.546923 |
| MSTRG.20754.7 | 5.83212 | 4.28317 |
| MSTRG.21087.3 | 0.146298 | 8.062831 |
| MSTRG.21102.12 | 2.643301 | 1.662246 |
| MSTRG.212.3 | 2.423633 | 2.945162 |
| MSTRG.21826.2 | 5.442931 | 10.111135 |
| MSTRG.21931.22 | 1.465589 | 0 |
| MSTRG.22056.1 | 2.091484 | 0.182324 |
| MSTRG.22135.10 | 4.191368 | 2.854448 |
| MSTRG.22262.12 | 4.539807 | 2.857865 |
| MSTRG.22431.1 | 7.950953 | 10.872099 |
| MSTRG.22431.3 | 7.414785 | 12.155231 |
| MSTRG.22503.2 | 18.405704 | 12.189852 |
| MSTRG.2275.12 | 2.19809 | 12.465183 |
| MSTRG.22876.1 | 20.721643 | 11.709185 |
| MSTRG.23306.1 | 5.190739 | 2.369923 |
| MSTRG.23559.19 | 4.423557 | 4.618327 |
| MSTRG.23716.2 | 2.909037 | 1.914127 |

| | | |
|---|---|---|
| MSTRG.23818.3 | 5.896229 | 6.45274 |
| MSTRG.23846.8 | 0.619048 | 3.709855 |
| MSTRG.24016.3 | 0.444764 | 8.046425 |
| MSTRG.2409.1 | 3.434726 | 9.691006 |
| MSTRG.2409.5 | 4.092635 | 2.978881 |
| MSTRG.2409.6 | 2.8924 | 1.54951 |
| MSTRG.24131.5 | 0.61222 | 1.831424 |
| MSTRG.24208.9 | 3.289324 | 3.155408 |
| MSTRG.24212.1 | 4.528104 | 0.410088 |
| MSTRG.24317.15 | 2.750537 | 2.816752 |
| MSTRG.24753.3 | 3.696918 | 5.39002 |
| MSTRG.2484.11 | 2.918374 | 3.47045 |
| MSTRG.24868.1 | 2.629567 | 4.434083 |
| MSTRG.24962.3 | 2.867173 | 3.921893 |
| MSTRG.25186.3 | 0.173705 | 6.792564 |
| MSTRG.2744.7 | 13.706056 | 15.451977 |
| MSTRG.2862.12 | 9.197124 | 0 |
| MSTRG.2915.16 | 0.153254 | 0.439733 |
| MSTRG.2930.3 | 0.036037 | 2.833518 |
| MSTRG.3003.9 | 2.777372 | 2.157914 |
| MSTRG.3087.3 | 7.885547 | 2.621609 |
| MSTRG.3161.1 | 5.198391 | 4.039767 |
| MSTRG.3161.2 | 0.557238 | 2.675981 |
| MSTRG.3186.8 | 3.388209 | 2.752197 |
| MSTRG.3520.6 | 3.546123 | 3.340207 |
| MSTRG.356.5 | 1.6218 | 0.303111 |
| MSTRG.3683.5 | 1.277661 | 5.979538 |
| MSTRG.3771.5 | 0.369901 | 4.361741 |
| MSTRG.3771.6 | 6.018212 | 0.026118 |
| MSTRG.4115.10 | 0.621606 | 7.452711 |
| MSTRG.4433.11 | 3.506747 | 5.834335 |
| MSTRG.4602.1 | 0.42391 | 11.237644 |
| MSTRG.476.7 | 4.755086 | 7.539613 |
| MSTRG.5152.14 | 0.7371 | 0.437354 |
| MSTRG.5216.1 | 11.140984 | 5.345765 |

| | | | |
|---|---|---|---|
| MSTRG.5216.2 | 4.674569 | 0 | |
| MSTRG.5222.3 | 1.526878 | 1.759319 | |
| MSTRG.5833.2 | 0.250011 | 6.453999 | |
| MSTRG.589.17 | 6.226369 | 0.022713 | |
| MSTRG.6135.1 | 2.142651 | 4.017758 | |
| MSTRG.6210.7 | 1.244806 | 1.9008 | |
| MSTRG.6374.6 | 0.496033 | 1.802167 | |
| MSTRG.7250.9 | 2.711342 | 2.84324 | |
| MSTRG.7960.14 | 0.876138 | 0.129373 | |
| MSTRG.8354.3 | 2.614027 | 0.709114 | |
| MSTRG.8354.4 | 2.833213 | 0 | |
| MSTRG.8404.10 | 2.950806 | 1.31797 | |
| MSTRG.87.1 | 0 | 9.800421 | |
| MSTRG.87.9 | 7.715435 | 12.417842 | |
| MSTRG.8701.4 | 6.279344 | 3.161952 | |
| MSTRG.8701.6 | 5.323295 | 15.158589 | |
| MSTRG.8889.20 | 0.576869 | 0 | |
| MSTRG.9190.6 | 0 | 0.764207 | |
| MSTRG.9357.14 | 0.93387 | 0.372007 | |
| MSTRG.23395.7 | 0 | 0.506135 | |
| U251 | | | |
| transcript_id | U251_1 | U251_2 | U251_3 |
| MSTRG.10527.19 | 0 | 8.129121 | 0 |
| MSTRG.11327.1 | 0.423966 | 0.874161 | 0.165443 |
| MSTRG.11388.1 | 3.647958 | 4.734281 | 3.754154 |
| MSTRG.11681.3 | 0.038411 | 1.688781 | 0.025505 |
| MSTRG.12399.5 | 3.739531 | 1.198316 | 0.617002 |
| MSTRG.12578.3 | 12.35131 | 0 | 0 |
| MSTRG.13746.5 | 2.58286 | 1.334667 | 6.044755 |
| MSTRG.14185.2 | 3.624097 | 0 | 2.232081 |
| MSTRG.1430.6 | 0.436352 | 1.317039 | 5.384473 |
| MSTRG.1462.18 | 0 | 2.129998 | 0 |
| MSTRG.15763.11 | 0 | 5.292066 | 0 |
| MSTRG.1594.6 | 1.494549 | 2.943663 | 9.679945 |
| MSTRG.1594.8 | 11.364431 | 3.448583 | 4.875589 |

| | | | |
|---|---|---|---|
| MSTRG.16479.6 | 0 | 0.394476 | 2.623051 |
| MSTRG.1675.11 | 1.600286 | 0 | 0 |
| MSTRG.1675.12 | 0.119799 | 0.347687 | 0.02944 |
| MSTRG.1675.14 | 4.597611 | 3.784731 | 3.419797 |
| MSTRG.1675.7 | 0.069002 | 2.261529 | 7.365218 |
| MSTRG.16922.16 | 0 | 5.271724 | 1.866353 |
| MSTRG.17010.17 | 11.759724 | 11.68181 | 6.504347 |
| MSTRG.17715.2 | 0.475208 | 5.448405 | 1.90784 |
| MSTRG.17762.1 | 0 | 4.165421 | 13.431605 |
| MSTRG.18176.7 | 1.278197 | 47.199188 | 20.000854 |
| MSTRG.18253.2 | 2.0035 | 3.996293 | 0.699145 |
| MSTRG.18280.3 | 9.228234 | 4.664783 | 10.046487 |
| MSTRG.19236.5 | 4.336853 | 0.335709 | 0.812096 |
| MSTRG.19830.19 | 0 | 3.164567 | 0 |
| MSTRG.19893.1 | 5.376198 | 10.831803 | 3.382302 |
| MSTRG.19984.15 | 2.595783 | 2.114841 | 3.476071 |
| MSTRG.20588.5 | 1.483456 | 0.940748 | 3.534595 |
| MSTRG.20746.1 | 7.420726 | 0.061898 | 0 |
| MSTRG.2082.1 | 11.475153 | 2.364675 | 1.379082 |
| MSTRG.21619.5 | 3.21223 | 0 | 0.35815 |
| MSTRG.21785.2 | 1.913928 | 0.381042 | 1.464656 |
| MSTRG.21991.3 | 0 | 0 | 22.93528 |
| MSTRG.22884.5 | 4.038834 | 1.560007 | 2.47744 |
| MSTRG.22982.9 | 0 | 24.479645 | 4.291255 |
| MSTRG.22985.7 | 4.602664 | 3.164434 | 1.430563 |
| MSTRG.22985.8 | 1.865483 | 2.693162 | 2.447377 |
| MSTRG.23002.3 | 21.692251 | 0 | 0.081158 |
| MSTRG.23216.1 | 16.053537 | 13.498445 | 5.755608 |
| MSTRG.23216.3 | 0 | 0 | 9.47587 |
| MSTRG.23451.8 | 1.416395 | 0 | 0.356643 |
| MSTRG.23453.2 | 3.308153 | 0.079294 | 1.687035 |
| MSTRG.24152.3 | 3.511139 | 0.322671 | 6.877823 |
| MSTRG.24493.6 | 13.688178 | 0 | 1.200452 |
| MSTRG.24518.14 | 5.565189 | 3.69212 | 0.829496 |
| MSTRG.24993.2 | 64.74202 | 34.123161 | 0 |

| | | | |
|---|---|---|---|
| MSTRG.25190.12 | 4.682845 | 2.796372 | 2.562433 |
| MSTRG.2541.4 | 28.383059 | 1.782903 | 21.753717 |
| MSTRG.25756.10 | 0 | 9.384965 | 7.877532 |
| MSTRG.25864.9 | 1.252673 | 0.220394 | 0 |
| MSTRG.26294.6 | 0 | 12.327079 | 0 |
| MSTRG.26371.5 | 15.47363 | 20.237617 | 9.919929 |
| MSTRG.26902.1 | 0 | 0.29817 | 6.683096 |
| MSTRG.2705.4 | 3.797837 | 0.358736 | 1.893439 |
| MSTRG.28153.4 | 19.00839 | 21.662468 | 24.5245 |
| MSTRG.312.2 | 5.75435 | 2.839127 | 2.556122 |
| MSTRG.3126.4 | 1.278561 | 5.405648 | 1.698244 |
| MSTRG.4116.13 | 0 | 0.516936 | 0 |
| MSTRG.4394.4 | 1.118462 | 0.722967 | 3.814794 |
| MSTRG.44.10 | 1.137041 | 7.12675 | 0.754835 |
| MSTRG.478.6 | 11.803887 | 6.661332 | 6.376598 |
| MSTRG.5420.17 | 0 | 0.109581 | 1.437163 |
| MSTRG.5420.20 | 0 | 2.259726 | 3.02267 |
| MSTRG.596.9 | 2.752344 | 0 | 0 |
| MSTRG.5968.2 | 12.176775 | 22.558132 | 22.776337 |
| MSTRG.6073.5 | 9.815979 | 0 | 0.494394 |
| MSTRG.6133.1 | 6.159118 | 0.3453 | 0.534223 |
| MSTRG.6232.7 | 1.384922 | 0 | 1.772823 |
| MSTRG.6294.2 | 0 | 0.115624 | 0 |
| MSTRG.6863.13 | 5.063575 | 5.322104 | 5.704775 |
| MSTRG.6866.6 | 3.192162 | 6.542596 | 6.671259 |
| MSTRG.6866.7 | 4.549248 | 1.863417 | 4.219464 |
| MSTRG.6870.5 | 0 | 0 | 0 |
| MSTRG.7682.8 | 1.350969 | 3.301006 | 1.528005 |
| MSTRG.8274.5 | 3.061231 | 5.917725 | 8.218792 |
| MSTRG.8296.35 | 2.617182 | 0 | 0.792202 |
| MSTRG.8511.7 | 0.755462 | 1.40215 | 2.876319 |
| MSTRG.9308.4 | 0.960817 | 1.882173 | 2.013555 |
| MSTRG.9313.2 | 57.180172 | 0 | 0 |
| MSTRG.9377.10 | 0 | 0.223908 | 0.797317 |
| MSTRG.23868.9 | 0 | 0 | 5.132808 |

| **Table S6. GBM Analisys** | | |
|---|---|---|
| transcript_id | Gene host | Coding (?) |
| MSTRG.10029.4 | ENSG00000276045.3 | Coding |
| MSTRG.10044.22 | ENSG00000158023.10 | Coding |
| MSTRG.10056.23 | ENSG00000130779.20 | Coding |
| MSTRG.10080.15 | ENSG00000150967.18 | Coding |
| MSTRG.10082.11 | ENSG00000090975.12 | Non-Coding |
| MSTRG.10087.15 | ENSG00000111328.7 | Non-Coding |
| MSTRG.10087.16 | ENSG00000111328.7 | Non-Coding |
| MSTRG.10087.18 | ENSG00000111328.7 | Non-Coding |
| MSTRG.10087.1 | ENSG00000111328.7 | Non-Coding |
| MSTRG.10087.2 | ENSG00000111328.7 | Coding |
| MSTRG.10094.10 | ENSG00000111364.16 | Coding |
| MSTRG.10094.17 | ENSG00000111364.16 | Coding |
| MSTRG.10248.76 | ENSG00000196458.11 | Coding |
| MSTRG.10316.3 | ENSG00000150459.12 | Coding |
| MSTRG.10335.2 | ENSG00000180776.15 | Coding |
| MSTRG.10335.3 | ENSG00000180776.15 | Coding |
| MSTRG.10335.4 | ENSG00000180776.15 | Coding |
| MSTRG.10413.14 | ENSG00000122033.14 | Coding |
| MSTRG.1041.3 | ENSG00000066056.14 | Coding |
| MSTRG.10441.7 | ENSG00000102781.14 | Coding |
| MSTRG.10545.1 | ENSG00000183722.9 | Non-Coding |
| MSTRG.10545.2 | ENSG00000183722.9 | Non-Coding |
| MSTRG.10545.3 | ENSG00000183722.9 | Non-Coding |
| MSTRG.10573.14 | ENSG00000172766.19 | Coding |
| MSTRG.10573.7 | ENSG00000172766.19 | Coding |
| MSTRG.10663.8 | ENSG00000139684.14 | Coding |
| MSTRG.10672.8 | ENSG00000136146.15 | Coding |
| MSTRG.10672.9 | ENSG00000136146.15 | Coding |
| MSTRG.10682.1 | ENSG00000139679.15 | Non-Coding |
| MSTRG.1077.16 | ENSG00000142937.12 | Coding |
| MSTRG.10967.8 | ENSG00000102595.20 | Coding |
| MSTRG.1115.4 | ENSG00000117461.15 | Coding |

| | | |
|---|---|---|
| MSTRG.1115.4 | ENSG00000278139.1 | Coding |
| MSTRG.11171.8 | ENSG00000185989.11 | Coding |
| MSTRG.11200.3 | ENSG00000169062.15 | Coding |
| MSTRG.1123.12 | ENSG00000085999.13 | Coding |
| MSTRG.11504.23 | ENSG00000129493.15 | Coding |
| MSTRG.11509.6 | ENSG00000151413.17 | Coding |
| MSTRG.11524.17 | ENSG00000129521.14 | Non-Coding |
| MSTRG.11540.4 | ENSG00000198604.11 | Coding |
| MSTRG.11668.3 | ENSG00000213741.10 | Non-Coding |
| MSTRG.11697.21 | ENSG00000125375.15 | Coding |
| MSTRG.11699.34 | ENSG00000012983.11 | Coding |
| MSTRG.11699.35 | ENSG00000012983.11 | Coding |
| MSTRG.11741.6 | ENSG00000180998.12 | Coding |
| MSTRG.11777.6 | ENSG00000168175.15 | Coding |
| MSTRG.11778.4 | ENSG00000131981.16 | Coding |
| MSTRG.11802.26 | ENSG00000126777.18 | Coding |
| MSTRG.118.26 | ENSG00000248333.8 | Coding |
| MSTRG.1183.1 | ENSG00000186094.17 | Coding |
| MSTRG.1183.4 | ENSG00000186094.17 | Coding |
| MSTRG.11846.10 | ENSG00000100578.17 | Coding |
| MSTRG.12024.5 | ENSG00000100632.11 | Non-Coding |
| MSTRG.12087.10 | ENSG00000100767.16 | Coding |
| MSTRG.12087.12 | ENSG00000100767.16 | Non-Coding |
| MSTRG.12094.11 | ENSG00000184227.8 | Coding |
| MSTRG.12154.4 | ENSG00000119640.9 | Non-Coding |
| MSTRG.12285.6 | ENSG00000070778.13 | Coding |
| MSTRG.12463.7 | ENSG00000176473.13 | Coding |
| MSTRG.1248.1 | ENSG00000154222.14 | Coding |
| MSTRG.12563.12 | ENSG00000022976.15 | Coding |
| MSTRG.12563.13 | ENSG00000022976.15 | Coding |
| MSTRG.12858.5 | ENSG00000275835.5 | Coding |
| MSTRG.12999.12 | ENSG00000166912.17 | Coding |
| MSTRG.13035.6 | ENSG00000134153.10 | Coding |
| MSTRG.1304.3 | ENSG00000116205.14 | Non-Coding |
| MSTRG.1318.13 | ENSG00000184313.20 | Coding |

| | | |
|---|---|---|
| MSTRG.13201.29 | ENSG00000137804.14 | Coding |
| MSTRG.13201.29 | ENSG00000285920.2 | Coding |
| MSTRG.13201.30 | ENSG00000137804.14 | Coding |
| MSTRG.13201.30 | ENSG00000285920.2 | Coding |
| MSTRG.13201.33 | ENSG00000137804.14 | Coding |
| MSTRG.13201.33 | ENSG00000285920.2 | Coding |
| MSTRG.13201.4 | ENSG00000137804.14 | Coding |
| MSTRG.13201.4 | ENSG00000285920.2 | Coding |
| MSTRG.13201.7 | ENSG00000137804.14 | Coding |
| MSTRG.13201.7 | ENSG00000285920.2 | Coding |
| MSTRG.13248.15 | ENSG00000137842.7 | Coding |
| MSTRG.13339.5 | ENSG00000137872.16 | Coding |
| MSTRG.13471.1 | ENSG00000047346.12 | Coding |
| MSTRG.13490.24 | ENSG00000151575.14 | Non-Coding |
| MSTRG.13490.8 | ENSG00000151575.14 | Coding |
| MSTRG.13622.28 | ENSG00000140455.17 | Non-Coding |
| MSTRG.13622.29 | ENSG00000140455.17 | Non-Coding |
| MSTRG.13633.5 | ENSG00000140451.13 | Coding |
| MSTRG.13772.13 | ENSG00000066933.16 | Coding |
| MSTRG.13902.19 | ENSG00000140386.13 | Coding |
| MSTRG.13914.13 | ENSG00000173517.10 | Coding |
| MSTRG.13914.14 | ENSG00000173517.10 | Coding |
| MSTRG.13914.15 | ENSG00000173517.10 | Coding |
| MSTRG.13914.19 | ENSG00000173517.10 | Coding |
| MSTRG.13914.22 | ENSG00000173517.10 | Coding |
| MSTRG.13914.26 | ENSG00000173517.10 | Coding |
| MSTRG.13914.28 | ENSG00000173517.10 | Coding |
| MSTRG.13914.7 | ENSG00000173517.10 | Coding |
| MSTRG.14170.3 | ENSG00000140534.14 | Non-Coding |
| MSTRG.14232.13 | ENSG00000166965.12 | Non-Coding |
| MSTRG.14350.59 | ENSG00000068305.17 | Coding |
| MSTRG.14419.13 | ENSG00000103326.12 | Coding |
| MSTRG.14509.8 | ENSG00000180185.11 | Coding |
| MSTRG.14603.6 | ENSG00000008516.17 | Non-Coding |
| MSTRG.14604.11 | ENSG00000008517.16 | Coding |

| | | |
|---|---|---|
| MSTRG.14659.6 | ENSG00000217930.8 | Coding |
| MSTRG.14677.42 | ENSG00000103199.14 | Coding |
| MSTRG.14959.44 | ENSG00000185864.16 | Non-Coding |
| MSTRG.15009.11 | ENSG00000103365.15 | Coding |
| MSTRG.1502.1 | ENSG00000116729.14 | Coding |
| MSTRG.1502.2 | ENSG00000116729.14 | Coding |
| MSTRG.15062.33 | ENSG00000077238.14 | Coding |
| MSTRG.15088.12 | ENSG00000176476.9 | Coding |
| MSTRG.15151.12 | ENSG00000174943.11 | Coding |
| MSTRG.15231.19 | ENSG00000103510.20 | Non-Coding |
| MSTRG.15454.1 | ENSG00000087258.15 | Coding |
| MSTRG.15500.8 | ENSG00000102934.10 | Non-Coding |
| MSTRG.15529.21 | ENSG00000181938.13 | Coding |
| MSTRG.15535.1 | ENSG00000103034.14 | Coding |
| MSTRG.1554.13 | ENSG00000254685.6 | Coding |
| MSTRG.1554.14 | ENSG00000254685.6 | Coding |
| MSTRG.15633.7 | ENSG00000102974.16 | Coding |
| MSTRG.15723.37 | ENSG00000090857.13 | Coding |
| MSTRG.15746.1 | ENSG00000157368.10 | Non-Coding |
| MSTRG.15769.16 | ENSG00000166747.12 | Coding |
| MSTRG.15769.1 | ENSG00000166747.12 | Coding |
| MSTRG.15769.21 | ENSG00000166747.12 | Coding |
| MSTRG.15769.8 | ENSG00000166747.12 | Coding |
| MSTRG.15858.6 | ENSG00000166848.7 | Non-Coding |
| MSTRG.15931.2 | ENSG00000135697.10 | Non-Coding |
| MSTRG.16080.4 | ENSG00000225614.3 | Coding |
| MSTRG.16080.5 | ENSG00000225614.3 | Coding |
| MSTRG.16136.10 | ENSG00000075399.14 | Non-Coding |
| MSTRG.16159.11 | ENSG00000003249.13 | Non-Coding |
| MSTRG.16159.2 | ENSG00000003249.13 | Coding |
| MSTRG.16214.2 | ENSG00000070444.15 | Coding |
| MSTRG.16277.4 | ENSG00000167740.9 | Non-Coding |
| MSTRG.16338.4 | ENSG00000029725.17 | Coding |
| MSTRG.16402.5 | ENSG00000174282.12 | Coding |
| MSTRG.16446.3 | ENSG00000125434.11 | Non-Coding |

| | | |
|---|---|---|
| MSTRG.1652.5 | ENSG00000162643.13 | Coding |
| MSTRG.16570.15 | ENSG00000141027.21 | Coding |
| MSTRG.16629.2 | ENSG00000108557.19 | Non-Coding |
| MSTRG.16629.3 | ENSG00000108557.19 | Non-Coding |
| MSTRG.16629.4 | ENSG00000108557.19 | Non-Coding |
| MSTRG.16709.6 | ENSG00000142494.13 | Coding |
| MSTRG.16830.4 | ENSG00000004142.12 | Coding |
| MSTRG.16851.6 | ENSG00000167525.13 | Coding |
| MSTRG.16861.6 | ENSG00000173065.13 | Coding |
| MSTRG.16861.8 | ENSG00000173065.13 | Coding |
| MSTRG.16885.5 | ENSG00000160551.11 | Coding |
| MSTRG.16923.7 | ENSG00000126653.18 | Coding |
| MSTRG.16938.1 | ENSG00000176390.12 | Coding |
| MSTRG.16955.2 | ENSG00000176208.9 | Coding |
| MSTRG.16960.11 | ENSG00000184060.11 | Non-Coding |
| MSTRG.16960.72 | ENSG00000181481.14 | Non-Coding |
| MSTRG.17102.13 | ENSG00000278259.4 | Coding |
| MSTRG.17187.3 | ENSG00000108306.13 | Coding |
| MSTRG.17301.2 | ENSG00000108799.13 | Coding |
| MSTRG.17322.3 | ENSG00000108830.10 | Coding |
| MSTRG.17481.6 | ENSG00000259207.7 | Coding |
| MSTRG.17481.6 | ENSG00000259753.1 | Coding |
| MSTRG.17556.4 | ENSG00000064300.9 | Coding |
| MSTRG.17696.3 | ENSG00000181610.13 | Coding |
| MSTRG.17700.14 | ENSG00000180891.13 | Coding |
| MSTRG.17700.9 | ENSG00000180891.13 | Coding |
| MSTRG.17729.2 | ENSG00000108384.15 | Coding |
| MSTRG.17739.30 | ENSG00000068489.12 | Non-Coding |
| MSTRG.1780.16 | ENSG00000117500.13 | Coding |
| MSTRG.17820.7 | ENSG00000173838.12 | Coding |
| MSTRG.17855.4 | ENSG00000108592.17 | Coding |
| MSTRG.17930.25 | ENSG00000171634.18 | Coding |
| MSTRG.180.21 | ENSG00000158286.13 | Non-Coding |
| MSTRG.18058.19 | ENSG00000188612.12 | Non-Coding |
| MSTRG.18078.37 | ENSG00000266714.9 | Coding |

| | | |
|---|---|---|
| MSTRG.18088.17 | ENSG00000132478.10 | Non-Coding |
| MSTRG.18095.15 | ENSG00000188878.19 | Coding |
| MSTRG.18095.53 | ENSG00000188878.19 | Coding |
| MSTRG.18095.58 | ENSG00000188878.19 | Coding |
| MSTRG.18095.59 | ENSG00000188878.19 | Coding |
| MSTRG.18112.21 | ENSG00000161542.17 | Coding |
| MSTRG.18317.28 | ENSG00000176155.19 | Non-Coding |
| MSTRG.18317.56 | ENSG00000176155.19 | Non-Coding |
| MSTRG.18368.12 | ENSG00000175711.8 | Coding |
| MSTRG.18550.17 | ENSG00000134278.15 | Non-Coding |
| MSTRG.18557.12 | ENSG00000085415.16 | Coding |
| MSTRG.18636.7 | ENSG00000134508.12 | Coding |
| MSTRG.18673.1 | ENSG00000141447.18 | Coding |
| MSTRG.18673.2 | ENSG00000141447.18 | Coding |
| MSTRG.18762.6 | ENSG00000101746.15 | Coding |
| MSTRG.18782.10 | ENSG00000153391.15 | Coding |
| MSTRG.18784.15 | ENSG00000141429.13 | Coding |
| MSTRG.18871.17 | ENSG00000167216.17 | Coding |
| MSTRG.18935.49 | ENSG00000141646.13 | Coding |
| MSTRG.19029.8 | ENSG00000141664.10 | Coding |
| MSTRG.19046.8 | ENSG00000081913.14 | Coding |
| MSTRG.19291.16 | ENSG00000160953.16 | Non-Coding |
| MSTRG.19291.19 | ENSG00000160953.16 | Coding |
| MSTRG.19322.12 | ENSG00000133275.16 | Coding |
| MSTRG.19356.13 | ENSG00000176490.5 | Coding |
| MSTRG.19356.1 | ENSG00000176490.5 | Coding |
| MSTRG.19356.2 | ENSG00000176490.5 | Coding |
| MSTRG.19356.3 | ENSG00000176490.5 | Coding |
| MSTRG.19356.4 | ENSG00000176490.5 | Coding |
| MSTRG.19426.2 | ENSG00000008382.15 | Coding |
| MSTRG.19434.17 | ENSG00000167671.12 | Coding |
| MSTRG.19434.4 | ENSG00000167671.12 | Coding |
| MSTRG.19524.40 | ENSG00000198816.7 | Coding |
| MSTRG.19554.8 | ENSG00000104980.8 | Coding |
| MSTRG.19684.8 | ENSG00000196361.10 | Coding |

| | | |
|---|---|---|
| MSTRG.19691.2 | ENSG00000198551.10 | Non-Coding |
| MSTRG.19692.2 | ENSG00000102575.13 | Coding |
| MSTRG.19692.3 | ENSG00000102575.13 | Coding |
| MSTRG.19731.5 | ENSG00000104774.13 | Coding |
| MSTRG.19731.6 | ENSG00000104774.13 | Coding |
| MSTRG.19753.1 | ENSG00000105607.13 | Coding |
| MSTRG.19753.7 | ENSG00000105607.13 | Coding |
| MSTRG.19774.6 | ENSG00000141837.20 | Coding |
| MSTRG.19801.21 | ENSG00000072071.16 | Coding |
| MSTRG.19801.6 | ENSG00000072071.16 | Coding |
| MSTRG.19830.14 | ENSG00000160961.12 | Non-Coding |
| MSTRG.20076.6 | ENSG00000182141.10 | Coding |
| MSTRG.20144.12 | ENSG00000183850.14 | Coding |
| MSTRG.20231.3 | ENSG00000213965.3 | Non-Coding |
| MSTRG.20335.6 | ENSG00000167604.14 | Coding |
| MSTRG.20349.4 | ENSG00000196357.11 | Coding |
| MSTRG.20349.6 | ENSG00000196357.11 | Coding |
| MSTRG.20357.14 | ENSG00000186017.14 | Coding |
| MSTRG.20367.2 | ENSG00000267041.6 | Coding |
| MSTRG.2037.6 | ENSG00000116793.16 | Coding |
| MSTRG.2037.7 | ENSG00000116793.16 | Coding |
| MSTRG.2037.8 | ENSG00000116793.16 | Coding |
| MSTRG.20486.9 | ENSG00000160460.16 | Coding |
| MSTRG.20605.10 | ENSG00000178386.13 | Coding |
| MSTRG.20618.1 | ENSG00000159915.12 | Coding |
| MSTRG.20622.11 | ENSG00000267508.5 | Coding |
| MSTRG.20643.14 | ENSG00000142252.11 | Non-Coding |
| MSTRG.20643.6 | ENSG00000142252.11 | Non-Coding |
| MSTRG.20665.12 | ENSG00000177051.6 | Coding |
| MSTRG.20665.14 | ENSG00000177051.6 | Coding |
| MSTRG.20665.16 | ENSG00000177051.6 | Coding |
| MSTRG.20729.5 | ENSG00000160007.19 | Coding |
| MSTRG.20732.2 | ENSG00000130748.7 | Non-Coding |
| MSTRG.20741.21 | ENSG00000197405.8 | Coding |
| MSTRG.20741.40 | ENSG00000197405.8 | Coding |

| | | |
|---|---|---|
| MSTRG.20743.10 | ENSG00000134815.19 | Coding |
| MSTRG.20766.13 | ENSG00000185453.13 | Coding |
| MSTRG.20771.3 | ENSG00000178150.10 | Non-Coding |
| MSTRG.20791.7 | ENSG00000063176.16 | Coding |
| MSTRG.20800.3 | ENSG00000105538.10 | Coding |
| MSTRG.20848.39 | ENSG00000142552.8 | Coding |
| MSTRG.20976.15 | ENSG00000161551.14 | Coding |
| MSTRG.21005.8 | ENSG00000167562.12 | Coding |
| MSTRG.21005.8 | ENSG00000198482.13 | Coding |
| MSTRG.21023.12 | ENSG00000170954.11 | Coding |
| MSTRG.21023.7 | ENSG00000170954.11 | Coding |
| MSTRG.21305.65 | ENSG00000278129.2 | Coding |
| MSTRG.21305.65 | ENSG00000283515.1 | Coding |
| MSTRG.21436.16 | ENSG00000119185.12 | Coding |
| MSTRG.21436.59 | ENSG00000119185.12 | Coding |
| MSTRG.21472.12 | ENSG00000143870.12 | Coding |
| MSTRG.21543.3 | ENSG00000118965.14 | Coding |
| MSTRG.21587.26 | ENSG00000219626.9 | Non-Coding |
| MSTRG.21587.42 | ENSG00000219626.9 | Non-Coding |
| MSTRG.21587.44 | ENSG00000219626.9 | Coding |
| MSTRG.21624.7 | ENSG00000084733.11 | Coding |
| MSTRG.21662.25 | ENSG00000084774.14 | Coding |
| MSTRG.21666.41 | ENSG00000115207.13 | Coding |
| MSTRG.21744.33 | ENSG00000162959.13 | Coding |
| MSTRG.21798.15 | ENSG00000138061.12 | Coding |
| MSTRG.21810.18 | ENSG00000218739.10 | Non-Coding |
| MSTRG.21945.30 | ENSG00000068724.17 | Coding |
| MSTRG.21976.21 | ENSG00000243244.6 | Coding |
| MSTRG.2198.17 | ENSG00000266338.6 | Coding |
| MSTRG.2198.23 | ENSG00000266338.6 | Non-Coding |
| MSTRG.21996.5 | ENSG00000115239.22 | Coding |
| MSTRG.21996.6 | ENSG00000115239.22 | Coding |
| MSTRG.22002.11 | ENSG00000170634.13 | Non-Coding |
| MSTRG.22002.16 | ENSG00000170634.13 | Non-Coding |
| MSTRG.22002.17 | ENSG00000170634.13 | Coding |

| | | |
|---|---|---|
| MSTRG.22002.19 | ENSG00000170634.13 | Coding |
| MSTRG.22002.24 | ENSG00000170634.13 | Coding |
| MSTRG.22002.3 | ENSG00000170634.13 | Coding |
| MSTRG.22002.6 | ENSG00000170634.13 | Non-Coding |
| MSTRG.22002.8 | ENSG00000170634.13 | Non-Coding |
| MSTRG.22002.9 | ENSG00000170634.13 | Non-Coding |
| MSTRG.22034.8 | ENSG00000275052.5 | Coding |
| MSTRG.22080.14 | ENSG00000162929.14 | Coding |
| MSTRG.22218.9 | ENSG00000087338.5 | Coding |
| MSTRG.22232.13 | ENSG00000143977.14 | Non-Coding |
| MSTRG.22291.4 | ENSG00000144034.15 | Non-Coding |
| MSTRG.22419.4 | ENSG00000042493.16 | Coding |
| MSTRG.22433.2 | ENSG00000115523.16 | Non-Coding |
| MSTRG.22440.76 | ENSG00000115525.18 | Non-Coding |
| MSTRG.22440.77 | ENSG00000115525.18 | Non-Coding |
| MSTRG.22443.7 | ENSG00000132300.19 | Coding |
| MSTRG.22453.13 | ENSG00000115561.16 | Coding |
| MSTRG.22453.13 | ENSG00000249884.8 | Coding |
| MSTRG.2265.1 | ENSG00000131791.8 | Coding |
| MSTRG.22677.21 | ENSG00000185414.20 | Non-Coding |
| MSTRG.22692.1 | ENSG00000196460.14 | Non-Coding |
| MSTRG.22694.10 | ENSG00000170485.17 | Coding |
| MSTRG.22803.6 | ENSG00000169756.16 | Coding |
| MSTRG.22814.11 | ENSG00000186522.15 | Coding |
| MSTRG.22857.13 | ENSG00000144152.13 | Non-Coding |
| MSTRG.22857.17 | ENSG00000144152.13 | Non-Coding |
| MSTRG.22857.19 | ENSG00000144152.13 | Non-Coding |
| MSTRG.22857.1 | ENSG00000144152.13 | Non-Coding |
| MSTRG.22857.23 | ENSG00000144152.13 | Non-Coding |
| MSTRG.22857.27 | ENSG00000144152.13 | Non-Coding |
| MSTRG.22857.28 | ENSG00000144152.13 | Non-Coding |
| MSTRG.22857.3 | ENSG00000144152.13 | Non-Coding |
| MSTRG.2286.1 | ENSG00000188092.15 | Coding |
| MSTRG.23001.10 | ENSG00000136709.12 | Coding |
| MSTRG.23040.30 | ENSG00000072135.13 | Non-Coding |

| | | |
|---|---|---|
| MSTRG.23068.6 | ENSG00000152102.18 | Coding |
| MSTRG.23068.7 | ENSG00000152102.18 | Coding |
| MSTRG.23068.9 | ENSG00000152102.18 | Coding |
| MSTRG.23110.7 | ENSG00000150551.11 | Coding |
| MSTRG.23110.8 | ENSG00000150551.11 | Coding |
| MSTRG.2314.23 | ENSG00000269713.7 | Coding |
| MSTRG.2314.25 | ENSG00000269713.7 | Coding |
| MSTRG.2314.27 | ENSG00000269713.7 | Coding |
| MSTRG.2314.28 | ENSG00000269713.7 | Coding |
| MSTRG.2314.29 | ENSG00000269713.7 | Coding |
| MSTRG.2314.2 | ENSG00000269713.7 | Coding |
| MSTRG.2314.3 | ENSG00000269713.7 | Coding |
| MSTRG.2314.4 | ENSG00000269713.7 | Coding |
| MSTRG.23254.5 | ENSG00000115145.10 | Coding |
| MSTRG.23327.1 | ENSG00000241399.7 | Coding |
| MSTRG.23374.3 | ENSG00000169507.9 | Coding |
| MSTRG.23400.9 | ENSG00000172292.14 | Coding |
| MSTRG.23451.4 | ENSG00000115840.14 | Coding |
| MSTRG.23547.65 | ENSG00000116044.16 | Non-Coding |
| MSTRG.2359.17 | ENSG00000023902.14 | Coding |
| MSTRG.2364.13 | ENSG00000266472.5 | Non-Coding |
| MSTRG.23712.3 | ENSG00000196950.14 | Coding |
| MSTRG.23742.10 | ENSG00000162944.11 | Coding |
| MSTRG.23780.15 | ENSG00000155744.9 | Coding |
| MSTRG.23820.2 | ENSG00000182329.14 | Coding |
| MSTRG.23880.18 | ENSG00000204186.10 | Coding |
| MSTRG.2392.15 | ENSG00000197622.13 | Non-Coding |
| MSTRG.2392.15 | ENSG00000213190.3 | Non-Coding |
| MSTRG.2403.3 | ENSG00000143373.18 | Coding |
| MSTRG.24178.3 | ENSG00000115009.13 | Non-Coding |
| MSTRG.24178.5 | ENSG00000115009.13 | Coding |
| MSTRG.24178.6 | ENSG00000115009.13 | Coding |
| MSTRG.24178.7 | ENSG00000115009.13 | Coding |
| MSTRG.24408.4 | ENSG00000115677.17 | Coding |
| MSTRG.24413.3 | ENSG00000176720.6 | Coding |

| | | |
|---|---|---|
| MSTRG.24506.28 | ENSG00000132670.20 | Coding |
| MSTRG.24506.29 | ENSG00000132670.20 | Coding |
| MSTRG.2453.24 | ENSG00000182134.16 | Coding |
| MSTRG.24534.11 | ENSG00000088826.18 | Coding |
| MSTRG.24568.6 | ENSG00000125772.13 | Coding |
| MSTRG.24612.1 | ENSG00000149346.15 | Coding |
| MSTRG.24612.6 | ENSG00000149346.15 | Coding |
| MSTRG.24689.11 | ENSG00000089091.16 | Coding |
| MSTRG.24689.19 | ENSG00000089091.16 | Coding |
| MSTRG.24721.19 | ENSG00000089101.18 | Coding |
| MSTRG.24811.10 | ENSG00000130684.14 | Non-Coding |
| MSTRG.24891.6 | ENSG00000101350.8 | Coding |
| MSTRG.24911.5 | ENSG00000101391.21 | Coding |
| MSTRG.24919.14 | ENSG00000078699.21 | Coding |
| MSTRG.24919.19 | ENSG00000078699.21 | Coding |
| MSTRG.24919.24 | ENSG00000078699.21 | Coding |
| MSTRG.24919.6 | ENSG00000078699.21 | Coding |
| MSTRG.24919.7 | ENSG00000078699.21 | Coding |
| MSTRG.24938.10 | ENSG00000078747.15 | Coding |
| MSTRG.24938.16 | ENSG00000078747.15 | Coding |
| MSTRG.24938.28 | ENSG00000125971.16 | Non-Coding |
| MSTRG.24938.31 | ENSG00000125971.16 | Non-Coding |
| MSTRG.24938.6 | ENSG00000078747.15 | Coding |
| MSTRG.24992.143 | ENSG00000088367.23 | Coding |
| MSTRG.25019.6 | ENSG00000101363.12 | Non-Coding |
| MSTRG.25070.5 | ENSG00000124177.15 | Coding |
| MSTRG.25180.65 | ENSG00000101040.19 | Coding |
| MSTRG.25387.16 | ENSG00000101181.17 | Coding |
| MSTRG.25442.9 | ENSG00000101213.7 | Non-Coding |
| MSTRG.25453.3 | ENSG00000130584.11 | Coding |
| MSTRG.25457.52 | ENSG00000101152.11 | Coding |
| MSTRG.25457.57 | ENSG00000101152.11 | Coding |
| MSTRG.25457.58 | ENSG00000101152.11 | Coding |
| MSTRG.25457.67 | ENSG00000101152.11 | Coding |
| MSTRG.25682.3 | ENSG00000154723.12 | Coding |

| | | |
|---|---|---|
| MSTRG.25683.11 | ENSG00000154727.10 | Coding |
| MSTRG.256.9 | ENSG00000171621.14 | Coding |
| MSTRG.25722.26 | ENSG00000156273.16 | Non-Coding |
| MSTRG.25722.27 | ENSG00000156273.16 | Non-Coding |
| MSTRG.25722.28 | ENSG00000156273.16 | Non-Coding |
| MSTRG.25722.30 | ENSG00000156273.16 | Non-Coding |
| MSTRG.25722.31 | ENSG00000156273.16 | Non-Coding |
| MSTRG.25722.32 | ENSG00000156273.16 | Non-Coding |
| MSTRG.25722.33 | ENSG00000156273.16 | Non-Coding |
| MSTRG.25737.12 | ENSG00000156299.13 | Coding |
| MSTRG.25818.2 | ENSG00000185917.13 | Coding |
| MSTRG.25833.5 | ENSG00000159256.13 | Coding |
| MSTRG.25833.6 | ENSG00000159256.13 | Coding |
| MSTRG.25843.7 | ENSG00000159267.14 | Coding |
| MSTRG.25961.17 | ENSG00000160193.12 | Coding |
| MSTRG.26052.6 | ENSG00000160298.17 | Non-Coding |
| MSTRG.26140.5 | ENSG00000070413.20 | Coding |
| MSTRG.26456.17 | ENSG00000100095.19 | Coding |
| MSTRG.26575.1 | ENSG00000138942.16 | Coding |
| MSTRG.26575.4 | ENSG00000138942.16 | Non-Coding |
| MSTRG.26575.8 | ENSG00000138942.16 | Non-Coding |
| MSTRG.26594.11 | ENSG00000241878.11 | Coding |
| MSTRG.26724.20 | ENSG00000128346.11 | Non-Coding |
| MSTRG.26732.3 | ENSG00000128298.17 | Coding |
| MSTRG.26732.3 | ENSG00000184381.20 | Coding |
| MSTRG.26811.1 | ENSG00000100387.9 | Non-Coding |
| MSTRG.2682.3 | ENSG00000143256.5 | Coding |
| MSTRG.26847.13 | ENSG00000198911.12 | Coding |
| MSTRG.26852.17 | ENSG00000100167.20 | Coding |
| MSTRG.26873.3 | ENSG00000100227.18 | Coding |
| MSTRG.26884.2 | ENSG00000242247.11 | Coding |
| MSTRG.26884.4 | ENSG00000242247.11 | Coding |
| MSTRG.26884.9 | ENSG00000242247.11 | Coding |
| MSTRG.26950.3 | ENSG00000205643.11 | Non-Coding |
| MSTRG.26957.7 | ENSG00000075275.16 | Coding |

| | | |
|---|---|---|
| MSTRG.27055.2 | ENSG00000175928.6 | Coding |
| MSTRG.27105.7 | ENSG00000114026.21 | Coding |
| MSTRG.27105.7 | ENSG00000134072.11 | Coding |
| MSTRG.27108.47 | ENSG00000214021.16 | Coding |
| MSTRG.27202.7 | ENSG00000131381.12 | Coding |
| MSTRG.27259.25 | ENSG00000144566.11 | Non-Coding |
| MSTRG.27491.5 | ENSG00000008324.11 | Coding |
| MSTRG.27500.51 | ENSG00000181061.13 | Non-Coding |
| MSTRG.27500.52 | ENSG00000181061.13 | Non-Coding |
| MSTRG.27517.2 | ENSG00000160746.13 | Coding |
| MSTRG.27525.7 | ENSG00000185219.17 | Coding |
| MSTRG.27586.9 | ENSG00000088727.12 | Non-Coding |
| MSTRG.27589.12 | ENSG00000114648.12 | Coding |
| MSTRG.27621.20 | ENSG00000164053.21 | Coding |
| MSTRG.27634.2 | ENSG00000177479.19 | Coding |
| MSTRG.27663.8 | ENSG00000173540.12 | Coding |
| MSTRG.27722.3 | ENSG00000248487.9 | Non-Coding |
| MSTRG.27728.5 | ENSG00000164091.12 | Coding |
| MSTRG.27735.3 | ENSG00000010318.21 | Non-Coding |
| MSTRG.27735.5 | ENSG00000010318.21 | Non-Coding |
| MSTRG.27782.12 | ENSG00000157445.15 | Coding |
| MSTRG.27793.6 | ENSG00000180376.17 | Coding |
| MSTRG.27793.7 | ENSG00000180376.17 | Coding |
| MSTRG.27944.5 | ENSG00000144736.14 | Non-Coding |
| MSTRG.2812.24 | ENSG00000143147.14 | Coding |
| MSTRG.28182.11 | ENSG00000091972.18 | Coding |
| MSTRG.28182.1 | ENSG00000091972.18 | Coding |
| MSTRG.28182.3 | ENSG00000091972.18 | Coding |
| MSTRG.28264.4 | ENSG00000031081.11 | Coding |
| MSTRG.28278.7 | ENSG00000121577.13 | Non-Coding |
| MSTRG.28305.11 | ENSG00000153767.10 | Coding |
| MSTRG.28337.1 | ENSG00000138496.16 | Coding |
| MSTRG.28337.2 | ENSG00000138496.16 | Coding |
| MSTRG.28337.5 | ENSG00000138496.16 | Coding |
| MSTRG.28337.8 | ENSG00000138496.16 | Coding |

| | | |
|---|---|---|
| MSTRG.28427.1 | ENSG00000163902.12 | Non-Coding |
| MSTRG.28439.5 | ENSG00000132394.11 | Coding |
| MSTRG.28545.6 | ENSG00000129055.12 | Non-Coding |
| MSTRG.28746.5 | ENSG00000120742.11 | Non-Coding |
| MSTRG.28746.6 | ENSG00000120742.11 | Non-Coding |
| MSTRG.28746.7 | ENSG00000120742.11 | Non-Coding |
| MSTRG.28746.8 | ENSG00000120742.11 | Non-Coding |
| MSTRG.28747.11 | ENSG00000198843.13 | Non-Coding |
| MSTRG.28795.1 | ENSG00000114790.13 | Coding |
| MSTRG.2880.11 | ENSG00000117533.15 | Non-Coding |
| MSTRG.2906.1 | ENSG00000117592.9 | Coding |
| MSTRG.2907.18 | ENSG00000183831.7 | Coding |
| MSTRG.2907.18 | ENSG00000285777.1 | Coding |
| MSTRG.29094.7 | ENSG00000284862.3 | Coding |
| MSTRG.29132.8 | ENSG00000180834.7 | Non-Coding |
| MSTRG.29132.8 | ENSG00000283765.1 | Non-Coding |
| MSTRG.29179.2 | ENSG00000163900.11 | Non-Coding |
| MSTRG.29218.39 | ENSG00000073849.15 | Coding |
| MSTRG.29294.8 | ENSG00000187527.11 | Coding |
| MSTRG.29320.7 | ENSG00000114331.15 | Coding |
| MSTRG.29320.8 | ENSG00000114331.15 | Coding |
| MSTRG.29356.26 | ENSG00000213123.11 | Coding |
| MSTRG.29362.4 | ENSG00000163960.12 | Coding |
| MSTRG.29374.2 | ENSG00000180370.10 | Coding |
| MSTRG.29403.1 | ENSG00000250312.8 | Non-Coding |
| MSTRG.29403.2 | ENSG00000250312.8 | Coding |
| MSTRG.29403.3 | ENSG00000250312.8 | Coding |
| MSTRG.29403.5 | ENSG00000250312.8 | Non-Coding |
| MSTRG.29403.9 | ENSG00000250312.8 | Non-Coding |
| MSTRG.29419.4 | ENSG00000122068.13 | Coding |
| MSTRG.29436.11 | ENSG00000182903.16 | Coding |
| MSTRG.29436.23 | ENSG00000182903.16 | Coding |
| MSTRG.29436.25 | ENSG00000182903.16 | Coding |
| MSTRG.29436.35 | ENSG00000182903.16 | Coding |
| MSTRG.29436.36 | ENSG00000182903.16 | Coding |

| | | |
|---|---|---|
| MSTRG.29436.39 | ENSG00000182903.16 | Coding |
| MSTRG.29436.49 | ENSG00000182903.16 | Non-Coding |
| MSTRG.29441.1 | ENSG00000215375.6 | Non-Coding |
| MSTRG.29441.2 | ENSG00000215375.6 | Non-Coding |
| MSTRG.29485.8 | ENSG00000159733.13 | Coding |
| MSTRG.29490.79 | ENSG00000125386.15 | Coding |
| MSTRG.29803.5 | ENSG00000154274.15 | Coding |
| MSTRG.29833.11 | ENSG00000174125.8 | Coding |
| MSTRG.29833.1 | ENSG00000174125.8 | Coding |
| MSTRG.29833.27 | ENSG00000174130.12 | Coding |
| MSTRG.29833.2 | ENSG00000174125.8 | Coding |
| MSTRG.29833.9 | ENSG00000174125.8 | Coding |
| MSTRG.29878.4 | ENSG00000179299.17 | Coding |
| MSTRG.29932.2 | ENSG00000163288.14 | Coding |
| MSTRG.29932.5 | ENSG00000163288.14 | Coding |
| MSTRG.30086.6 | ENSG00000157426.14 | Coding |
| MSTRG.30351.1 | ENSG00000163319.11 | Non-Coding |
| MSTRG.30735.14 | ENSG00000164074.15 | Coding |
| MSTRG.30794.26 | ENSG00000109381.19 | Coding |
| MSTRG.30967.2 | ENSG00000164144.16 | Coding |
| MSTRG.31056.9 | ENSG00000052795.13 | Coding |
| MSTRG.31154.5 | ENSG00000109586.12 | Coding |
| MSTRG.31154.6 | ENSG00000109586.12 | Coding |
| MSTRG.31156.3 | ENSG00000164104.12 | Coding |
| MSTRG.31157.28 | ENSG00000164106.8 | Non-Coding |
| MSTRG.31157.33 | ENSG00000164106.8 | Non-Coding |
| MSTRG.31166.15 | ENSG00000164118.13 | Coding |
| MSTRG.31265.7 | ENSG00000109762.16 | Coding |
| MSTRG.3142.10 | ENSG00000151414.15 | Coding |
| MSTRG.31582.2 | ENSG00000113384.14 | Coding |
| MSTRG.31629.13 | ENSG00000082196.20 | Coding |
| MSTRG.31629.13 | ENSG00000273294.1 | Coding |
| MSTRG.31629.20 | ENSG00000273294.1 | Coding |
| MSTRG.31629.2 | ENSG00000273294.1 | Coding |
| MSTRG.31642.2 | ENSG00000113456.19 | Coding |

| | | |
|---|---|---|
| MSTRG.31656.16 | ENSG00000152620.13 | Coding |
| MSTRG.31725.7 | ENSG00000205765.9 | Coding |
| MSTRG.31734.3 | ENSG00000112972.15 | Coding |
| MSTRG.31734.4 | ENSG00000112972.15 | Coding |
| MSTRG.31762.3 | ENSG00000172244.9 | Coding |
| MSTRG.31790.2 | ENSG00000164171.11 | Coding |
| MSTRG.31854.15 | ENSG00000155542.12 | Coding |
| MSTRG.3206.17 | ENSG00000134369.15 | Coding |
| MSTRG.32098.1 | ENSG00000164347.18 | Coding |
| MSTRG.32323.10 | ENSG00000113391.19 | Coding |
| MSTRG.32323.11 | ENSG00000113391.19 | Coding |
| MSTRG.32323.12 | ENSG00000113391.19 | Coding |
| MSTRG.32323.13 | ENSG00000113391.19 | Coding |
| MSTRG.32323.14 | ENSG00000113391.19 | Coding |
| MSTRG.32323.15 | ENSG00000113391.19 | Coding |
| MSTRG.32363.43 | ENSG00000153113.23 | Coding |
| MSTRG.32447.12 | ENSG00000112893.9 | Coding |
| MSTRG.32447.20 | ENSG00000112893.9 | Coding |
| MSTRG.32447.21 | ENSG00000112893.9 | Coding |
| MSTRG.32572.1 | ENSG00000064652.11 | Coding |
| MSTRG.32619.2 | ENSG00000145794.17 | Coding |
| MSTRG.32669.22 | ENSG00000113522.14 | Coding |
| MSTRG.32712.32 | ENSG00000113575.10 | Coding |
| MSTRG.32712.37 | ENSG00000113575.10 | Coding |
| MSTRG.32734.15 | ENSG00000152705.8 | Coding |
| MSTRG.32803.11 | ENSG00000120733.14 | Coding |
| MSTRG.32803.12 | ENSG00000120733.14 | Coding |
| MSTRG.32969.6 | ENSG00000091009.8 | Coding |
| MSTRG.32982.59 | ENSG00000156475.18 | Non-Coding |
| MSTRG.32989.13 | ENSG00000169302.16 | Coding |
| MSTRG.33017.6 | ENSG00000132915.11 | Coding |
| MSTRG.33031.2 | ENSG00000183111.12 | Coding |
| MSTRG.33036.7 | ENSG00000113716.13 | Coding |
| MSTRG.33111.4 | ENSG00000145850.9 | Coding |
| MSTRG.33162.14 | ENSG00000170234.13 | Coding |

| | | |
|---|---|---|
| MSTRG.33162.1 | ENSG00000170234.13 | Coding |
| MSTRG.33162.7 | ENSG00000170234.13 | Coding |
| MSTRG.33162.8 | ENSG00000170234.13 | Coding |
| MSTRG.33318.1 | ENSG00000051596.9 | Coding |
| MSTRG.33327.16 | ENSG00000170085.18 | Coding |
| MSTRG.33385.2 | ENSG00000145912.8 | Non-Coding |
| MSTRG.33435.3 | ENSG00000131459.13 | Coding |
| MSTRG.33515.1 | ENSG00000170542.6 | Coding |
| MSTRG.33528.5 | ENSG00000137275.14 | Coding |
| MSTRG.33658.11 | ENSG00000124523.16 | Coding |
| MSTRG.33690.3 | ENSG00000230873.8 | Non-Coding |
| MSTRG.33933.28 | ENSG00000137338.5 | Non-Coding |
| MSTRG.34104.5 | ENSG00000204394.13 | Coding |
| MSTRG.34149.15 | ENSG00000168394.11 | Coding |
| MSTRG.34190.111 | ENSG00000197283.17 | Coding |
| MSTRG.34196.3 | ENSG00000137309.19 | Coding |
| MSTRG.34205.1 | ENSG00000196821.10 | Coding |
| MSTRG.34205.6 | ENSG00000196821.10 | Coding |
| MSTRG.34261.11 | ENSG00000198663.16 | Coding |
| MSTRG.34302.2 | ENSG00000112167.10 | Non-Coding |
| MSTRG.34335.30 | ENSG00000001167.14 | Coding |
| MSTRG.34377.36 | ENSG00000137161.17 | Coding |
| MSTRG.34382.9 | ENSG00000124541.7 | Coding |
| MSTRG.34389.14 | ENSG00000112659.14 | Coding |
| MSTRG.34506.15 | ENSG00000096093.16 | Coding |
| MSTRG.3454.15 | ENSG00000117697.15 | Coding |
| MSTRG.34559.1 | ENSG00000151917.18 | Non-Coding |
| MSTRG.34650.2 | ENSG00000119899.13 | Coding |
| MSTRG.34650.3 | ENSG00000119899.13 | Coding |
| MSTRG.34863.12 | ENSG00000279170.2 | Non-Coding |
| MSTRG.34864.16 | ENSG00000123552.17 | Coding |
| MSTRG.34911.1 | ENSG00000130347.13 | Coding |
| MSTRG.34911.2 | ENSG00000130347.13 | Coding |
| MSTRG.34918.3 | ENSG00000164494.12 | Coding |
| MSTRG.34918.4 | ENSG00000164494.12 | Coding |

| | | |
|---|---|---|
| MSTRG.34918.7 | ENSG00000164494.12 | Coding |
| MSTRG.34989.1 | ENSG00000155115.7 | Coding |
| MSTRG.35072.7 | ENSG00000047936.10 | Coding |
| MSTRG.35072.8 | ENSG00000047936.10 | Coding |
| MSTRG.35178.4 | ENSG00000164484.11 | Coding |
| MSTRG.35202.2 | ENSG00000112299.8 | Non-Coding |
| MSTRG.352.13 | ENSG00000048707.15 | Coding |
| MSTRG.35445.13 | ENSG00000131016.17 | Coding |
| MSTRG.35623.3 | ENSG00000112539.15 | Coding |
| MSTRG.35764.37 | ENSG00000122687.18 | Non-Coding |
| MSTRG.35764.37 | ENSG00000286192.1 | Non-Coding |
| MSTRG.35772.1 | ENSG00000106012.18 | Coding |
| MSTRG.35831.23 | ENSG00000122674.12 | Coding |
| MSTRG.35835.10 | ENSG00000106305.10 | Coding |
| MSTRG.35835.6 | ENSG00000106305.10 | Coding |
| MSTRG.35843.1 | ENSG00000164535.15 | Coding |
| MSTRG.35843.3 | ENSG00000164535.15 | Coding |
| MSTRG.35848.19 | ENSG00000146576.13 | Coding |
| MSTRG.35869.31 | ENSG00000106415.13 | Coding |
| MSTRG.3594.12 | ENSG00000143748.18 | Coding |
| MSTRG.36039.3 | ENSG00000188732.11 | Coding |
| MSTRG.36039.6 | ENSG00000188732.11 | Coding |
| MSTRG.36039.9 | ENSG00000188732.11 | Coding |
| MSTRG.36041.14 | ENSG00000196335.13 | Coding |
| MSTRG.3607.13 | ENSG00000154380.17 | Non-Coding |
| MSTRG.3607.19 | ENSG00000154380.17 | Coding |
| MSTRG.3607.21 | ENSG00000154380.17 | Coding |
| MSTRG.3607.23 | ENSG00000154380.17 | Coding |
| MSTRG.3607.24 | ENSG00000154380.17 | Coding |
| MSTRG.3607.2 | ENSG00000154380.17 | Non-Coding |
| MSTRG.3607.30 | ENSG00000154380.17 | Coding |
| MSTRG.3607.31 | ENSG00000154380.17 | Coding |
| MSTRG.3607.33 | ENSG00000154380.17 | Coding |
| MSTRG.3607.3 | ENSG00000154380.17 | Coding |
| MSTRG.3607.4 | ENSG00000154380.17 | Non-Coding |

| | | |
|---|---|---|
| MSTRG.36108.17 | ENSG00000106052.13 | Coding |
| MSTRG.36108.18 | ENSG00000106052.13 | Coding |
| MSTRG.36108.19 | ENSG00000106052.13 | Coding |
| MSTRG.36108.20 | ENSG00000106052.13 | Coding |
| MSTRG.36122.41 | ENSG00000106069.22 | Non-Coding |
| MSTRG.36122.41 | ENSG00000285162.1 | Non-Coding |
| MSTRG.36164.3 | ENSG00000154678.18 | Coding |
| MSTRG.36310.10 | ENSG00000175600.15 | Coding |
| MSTRG.36586.12 | ENSG00000146733.14 | Coding |
| MSTRG.36586.13 | ENSG00000146733.14 | Coding |
| MSTRG.36586.14 | ENSG00000146733.14 | Coding |
| MSTRG.36586.27 | ENSG00000146733.14 | Non-Coding |
| MSTRG.36586.7 | ENSG00000146733.14 | Coding |
| MSTRG.36595.1 | ENSG00000197008.9 | Coding |
| MSTRG.36635.43 | ENSG00000241258.7 | Non-Coding |
| MSTRG.36741.12 | ENSG00000106683.15 | Coding |
| MSTRG.36742.4 | ENSG00000086730.17 | Coding |
| MSTRG.36743.12 | ENSG00000049541.11 | Coding |
| MSTRG.36743.5 | ENSG00000049541.11 | Coding |
| MSTRG.36749.12 | ENSG00000263001.6 | Coding |
| MSTRG.36750.1 | ENSG00000263001.6 | Non-Coding |
| MSTRG.36757.10 | ENSG00000196275.14 | Coding |
| MSTRG.36797.2 | ENSG00000146700.9 | Non-Coding |
| MSTRG.36798.4 | ENSG00000188372.15 | Coding |
| MSTRG.36798.7 | ENSG00000188372.15 | Coding |
| MSTRG.36960.18 | ENSG00000004766.17 | Coding |
| MSTRG.37032.32 | ENSG00000196652.11 | Coding |
| MSTRG.37033.11 | ENSG00000196367.13 | Coding |
| MSTRG.37071.4 | ENSG00000185955.5 | Non-Coding |
| MSTRG.37092.6 | ENSG00000106351.13 | Coding |
| MSTRG.37092.7 | ENSG00000106351.13 | Coding |
| MSTRG.37097.11 | ENSG00000106327.13 | Coding |
| MSTRG.37097.19 | ENSG00000106327.13 | Coding |
| MSTRG.37097.2 | ENSG00000106327.13 | Coding |
| MSTRG.37110.5 | ENSG00000167011.9 | Coding |

| | | |
|---|---|---|
| MSTRG.37135.46 | ENSG00000160999.10 | Coding |
| MSTRG.37250.4 | ENSG00000135241.17 | Coding |
| MSTRG.37276.1 | ENSG00000164603.12 | Non-Coding |
| MSTRG.37415.29 | ENSG00000165055.15 | Coding |
| MSTRG.37514.2 | ENSG00000221866.9 | Coding |
| MSTRG.37514.3 | ENSG00000221866.9 | Coding |
| MSTRG.37531.2 | ENSG00000227471.9 | Coding |
| MSTRG.37544.9 | ENSG00000155561.15 | Coding |
| MSTRG.37582.2 | ENSG00000122779.18 | Coding |
| MSTRG.37582.3 | ENSG00000122779.18 | Coding |
| MSTRG.37582.9 | ENSG00000122779.18 | Coding |
| MSTRG.37783.2 | ENSG00000197362.15 | Coding |
| MSTRG.37795.23 | ENSG00000106479.11 | Coding |
| MSTRG.37824.8 | ENSG00000106565.18 | Coding |
| MSTRG.37825.1 | ENSG00000002933.9 | Coding |
| MSTRG.37858.1 | ENSG00000196584.3 | Non-Coding |
| MSTRG.37862.10 | ENSG00000133627.18 | Coding |
| MSTRG.37862.12 | ENSG00000133627.18 | Coding |
| MSTRG.37862.17 | ENSG00000133627.18 | Coding |
| MSTRG.37862.5 | ENSG00000133627.18 | Coding |
| MSTRG.37862.8 | ENSG00000133627.18 | Coding |
| MSTRG.38027.10 | ENSG00000155189.12 | Coding |
| MSTRG.3805.12 | ENSG00000054267.22 | Coding |
| MSTRG.3805.12 | ENSG00000188739.15 | Coding |
| MSTRG.3805.2 | ENSG00000054267.22 | Coding |
| MSTRG.3805.2 | ENSG00000188739.15 | Coding |
| MSTRG.3805.4 | ENSG00000054267.22 | Coding |
| MSTRG.3805.4 | ENSG00000188739.15 | Coding |
| MSTRG.3805.5 | ENSG00000054267.22 | Coding |
| MSTRG.3805.5 | ENSG00000188739.15 | Coding |
| MSTRG.3805.7 | ENSG00000054267.22 | Coding |
| MSTRG.3805.7 | ENSG00000188739.15 | Coding |
| MSTRG.3811.13 | ENSG00000162885.13 | Coding |
| MSTRG.38302.18 | ENSG00000147408.14 | Coding |
| MSTRG.38338.2 | ENSG00000104635.14 | Coding |

| | | |
|---|---|---|
| MSTRG.38338.3 | ENSG00000104635.14 | Coding |
| MSTRG.38338.8 | ENSG00000104635.14 | Coding |
| MSTRG.38340.7 | ENSG00000120910.14 | Coding |
| MSTRG.38359.12 | ENSG00000173535.14 | Non-Coding |
| MSTRG.38544.3 | ENSG00000187840.5 | Non-Coding |
| MSTRG.38565.31 | ENSG00000147548.17 | Coding |
| MSTRG.38565.33 | ENSG00000147548.17 | Coding |
| MSTRG.38570.11 | ENSG00000077782.20 | Coding |
| MSTRG.38630.15 | ENSG00000176209.11 | Non-Coding |
| MSTRG.38636.4 | ENSG00000168172.9 | Non-Coding |
| MSTRG.38733.10 | ENSG00000120992.18 | Coding |
| MSTRG.38770.12 | ENSG00000169122.11 | Coding |
| MSTRG.38781.2 | ENSG00000035681.9 | Non-Coding |
| MSTRG.38867.1 | ENSG00000165084.16 | Coding |
| MSTRG.3887.9 | ENSG00000203668.2 | Coding |
| MSTRG.3889.10 | ENSG00000174371.17 | Coding |
| MSTRG.3889.15 | ENSG00000174371.17 | Coding |
| MSTRG.3889.17 | ENSG00000174371.17 | Coding |
| MSTRG.3889.24 | ENSG00000174371.17 | Coding |
| MSTRG.3889.6 | ENSG00000174371.17 | Coding |
| MSTRG.3889.8 | ENSG00000174371.17 | Coding |
| MSTRG.39070.17 | ENSG00000133740.11 | Coding |
| MSTRG.39123.5 | ENSG00000253250.3 | Coding |
| MSTRG.39127.9 | ENSG00000155100.11 | Non-Coding |
| MSTRG.3915.10 | ENSG00000054282.16 | Coding |
| MSTRG.3915.11 | ENSG00000054282.16 | Coding |
| MSTRG.3915.12 | ENSG00000054282.16 | Coding |
| MSTRG.3915.14 | ENSG00000054282.16 | Coding |
| MSTRG.3915.18 | ENSG00000054282.16 | Coding |
| MSTRG.3915.1 | ENSG00000054282.16 | Coding |
| MSTRG.3915.23 | ENSG00000054282.16 | Coding |
| MSTRG.3915.26 | ENSG00000054282.16 | Coding |
| MSTRG.3915.3 | ENSG00000054282.16 | Coding |
| MSTRG.3915.5 | ENSG00000054282.16 | Non-Coding |
| MSTRG.3915.8 | ENSG00000054282.16 | Coding |

| | | |
|---|---|---|
| MSTRG.3918.15 | ENSG00000117020.18 | Coding |
| MSTRG.3930.25 | ENSG00000203666.12 | Non-Coding |
| MSTRG.3930.26 | ENSG00000203666.12 | Non-Coding |
| MSTRG.3940.2 | ENSG00000185420.19 | Coding |
| MSTRG.39416.17 | ENSG00000136982.6 | Coding |
| MSTRG.39416.4 | ENSG00000064313.12 | Non-Coding |
| MSTRG.39420.15 | ENSG00000187955.12 | Coding |
| MSTRG.39492.1 | ENSG00000147687.19 | Coding |
| MSTRG.39492.2 | ENSG00000147687.19 | Coding |
| MSTRG.39551.14 | ENSG00000153317.15 | Coding |
| MSTRG.39551.6 | ENSG00000153317.15 | Coding |
| MSTRG.39551.8 | ENSG00000153317.15 | Coding |
| MSTRG.39653.1 | ENSG00000130193.8 | Coding |
| MSTRG.39653.3 | ENSG00000130193.8 | Coding |
| MSTRG.39671.51 | ENSG00000185730.8 | Non-Coding |
| MSTRG.39740.2 | ENSG00000196378.11 | Coding |
| MSTRG.39740.4 | ENSG00000196378.11 | Coding |
| MSTRG.39740.7 | ENSG00000196378.11 | Coding |
| MSTRG.39740.8 | ENSG00000196378.11 | Coding |
| MSTRG.39780.39 | ENSG00000107104.18 | Coding |
| MSTRG.39780.40 | ENSG00000107104.18 | Coding |
| MSTRG.39819.12 | ENSG00000147853.17 | Coding |
| MSTRG.39824.1 | ENSG00000120158.12 | Coding |
| MSTRG.3984.1 | ENSG00000169224.13 | Coding |
| MSTRG.39907.9 | ENSG00000107186.16 | Coding |
| MSTRG.39994.6 | ENSG00000188352.12 | Non-Coding |
| MSTRG.40008.15 | ENSG00000099810.21 | Non-Coding |
| MSTRG.40008.15 | ENSG00000264545.2 | Non-Coding |
| MSTRG.40191.4 | ENSG00000159921.16 | Coding |
| MSTRG.40200.13 | ENSG00000165304.8 | Coding |
| MSTRG.40234.36 | ENSG00000122696.14 | Non-Coding |
| MSTRG.40234.37 | ENSG00000122696.14 | Non-Coding |
| MSTRG.40234.38 | ENSG00000122696.14 | Non-Coding |
| MSTRG.40234.39 | ENSG00000107338.10 | Coding |
| MSTRG.40234.39 | ENSG00000255872.3 | Coding |

| | | |
|---|---|---|
| MSTRG.40234.40 | ENSG00000122696.14 | Coding |
| MSTRG.40234.41 | ENSG00000122696.14 | Non-Coding |
| MSTRG.40234.42 | ENSG00000122696.14 | Non-Coding |
| MSTRG.40234.44 | ENSG00000122696.14 | Non-Coding |
| MSTRG.40238.25 | ENSG00000165275.10 | Non-Coding |
| MSTRG.40319.23 | ENSG00000154529.14 | Coding |
| MSTRG.403.1 | ENSG00000037637.11 | Coding |
| MSTRG.403.4 | ENSG00000037637.11 | Coding |
| MSTRG.403.5 | ENSG00000037637.11 | Coding |
| MSTRG.403.7 | ENSG00000037637.11 | Coding |
| MSTRG.40499.2 | ENSG00000107282.8 | Coding |
| MSTRG.40520.3 | ENSG00000107362.13 | Coding |
| MSTRG.40520.6 | ENSG00000107362.13 | Coding |
| MSTRG.40544.1 | ENSG00000135045.7 | Coding |
| MSTRG.40544.3 | ENSG00000135045.7 | Non-Coding |
| MSTRG.40612.1 | ENSG00000165118.15 | Coding |
| MSTRG.40626.3 | ENSG00000135049.15 | Coding |
| MSTRG.40672.10 | ENSG00000106723.17 | Coding |
| MSTRG.40672.16 | ENSG00000106723.17 | Coding |
| MSTRG.40672.29 | ENSG00000106723.17 | Coding |
| MSTRG.40672.5 | ENSG00000106723.17 | Coding |
| MSTRG.40735.5 | ENSG00000127080.10 | Coding |
| MSTRG.40752.17 | ENSG00000188938.17 | Non-Coding |
| MSTRG.40752.2 | ENSG00000188938.17 | Non-Coding |
| MSTRG.40752.6 | ENSG00000188938.17 | Non-Coding |
| MSTRG.40752.7 | ENSG00000188938.17 | Non-Coding |
| MSTRG.41161.23 | ENSG00000173611.17 | Non-Coding |
| MSTRG.41161.24 | ENSG00000173611.17 | Coding |
| MSTRG.41231.6 | ENSG00000167114.13 | Coding |
| MSTRG.41234.1 | ENSG00000167112.10 | Coding |
| MSTRG.41266.35 | ENSG00000171097.14 | Coding |
| MSTRG.41266.35 | ENSG00000286112.1 | Coding |
| MSTRG.41268.52 | ENSG00000095319.14 | Coding |
| MSTRG.41315.8 | ENSG00000107164.16 | Coding |
| MSTRG.41341.5 | ENSG00000125482.13 | Coding |

| | | |
|---|---|---|
| MSTRG.41341.6 | ENSG00000125482.13 | Coding |
| MSTRG.41437.21 | ENSG00000148396.18 | Coding |
| MSTRG.41487.24 | ENSG00000187609.16 | Coding |
| MSTRG.41487.27 | ENSG00000187609.16 | Coding |
| MSTRG.4164.8 | ENSG00000107537.14 | Coding |
| MSTRG.41655.32 | ENSG00000169239.13 | Non-Coding |
| MSTRG.41669.5 | ENSG00000169895.6 | Non-Coding |
| MSTRG.41671.5 | ENSG00000086712.13 | Coding |
| MSTRG.41866.1 | ENSG00000183690.13 | Coding |
| MSTRG.41891.1 | ENSG00000065923.10 | Coding |
| MSTRG.41928.15 | ENSG00000221994.10 | Coding |
| MSTRG.41967.14 | ENSG00000147144.13 | Coding |
| MSTRG.42050.4 | ENSG00000184083.12 | Non-Coding |
| MSTRG.42050.5 | ENSG00000184083.12 | Non-Coding |
| MSTRG.42050.6 | ENSG00000184083.12 | Coding |
| MSTRG.42058.8 | ENSG00000130119.16 | Coding |
| MSTRG.4207.3 | ENSG00000148468.17 | Coding |
| MSTRG.4239.10 | ENSG00000241058.4 | Non-Coding |
| MSTRG.42544.12 | ENSG00000101901.12 | Coding |
| MSTRG.42560.1 | ENSG00000123496.8 | Coding |
| MSTRG.42560.4 | ENSG00000123496.8 | Coding |
| MSTRG.42560.5 | ENSG00000123496.8 | Coding |
| MSTRG.42618.3 | ENSG00000125351.11 | Coding |
| MSTRG.42618.9 | ENSG00000125351.11 | Coding |
| MSTRG.42620.3 | ENSG00000125356.7 | Non-Coding |
| MSTRG.42620.4 | ENSG00000125356.7 | Non-Coding |
| MSTRG.42734.10 | ENSG00000134597.16 | Non-Coding |
| MSTRG.42832.11 | ENSG00000129675.16 | Coding |
| MSTRG.42832.14 | ENSG00000129675.16 | Coding |
| MSTRG.42832.17 | ENSG00000129675.16 | Coding |
| MSTRG.4290.5 | ENSG00000150867.14 | Coding |
| MSTRG.4290.7 | ENSG00000150867.14 | Coding |
| MSTRG.42997.20 | ENSG00000183479.12 | Coding |
| MSTRG.42997.20 | ENSG00000213397.10 | Coding |
| MSTRG.43010.31 | ENSG00000067829.19 | Coding |

| | | |
|---|---|---|
| MSTRG.43029.13 | ENSG00000013563.14 | Coding |
| MSTRG.43029.1 | ENSG00000013563.14 | Coding |
| MSTRG.43029.9 | ENSG00000013563.14 | Coding |
| MSTRG.43095.3 | ENSG00000067646.12 | Coding |
| MSTRG.4313.1 | ENSG00000151023.17 | Coding |
| MSTRG.4656.22 | ENSG00000099290.17 | Coding |
| MSTRG.4748.6 | ENSG00000183230.17 | Coding |
| MSTRG.477.19 | ENSG00000127472.11 | Non-Coding |
| MSTRG.4896.3 | ENSG00000107758.15 | Coding |
| MSTRG.4896.6 | ENSG00000107758.15 | Coding |
| MSTRG.491.3 | ENSG00000158828.8 | Coding |
| MSTRG.4922.4 | ENSG00000185009.12 | Coding |
| MSTRG.4935.8 | ENSG00000156671.15 | Coding |
| MSTRG.5074.8 | ENSG00000122376.11 | Coding |
| MSTRG.5084.4 | ENSG00000107789.16 | Coding |
| MSTRG.5199.15 | ENSG00000119969.15 | Coding |
| MSTRG.5260.20 | ENSG00000155229.21 | Coding |
| MSTRG.5344.10 | ENSG00000198408.14 | Coding |
| MSTRG.5344.12 | ENSG00000198408.14 | Coding |
| MSTRG.5344.19 | ENSG00000198408.14 | Coding |
| MSTRG.5344.20 | ENSG00000198408.14 | Coding |
| MSTRG.5354.24 | ENSG00000166197.16 | Coding |
| MSTRG.535.8 | ENSG00000133216.16 | Coding |
| MSTRG.5507.4 | ENSG00000107518.18 | Coding |
| MSTRG.5507.8 | ENSG00000107518.18 | Coding |
| MSTRG.5622.3 | ENSG00000019995.6 | Coding |
| MSTRG.5699.5 | ENSG00000068383.19 | Coding |
| MSTRG.5772.9 | ENSG00000023191.17 | Coding |
| MSTRG.6030.13 | ENSG00000130413.15 | Coding |
| MSTRG.6030.35 | ENSG00000130413.15 | Coding |
| MSTRG.6030.37 | ENSG00000130413.15 | Coding |
| MSTRG.6030.39 | ENSG00000130413.15 | Coding |
| MSTRG.6202.1 | ENSG00000151116.17 | Coding |
| MSTRG.625.13 | ENSG00000130695.15 | Coding |
| MSTRG.625.14 | ENSG00000130695.15 | Coding |

| | | |
|---|---|---|
| MSTRG.6462.1 | ENSG00000181830.8 | Coding |
| MSTRG.65.1 | ENSG00000187608.10 | Coding |
| MSTRG.6524.14 | ENSG00000149177.13 | Coding |
| MSTRG.65.3 | ENSG00000187608.10 | Non-Coding |
| MSTRG.688.4 | ENSG00000130768.15 | Coding |
| MSTRG.6889.45 | ENSG00000173653.8 | Coding |
| MSTRG.6952.5 | ENSG00000110090.13 | Coding |
| MSTRG.699.2 | ENSG00000120656.11 | Coding |
| MSTRG.709.7 | ENSG00000180098.9 | Coding |
| MSTRG.7124.8 | ENSG00000077514.9 | Non-Coding |
| MSTRG.7188.3 | ENSG00000074201.9 | Coding |
| MSTRG.7195.23 | ENSG00000087884.14 | Non-Coding |
| MSTRG.7306.29 | ENSG00000151376.16 | Coding |
| MSTRG.7377.32 | ENSG00000182919.14 | Coding |
| MSTRG.7377.35 | ENSG00000182919.14 | Coding |
| MSTRG.7377.37 | ENSG00000182919.14 | Coding |
| MSTRG.7377.38 | ENSG00000182919.14 | Coding |
| MSTRG.7377.3 | ENSG00000182919.14 | Coding |
| MSTRG.7422.4 | ENSG00000165895.19 | Coding |
| MSTRG.7570.16 | ENSG00000150764.14 | Coding |
| MSTRG.7570.18 | ENSG00000150768.15 | Coding |
| MSTRG.765.54 | ENSG00000121775.18 | Coding |
| MSTRG.765.63 | ENSG00000025800.14 | Coding |
| MSTRG.7659.19 | ENSG00000110274.16 | Coding |
| MSTRG.7659.20 | ENSG00000110274.16 | Coding |
| MSTRG.7659.23 | ENSG00000110274.16 | Coding |
| MSTRG.7699.16 | ENSG00000167283.8 | Non-Coding |
| MSTRG.7699.57 | ENSG00000118058.22 | Coding |
| MSTRG.7705.11 | ENSG00000095139.14 | Coding |
| MSTRG.7795.11 | ENSG00000023171.18 | Coding |
| MSTRG.7858.20 | ENSG00000150455.13 | Non-Coding |
| MSTRG.7890.7 | ENSG00000134909.18 | Coding |
| MSTRG.795.36 | ENSG00000116497.18 | Coding |
| MSTRG.795.38 | ENSG00000116497.18 | Coding |
| MSTRG.795.39 | ENSG00000116497.18 | Coding |

| | | |
|---|---|---|
| MSTRG.7964.17 | ENSG00000120647.10 | Coding |
| MSTRG.7986.5 | ENSG00000006831.10 | Coding |
| MSTRG.822.13 | ENSG00000121904.17 | Coding |
| MSTRG.8235.37 | ENSG00000134545.13 | Coding |
| MSTRG.8235.38 | ENSG00000255641.1 | Non-Coding |
| MSTRG.8235.38 | ENSG00000255819.7 | Non-Coding |
| MSTRG.8235.39 | ENSG00000255641.1 | Coding |
| MSTRG.8235.40 | ENSG00000255641.1 | Coding |
| MSTRG.8235.41 | ENSG00000183542.5 | Coding |
| MSTRG.8235.41 | ENSG00000255819.7 | Coding |
| MSTRG.8235.42 | ENSG00000183542.5 | Non-Coding |
| MSTRG.8235.42 | ENSG00000255819.7 | Non-Coding |
| MSTRG.8235.43 | ENSG00000134545.13 | Coding |
| MSTRG.8235.46 | ENSG00000255819.7 | Non-Coding |
| MSTRG.8479.8 | ENSG00000152944.9 | Coding |
| MSTRG.848.4 | ENSG00000134698.11 | Coding |
| MSTRG.8490.3 | ENSG00000087448.11 | Coding |
| MSTRG.850.8 | ENSG00000092847.12 | Coding |
| MSTRG.8600.4 | ENSG00000139131.13 | Coding |
| MSTRG.8919.19 | ENSG00000139631.18 | Coding |
| MSTRG.8924.10 | ENSG00000185591.10 | Coding |
| MSTRG.8924.9 | ENSG00000185591.10 | Coding |
| MSTRG.8983.3 | ENSG00000094916.16 | Non-Coding |
| MSTRG.9013.8 | ENSG00000170473.17 | Coding |
| MSTRG.9020.22 | ENSG00000111540.16 | Coding |
| MSTRG.9020.54 | ENSG00000123411.15 | Coding |
| MSTRG.9023.55 | ENSG00000196465.10 | Coding |
| MSTRG.9077.109 | ENSG00000179912.20 | Non-Coding |
| MSTRG.9077.110 | ENSG00000179912.20 | Non-Coding |
| MSTRG.9077.113 | ENSG00000179912.20 | Non-Coding |
| MSTRG.9077.114 | ENSG00000179912.20 | Non-Coding |
| MSTRG.9077.115 | ENSG00000179912.20 | Non-Coding |
| MSTRG.9077.117 | ENSG00000179912.20 | Non-Coding |
| MSTRG.9077.118 | ENSG00000179912.20 | Non-Coding |
| MSTRG.9077.120 | ENSG00000179912.20 | Non-Coding |

| | | |
|---|---|---|
| MSTRG.9083.3 | ENSG00000166986.15 | Coding |
| MSTRG.9086.1 | ENSG00000166987.15 | Non-Coding |
| MSTRG.9115.1 | ENSG00000135439.11 | Coding |
| MSTRG.9115.2 | ENSG00000135439.11 | Coding |
| MSTRG.9325.21 | ENSG00000111581.10 | Coding |
| MSTRG.9351.51 | ENSG00000166225.8 | Coding |
| MSTRG.9642.12 | ENSG00000120802.13 | Non-Coding |
| MSTRG.9670.10 | ENSG00000120800.5 | Coding |
| MSTRG.9721.4 | ENSG00000111696.12 | Coding |
| MSTRG.9857.15 | ENSG00000204842.17 | Coding |
| MSTRG.9857.2 | ENSG00000204842.17 | Coding |
| MSTRG.9857.5 | ENSG00000204842.17 | Coding |
| MSTRG.9930.13 | ENSG00000174989.13 | Coding |
| MSTRG.9930.18 | ENSG00000174989.13 | Non-Coding |
| MSTRG.9930.20 | ENSG00000174989.13 | Non-Coding |
| MSTRG.9930.2 | ENSG00000174989.13 | Coding |
| MSTRG.9930.7 | ENSG00000174989.13 | Coding |
| MSTRG.103.4 | ENSG00000116151.14 | Coding |
| MSTRG.103.5 | ENSG00000116151.14 | Coding |
| MSTRG.103.6 | ENSG00000116151.14 | Coding |
| MSTRG.103.7 | ENSG00000116151.14 | Coding |
| MSTRG.11846.10 | ENSG00000100578.17 | Coding |
| MSTRG.1264.11 | ENSG00000174348.13 | Non-Coding |
| MSTRG.1264.5 | ENSG00000174348.13 | Coding |
| MSTRG.1276.80 | ENSG00000121310.17 | Non-Coding |
| MSTRG.1276.85 | ENSG00000121310.17 | Non-Coding |
| MSTRG.1276.87 | ENSG00000121310.17 | Non-Coding |
| MSTRG.13772.13 | ENSG00000066933.16 | Coding |
| MSTRG.13976.17 | ENSG00000180953.11 | Non-Coding |
| MSTRG.13976.18 | ENSG00000180953.11 | Non-Coding |
| MSTRG.14976.2 | ENSG00000140740.11 | Coding |
| MSTRG.19603.24 | ENSG00000188321.13 | Coding |
| MSTRG.19603.55 | ENSG00000270011.7 | Coding |
| MSTRG.19919.10 | ENSG00000141971.13 | Non-Coding |
| MSTRG.19919.11 | ENSG00000141971.13 | Non-Coding |

| | | |
|---|---|---|
| MSTRG.19919.22 | ENSG00000141971.13 | Non-Coding |
| MSTRG.19919.9 | ENSG00000141971.13 | Non-Coding |
| MSTRG.21035.27 | ENSG00000160336.15 | Non-Coding |
| MSTRG.21035.27 | ENSG00000286261.1 | Non-Coding |
| MSTRG.21035.39 | ENSG00000160336.15 | Non-Coding |
| MSTRG.21035.39 | ENSG00000286261.1 | Non-Coding |
| MSTRG.21035.46 | ENSG00000160336.15 | Non-Coding |
| MSTRG.21035.46 | ENSG00000286261.1 | Non-Coding |
| MSTRG.22021.10 | ENSG00000162994.16 | Coding |
| MSTRG.22021.7 | ENSG00000162994.16 | Coding |
| MSTRG.24250.11 | ENSG00000144535.19 | Coding |
| MSTRG.24250.20 | ENSG00000144535.19 | Coding |
| MSTRG.24250.5 | ENSG00000144535.19 | Coding |
| MSTRG.24250.9 | ENSG00000144535.19 | Coding |
| MSTRG.24865.34 | ENSG00000101294.17 | Coding |
| MSTRG.24865.37 | ENSG00000101294.17 | Coding |
| MSTRG.24865.41 | ENSG00000101294.17 | Coding |
| MSTRG.24865.49 | ENSG00000101294.17 | Coding |
| MSTRG.24865.52 | ENSG00000101294.17 | Coding |
| MSTRG.25066.4 | ENSG00000124181.14 | Coding |
| MSTRG.26560.10 | ENSG00000167065.13 | Coding |
| MSTRG.26560.11 | ENSG00000167065.13 | Coding |
| MSTRG.26560.12 | ENSG00000167065.13 | Coding |
| MSTRG.26560.13 | ENSG00000167065.13 | Coding |
| MSTRG.26560.6 | ENSG00000167065.13 | Coding |
| MSTRG.26560.8 | ENSG00000167065.13 | Coding |
| MSTRG.26560.9 | ENSG00000167065.13 | Coding |
| MSTRG.27391.48 | ENSG00000093167.18 | Non-Coding |
| MSTRG.27406.18 | ENSG00000144668.12 | Non-Coding |
| MSTRG.27406.20 | ENSG00000144668.12 | Non-Coding |
| MSTRG.27613.1 | ENSG00000164048.14 | Coding |
| MSTRG.27613.2 | ENSG00000164048.14 | Coding |
| MSTRG.27613.4 | ENSG00000164048.14 | Coding |
| MSTRG.28203.12 | ENSG00000285943.1 | Non-Coding |
| MSTRG.30794.26 | ENSG00000109381.19 | Coding |

| | | |
|---|---|---|
| MSTRG.31280.12 | ENSG00000109794.13 | Non-Coding |
| MSTRG.31280.18 | ENSG00000109794.13 | Non-Coding |
| MSTRG.31280.25 | ENSG00000109794.13 | Non-Coding |
| MSTRG.31280.28 | ENSG00000109794.13 | Non-Coding |
| MSTRG.31280.29 | ENSG00000109794.13 | Non-Coding |
| MSTRG.31280.30 | ENSG00000109794.13 | Non-Coding |
| MSTRG.31280.31 | ENSG00000109794.13 | Coding |
| MSTRG.31280.3 | ENSG00000109794.13 | Non-Coding |
| MSTRG.31280.9 | ENSG00000109794.13 | Coding |
| MSTRG.31896.1 | ENSG00000049167.14 | Coding |
| MSTRG.31896.2 | ENSG00000049167.14 | Coding |
| MSTRG.32115.19 | ENSG00000113163.16 | Coding |
| MSTRG.32115.20 | ENSG00000113163.16 | Coding |
| MSTRG.32323.10 | ENSG00000113391.19 | Coding |
| MSTRG.32323.11 | ENSG00000113391.19 | Coding |
| MSTRG.32323.12 | ENSG00000113391.19 | Coding |
| MSTRG.32323.13 | ENSG00000113391.19 | Coding |
| MSTRG.32323.14 | ENSG00000113391.19 | Coding |
| MSTRG.32323.15 | ENSG00000113391.19 | Coding |
| MSTRG.33017.6 | ENSG00000132915.11 | Coding |
| MSTRG.33036.7 | ENSG00000113716.13 | Coding |
| MSTRG.3312.26 | ENSG00000162873.14 | Non-Coding |
| MSTRG.3312.5 | ENSG00000162873.14 | Non-Coding |
| MSTRG.34379.1 | ENSG00000124713.6 | Non-Coding |
| MSTRG.37078.24 | ENSG00000066923.17 | Non-Coding |
| MSTRG.37078.3 | ENSG00000066923.17 | Non-Coding |
| MSTRG.37079.19 | ENSG00000066923.17 | Non-Coding |
| MSTRG.37079.9 | ENSG00000066923.17 | Coding |
| MSTRG.38125.6 | ENSG00000104626.14 | Coding |
| MSTRG.39254.15 | ENSG00000104450.12 | Coding |
| MSTRG.40058.8 | ENSG00000235453.10 | Non-Coding |
| MSTRG.41166.3 | ENSG00000119414.11 | Coding |
| MSTRG.41166.4 | ENSG00000119414.11 | Coding |
| MSTRG.41231.6 | ENSG00000167114.13 | Coding |
| MSTRG.41866.4 | ENSG00000183690.13 | Coding |

| | | |
|---|---|---|
| MSTRG.7195.3 | ENSG00000048649.13 | Coding |
| MSTRG.8600.10 | ENSG00000139131.13 | Coding |
| MSTRG.8600.12 | ENSG00000139131.13 | Coding |
| MSTRG.8600.14 | ENSG00000139131.13 | Coding |
| MSTRG.8600.2 | ENSG00000139131.13 | Coding |
| MSTRG.8600.3 | ENSG00000139131.13 | Coding |
| MSTRG.8600.5 | ENSG00000139131.13 | Coding |
| MSTRG.8600.6 | ENSG00000139131.13 | Coding |
| MSTRG.8600.7 | ENSG00000139131.13 | Coding |
| MSTRG.8600.8 | ENSG00000139131.13 | Coding |
| MSTRG.8600.9 | ENSG00000139131.13 | Coding |
| MSTRG.9325.17 | ENSG00000111581.10 | Coding |
